# Cancer cell genetics shaping of the tumor microenvironment reveals myeloid cell-centric exploitable vulnerabilities in hepatocellular carcinoma

**DOI:** 10.1101/2023.10.29.564350

**Authors:** Christel FA Ramirez, Daniel Taranto, Masami Ando-Kuri, Marnix HP de Groot, Efi Tsouri, Zhijie Huang, Daniel de Groot, Roelof JC Kluin, Daan J Kloosterman, Joanne Verheij, Jing Xu, Serena Vegna, Leila Akkari

## Abstract

Myeloid cells are abundant and plastic immune cell subsets in the liver, to which pro-tumorigenic, inflammatory and immunosuppressive roles have been assigned in the course of tumorigenesis. Yet several aspects underlying their dynamic alterations in hepatocellular carcinoma (HCC) progression remain elusive, including the impact of distinct genetic mutations in shaping a cancer-permissive tumor microenvironment (TME). Here, we generated somatic HCC mouse models bearing clinically-relevant oncogenic driver combinations and subsequent pathway activation that faithfully recapitulated different human HCC subclasses. We identified cancer genetics’ specific and stage-dependent alterations of the liver TME associated with distinct histopathological and malignant HCC features. These models ranged from T cell-rich, more indolent HCC to aggressive tumors exhibiting heightened myeloid cell infiltration. Interestingly, MAPK-activated, *Nras*^G12D^-driven tumors presented a mixed phenotype of prominent inflammation and immunosuppression in a T cell-excluded TME, contrasting with *Nras*^G12V^ HCC, enriched in adaptive immune cells. Mechanistically, we identified a *Nras*^G12D^ cancer cell-driven, MEK-ERK1/2-SP1-dependent GM-CSF secretion enabling the accumulation of immunosuppressive and proinflammatory monocyte-derived Ly6C^low^ cells. GM-CSF blockade curbed the accumulation of this myeloid cell subset, reduced inflammation, induced cancer cell death and prolonged animal survival. Furthermore, the anti-tumor effect of GM-CSF neutralization synergized with the clinically-approved inhibition of the vascular endothelial growth factor (VEGF) to inhibit HCC outgrowth. These findings underscore the striking alterations of the myeloid TME consequential to MAPK pathway activation intensity and the potential of GM-CSF inhibition as a myeloid-centric therapy tailored to subsets of HCC patients.

## Introduction

Hepatocellular carcinoma, the most common form of primary liver cancer, is a highly heterogenous disease both at the pathological and molecular levels^1-3^. These tumors exhibit high intra- and inter-tumor heterogeneity, which challenges the development of effective therapies for advanced HCC patients^4^. As a consequence, HCC is a leading cause of cancer death worldwide^5^. Different environmental risk factors, such as viral hepatitis, metabolic syndromes or alcohol abuse contribute to the multifaceted HCC molecular pathogenesis^4^, fueled by the underlying chronic inflammatory background characterizing all etiologies^6^.

While pivotal to define precise molecular and immune HCC subclasses^4^, extensive genetic, epigenetic and transcriptomic analyses over the last decade have failed to advance novel precision medicine treatments targeting liver cancer cells^4^. Meanwhile, immune and stromal cell targeting strategies based on vascular endothelial growth factor (VEGF)-inhibition and program death-ligand 1 (PD-L1) blockade have shown unprecedent results^7-9^, being the first treatment regimen with improved overall survival relative to Sorafenib, a decade-long mainstay treatment for advanced-stage HCC patients^10,11^. However, the multifaceted heterogeneity of the tumor microenvironment (TME) remains a major challenge to immunotherapy efficacy, a therapeutic approach only benefitting a minority of HCC patients^12,13^.

Mounting evidence suggests that different cancer cell-intrinsic features, such as the genetic make-up and/or signaling pathway deregulation, play a critical role in shaping different TME^14,15^, thus emphasizing the need to implement tailored immunomodulation strategies according to cancer molecular profiles. In HCCs and other solid tumor types, well-established oncogenic signals such as *MYC*^16-19^, Wnt/ β- Catenin^12,20,21^, Ras-MAPK-ERK pathway activation or loss of the tumor suppressor gene *TP53*^22-26^ distinctively reprogram the liver local and systemic environment through either promoting inflammation, immunosuppression or dampening anti-tumor immunity^27^. Importantly, quantitative differences in the Ras-MAPK-ERK signaling pathway activation state can exert unique biological consequences in response to extrinsic cues specific to different TME^28^. Yet, the non-cell autonomous effects and cellular mediators ensuing oncogenic pathways activation threshold that may dictate the HCC TME landscape remain to be elucidated.

It is well established that a dynamic interplay between cancer cells and the immune system impacts disease progression and response to therapy^15,21,29-34^, a process further complexified when considering the fine balance between immunotolerance and inflammatory wound healing response peculiar to the liver regenerative capacity. Myeloid cells constitute a heterogenous and dynamic population of functionally plastic innate immune cells, whose recruitment, differentiation and activation are influenced by cancer- and stromal-derived cues^6,35,36^. Reciprocally, tumor-educated myeloid cells promote cancer cell proliferation, immunosuppression and immune evasion^6,35,37-39^. For instance, macrophage and NK cell recruitment and pro-tumorigenic functions are influenced by p53 expression in hepatic stellate cells or transformed hepatocytes^25,40^, respectively. Moreover, such effects are reversible, as exemplified by p53 restoration in cancer cells, which curbed the myeloid-driven immunosuppressive TME and reestablished anti-tumor immune surveillance^41^. Hence, to apply personalized immunomodulation strategies overcoming the limited effects of T cell-centric immunotherapy, the mechanisms underlying HCC interpatient heterogeneity that impact myeloid cell subset composition and activation ought to be unraveled.

Here, we generated preclinical murine models of HCC mimicking the deregulation of oncogenic signaling pathways commonly altered in HCC patients^26,42- 46^ and recapitulating distinct molecular and histopathological features characteristic of human HCC subclasses. Utilizing complementary multi-omics approaches and functional assays, we revealed the heterogeneity of the myeloid cell pool and identified dichotomic proinflammatory and immunosuppressive features fueling HCC pathogenesis in tumors presenting heightened MAPK activity. Mechanistically, a cancer cell-intrinsic MEK-ERK1/2-SP1 signaling cascade enforced expression and secretion of GM-CSF, an exploitable vulnerability synergistic with the current standard of care treatment for HCC employing VEGF neutralization. Overall, our results highlight the importance of inter-patient heterogeneity and stratification and reveal novel combinatorial immunomodulatory strategies as a potential therapeutic avenue for HCC patients.

## Results

### Genetically-distinct liver cancer mouse models display unique histopathological and molecular features

To address the intricate relationships between distinct oncogenic signaling pathway activation and immune cell shaping in hepatocarcinogenesis, we generated genetically-distinct, immunocompetent HCC mouse models using hydrodynamic tail vein-delivery of genetic elements^47-49^. These preclinical models comprised combinations of oncogene overexpression and tumor-suppressor gene knockout mimicking the most frequently altered signaling pathways in HCC patients (**Fig. 1A**). *Myc*, amplified in up to 30% of HCC patients^26,43^, was the oncogenic driver used in two of these HCC preclinical models, in combination with either loss of *Trp53* (mutated in up to 50% of HCCs^44^) or activation of AKT/PI3K/mTOR signaling pathway (activated in 50% of HCC patients^45^) through *Pten* deletion. Furthermore, we mimicked the activation of the mitogen-activated protein kinase (MAPK) signaling pathway (aberrantly activated in ∼50% of HCCs^46^) using constitutive expression of the proto-oncogene *RAS.* In light of the evident differences in MAPK activation states and cancer malignant features incumbered by distinct Ras isoforms and point mutations in several cancers^50-54^, including HCCs^28,41,55^, we employed distinct oncogenic mutations of *Nras*: *Nras*^G12D^ (*pT3-Nras^G12D^*-GFP) and *Nras*^G12V^ (*pT/CaggsNras*^G12V^-IRES-Luc), to generate two additional HCC models, both combined with *Pten* loss.

**Figure 1.**
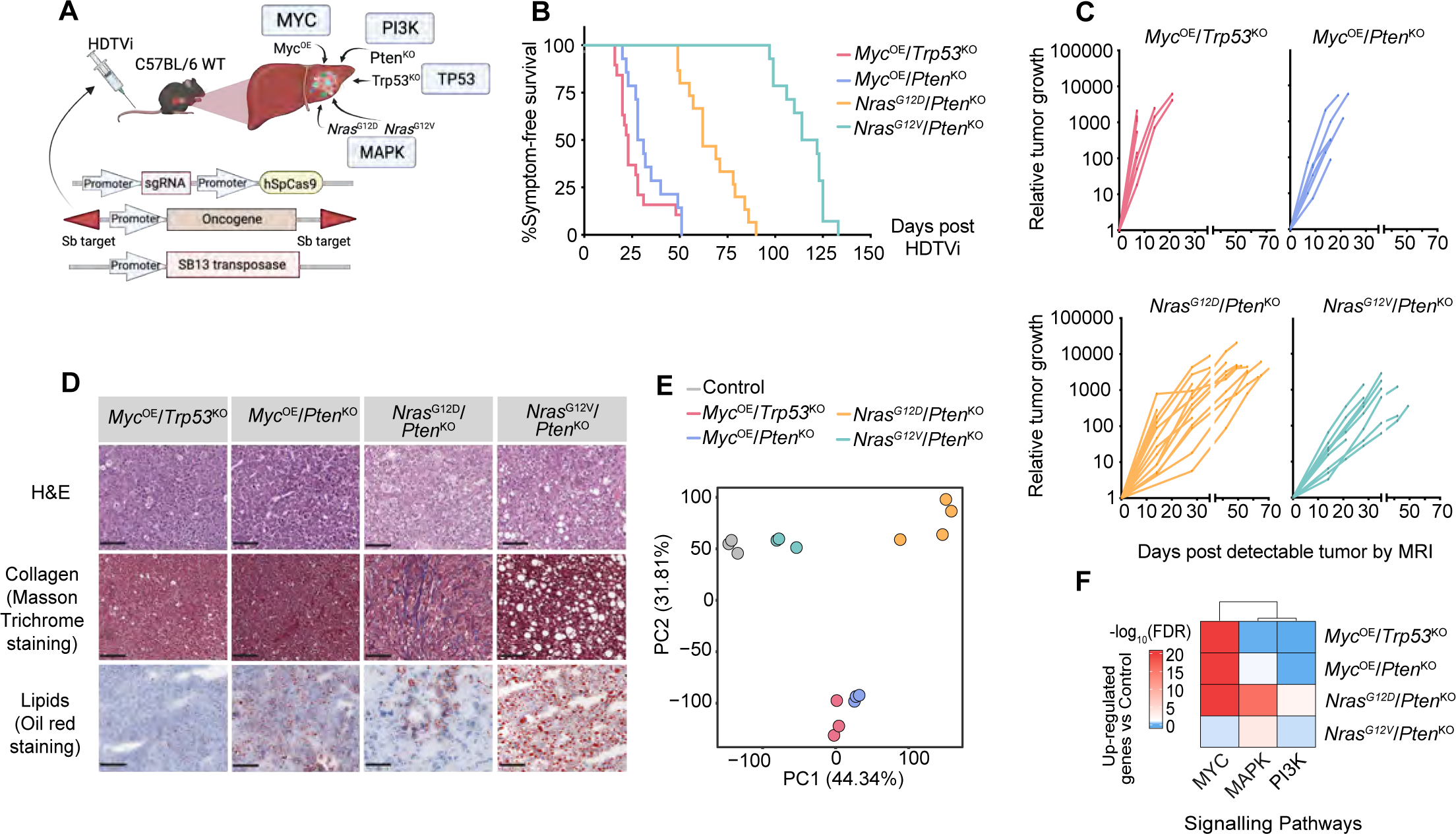
Genetically-distinct murine HCCs display distinct transcriptomic and histopathological features. **A.** Schematic diagram of experimental design: Hydrodynamic tail vein injection (HDTVi) method used to supply the Sleeping beauty (SB) transposon and CrispR- Cas9 constructs that enforce expression of the depicted oncogenic drivers in hepatocytes in order to generate genetically-distinct HCCs. **B.** Kaplan–Meier survival curves of HCC-bearing mice HDTV injected with the indicated combinations of oncogenic drivers (*Myc*^OE^/*Trp53*^KO^ n=19; median survival of 23 day, *Myc*^OE^/*Pten*^KO^ n=14; median survival of 29,5 days, *Nras*^G12D^/*Pten*^KO^ n=15; median survival of 62 days, *Nras^G12V^*/*Pten*^KO^ n=14; median survival of 118 days). **C.** Longitudinal HCC volumes of individual tumor-bearing mice (*Myc*^OE^/*Trp53*^KO^ n=9, *Myc*^OE^/*Pten*^KO^ n=6, *Nras*^G12D^/*Pten*^KO^ n=15, *Nras^G12V^*/*Pten*^KO^ n=9) determined from weekly/bi-weekly magnetic resonance imaging (MRI), and represented relative to each animal initial tumor volume. **D.** Representative images of H&E, Masson Trichrome and Oil Red staining performed on sectioned livers collected from end-stage HCC-bearing mice. Scale bars=100 µm. **E.** Principal Component Analysis (PCA) plot depicting the transcriptome differences between genetically-distinct HCC bulk tumors (*Myc*^OE^/*Trp53*^KO^ n=3, *Myc*^OE^/*Pten*^KO^ n=3, *Nras*^G12D^/*Pten*^KO^ n=4, *Nras^G12V^*/*Pten*^KO^ n=3) and control livers injected with empty vectors, hereafter referred to as control (n=3), following RNA-seq analyses (see **Supplementary Table 1**). **F.** Heatmap of unsupervised hierarchical clustering depicting the geneset enrichment of the MAPK, PI3K and MYC signaling pathways in genetically-distinct HCCs compared to control. The color scale represents the significance of the enrichment in –log10(FDR). FDR False Discovery Rate.

We first confirmed the oncogenic driver-mediated activation/inhibition of these pathways in established *Myc*^OE^/*Trp53*^KO^, *Myc*^OE^/*Pten*^KO^, *Nras*^G12D^/*Pten*^KO^, and *Nras*^G12V^/*Pten*^KO^ liver tumors (**Fig. S1A**). Tumor development and disease progression were assessed in each model using longitudinal magnetic resonance imaging (MRI; **Fig. S1B**) and survival curves were generated for each models following HDTVi- mediated tumor induction, with varying tumor penetrance rates (**Fig. 1B-C and Fig. S1B**). Both *Myc*^OE^-driven liver cancer models displayed multinodular tumors with brisk tumor latency and rapid outgrowth, consequently leading to short median survivals. This effect was irrespective of the invalidated *Trp53* or *Pten* tumor suppressor genes, thus confirming the dominant role of MYC in cancer progression^26,56^. Contrastingly, HCCs driven by *Nras* overexpression combined with *Pten*^KO^ displayed longer median survival, with different *Nras* point mutations resulting in distinct tumor penetrance, growth rates and nodule numbers. Indeed, *Nras*^G12D^ point mutation gave rise single-nodular tumors with a shorter latency compared to *Nras*^G12V^ multifocal HCC, which presented the longest median survival of all four liver cancer models. To exclude the use of distinct *Nras* vector backbones as a factor influencing these murine model features, we generated an additional *Nras*^G12V^/*Pten*^KO^ HCC model by directly mutating the *pT3-Nras^G12D^*-GFP vector in codon 12 from D to V, thus engineering a pT3- *Nras*^G12V^/*Pten*^KO^ HCC mouse model. We verified that pT3-*Nras*^G12V^/*Pten*^KO^ HCC model recapitulated the features of *pT/CaggsNras*^G12V^-IRES-Luc *Nras*^G12V^/*Pten*^KO^ (**Fig. S1C**), confirming that the aggressivity of the distinct *Nras*-driven models was subsequent to the introduced point mutations in hepatocytes.

We next investigated the histopathological and systemic malignant features of the genetically-distinct liver cancers models. We first confirmed the hepatocytic origin of these tumors with the readout of Arginase-1 expression^57^ (**Fig. S1D-E**). Increased systemic alpha fetoprotein (AFP)^58^ (**Fig. S1F**) and bulk tumor *Afp* expression (**Fig. S1G**) validated all four models as bona fide HCCs. Additionally, accumulation of seric alanine amino transferase (ALT) activity was evident in *Myc*^OE^/*Trp53*^KO^, *Myc*^OE^/*Pten*^KO^, and *Nras*^G12V^/*Pten*^KO^ tumor-bearing mice (**Fig. S1H**), indicating increased liver damage in multinodular tumors^59^. Consistent with the role of PI3K/AKT activation in triggering features of fatty liver disease^60^, lipid accumulation was observed in *Pten*^KO^ tumors. However, overt fibrosis and steatosis characterized *Nras*^G12D^/*Pten*^KO^ and *Nras*^G12V^/*Pten*^KO^ tumors, respectively (**Fig. 1D and Fig. S1I**). Pathological assessment revealed that *Myc*^OE^-driven tumors were poorly differentiated HCCs, while *Nras*^G12D^/*Pten*^KO^ and *Nras*^G12V^/*Pten*^KO^ tumors were moderate and well-differentiated HCCs, respectively (**Fig. S1J**), altogether confirming the correlation between tumor grade and overall survival reported in the human setting^4,61,62^.

We next exposed the transcriptome profiles of genetically-distinct end-stage HCCs by performing RNA-seq on bulk tumors. Interestingly, both *Myc*^OE^-driven models shared similar transcriptional profiles, supporting the dominant influence of Myc as the main oncogenic driver in these models, whereas *Nras*^G12V^/*Pten*^KO^ transcriptional signature resembled empty vector-injected control livers, likely due to their more differentiated features (**Fig. 1E and Fig. S1J-K**). In contrast, *Nras*^G12D^/*Pten*^KO^ tumors clustered away from the other samples (**Fig. 1E**), indicative of its unique transcriptome profile. We further validated the states of the signaling pathways governed by the enforced oncogenic drivers (**Fig. 1F and Fig. S1L, Supplementary Table 1**). Expectedly, *Myc*^OE^ models exhibited increased activity in the MYC pathway, potentially overriding other more subtle changes related to *Pten* loss-induced PI3K signaling activation in *Myc*^OE^/*Pten*^KO^ tumors. MAPK activity was heightened in both *Nras*-overexpressing models, albeit to a stronger extent in *Nras*^G12D^-driven HCC (**Fig. 1F and Fig. S1A,L**), suggesting differential MAPK intensity downstream of distinct *Nras* point mutations. Altogether, these results highlight the distinctive contribution of oncogenic pathways in influencing HCC progression and features, thereby providing novel tools to study the impact of cancer cell-intrinsic signaling states on HCC multilayered complexity.

### Genetically-distinct murine HCC models correlate with distinct human HCC molecular subclasses and prognostic rates

The heterogeneity of human HCCs has recently been revisited to incorporate molecular subclasses^4^. In order to validate the relevance of our HCC models to the human pathology, we used the transcriptomic profiles of genetically-distinct tumors (**Fig. 1E**) to establish gene signatures specific to each HCC model (**Fig. S2A and Supplementary Table 1**). We next applied a class prediction algorithm^26,63,64^ to the normalized transcriptomes and assessed their resemblance to human HCCs. Each of the genetically-distinct tumors segregated into different human molecular subtypes that matched their survival and histological features (**Fig. 2A and Fig. S2B**). While *Myc*^OE^/*Trp53*^KO^, *Myc*^OE^/*Pten*^KO^, and *Nras*^G12D^/*Pten*^KO^ transcriptional profiles all correlated with the human HCC proliferation class, *Myc*^OE^-driven HCCs closely associated with the human iCluster 3 and S2 classification and *Nras*^G12D^-driven HCC fitted within the iCluster 1 and S1 subclasses. In sharp contrast, *Nras*^G12V^/*Pten*^KO^- driven tumors associated with the non-proliferation class of HCCs, which correlated with better patient survival, low proliferation signatures and tallied to the human S3 subclass of patients bearing hepatocyte-like, more indolent liver tumors^64^ (**Fig. 2A and Fig. S2B**). Altogether, these findings closely recapitulated the survival rates and histological features observed *in vivo* (**Fig. 1B-D),** with fast-growing, less differentiated *Myc*^OE^- and *Nras*^G12D^-driven models, while *Nras*^G12V^-driven tumors correspond to the well-differentiated human molecular classification exhibiting better prognosis (**Fig. 2A**). These findings underscore the effect of *Nras* point mutations in driving divergent HCC models, further singularizing *Nras*^G12D^-driven HCC as a distinct aggressive model compared to *Myc*^OE^-associated liver tumors.

**Figure 2:**
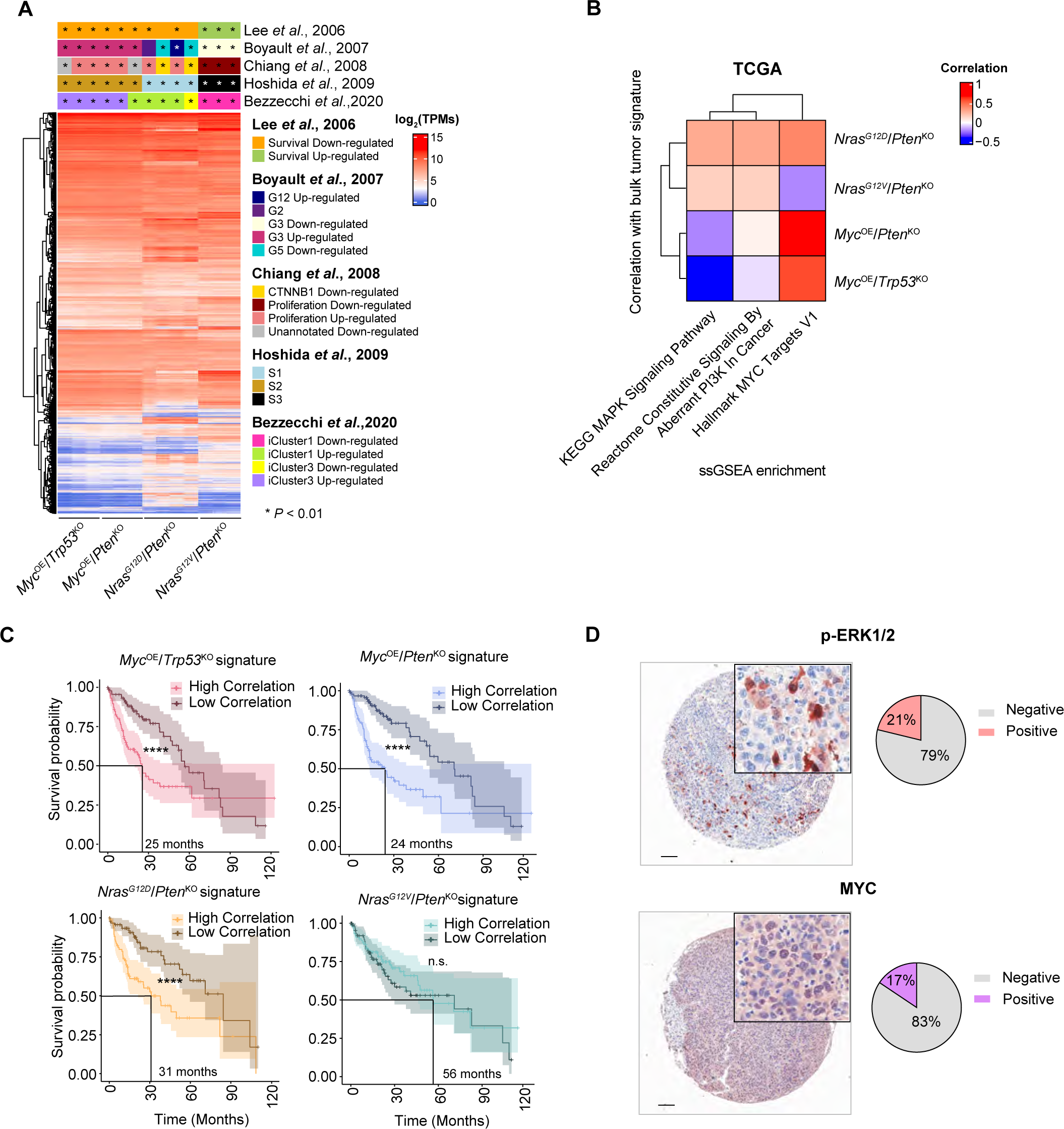
Transcriptional gene signatures derived from genetically-distinct HCC models recapitulate human HCC heterogeneity and predict patient prognosis. **A.** Heatmap of unsupervised hierarchical clustering depicting the global gene expression level as TPMs of genetically-distinct HCCs. Top annotations represent the classification of HCC mouse models’ transcriptomic profiles based on the human molecular HCC subtypes. **B.** Heatmap of unsupervised hierarchical clustering depicting the link between the MAPK, PI3K and MYC signaling pathway enrichment (**Fig. S2C**) and patients correlating to genetically-distinct HCC-derived transcriptional signatures across TCGA: Liver Hepatocellular Cancer (LIHC) patients (**Supplementary Table 2**). **C.** Kaplan-Meier survival curves displaying TCGA: LIHC patients segregated according to their high/low correlation with the transcriptional signatures of each genetically-distinct HCC relative to control. Lines at survival probability=0.5 depict median survival (see **Methods** for sample size and **Supplementary Table 2**). **D.** Representative IHC image for p-ERK1/2 and MYC performed on human HCC TMA sections. Pie charts depict the percentage of patients positive for p-ERK1/2 and Myc. Scale bars = 100 µm. Statistical significance was determined by unpaired Student’s t-test (**A**) and log-rank test (**C**). *p < 0.01, ****p < 0.0001; n.s. non-significant. TPMs Transcripts per Million.

We next sought to examine the enrichment profile of these oncogenic signaling pathways (**Fig. 1F**) across TCGA HCC patients and interrogated their correlation with murine HCC-derived signatures (**Fig. 2B and Fig. S2C**). These analyses asserted the dominance of *Myc*-driven transcriptional education in patients correlating with the *Myc*^OE^-derived murine signatures, and indicated that *Nras*^G12D^/*Pten*^KO^-correlating patients exhibited higher MAPK activation compared to their *Nras*^G12V^/*Pten*^KO^ counterpart, as observed in murine tumors. Moreover, the aggressive HCC models and human patients they associated with displayed matching overall survival outcome (**Fig. 2C and Supplementary Table 2;** see **Methods** for patient classification). The translational relevance of the murine HCC models was further supported by tumor mutation burden (TMB) (**Fig. S2D**) and mutational pattern (**Fig. S2E** and **Supplementary Table 3**) analyses from whole exome sequencing performed in each HCC murine model, which were overall comparable to Liver Hepatocellular Cancer (LIHC) patients.

To validate our findings in an independent cohort of HCC patients, we used a large tumor microarray dataset of 488 human HCC samples^65^ consisting of mostly poorly to moderately differentiated HCCs (**Supplementary Table 4**). Analyses of oncogenic signaling activity using immunohistochemistry staining revealed that MYC and MAPK/ERK were active in 17% and 21% of patients, respectively (**Fig. 2D**) and both correlated with poor patient prognosis (**Fig. S2F**). Altogether, our findings indicate that the activation state of oncogenic signaling pathway downstream of cancer cell mutations share similar prognostic rates in murine and human HCCs, both at the protein and transcriptome levels.

### Distinct oncogenic signaling pathways modulates the systemic and local tumor immune landscape

We next interrogated the impact of cancer cell genetic heterogeneity and related pathway activation on shaping the tumor immune landscape at the local and systemic levels during disease progression. Clear differences in the proportion of lymphoid and myeloid cell content were identified, with a significant dominance of myeloid cells in the more aggressive *Myc*^OE^/*Pten*^KO^, *Myc*^OE^/*Trp53*^KO^ and *Nras*^G12D^/*Pten*^KO^ HCCs (**Fig. 3A; Supplementary Data 1 and 2**). Longitudinal blood analyses of tumor-bearing mice suggested that this local increase partly originated from a systemic, tumor-induced myeloid cell response (**Fig. 3B**), with different myeloid cell subsets being mobilized in a cancer cell genetics-dependent manner (**Fig. S3A**). While systemic neutrophilia was observed in all four models to varying extents, the proportion of monocyte subsets increase – classical Ly6C^high^ versus the non-classical Ly6C^low^ monocytes - was specific to the myeloid-enriched HCC models (**Fig. S3A**).

**Figure 3:**
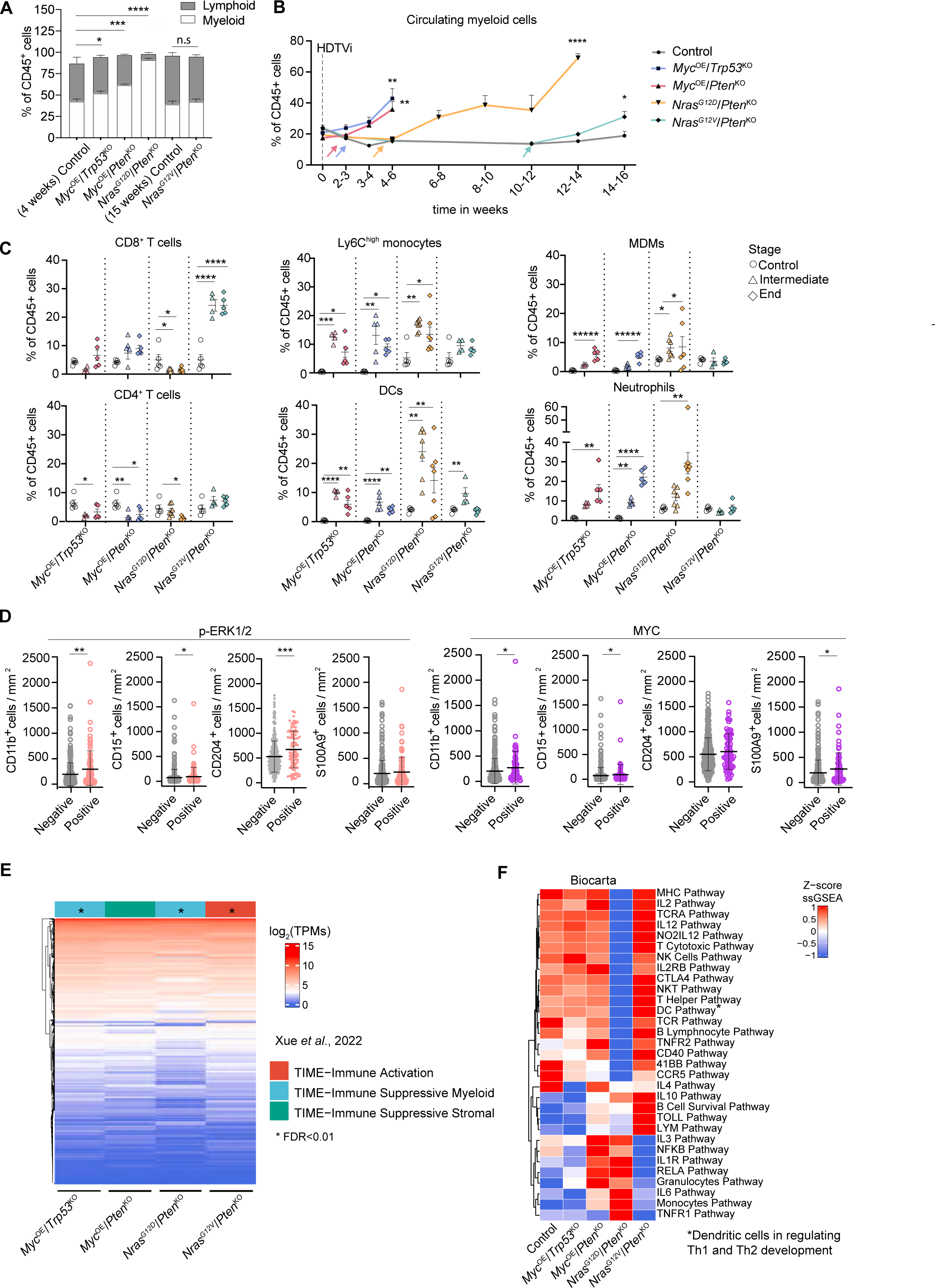
Activation of distinct oncogenic drivers differently shapes the HCC immune landscape. **A.** Quantification of the content of myeloid (CD45^+^ CD11b^+^) and lymphoid (CD45^+^ CD11b^-^) cells relative to total CD45^+^ leukocytes in tumors collected from end-stage genetically-distinct HCC-bearing mice and age-matched control livers (4 weeks and 15 weeks following HDTVi of empty vectors; Control-4-weeks n=5, *Myc*^OE^/*Trp53*^KO^ n=6, *Myc*^OE^/*Pten*^KO^ n=5, Control-15-weeks n=6, *Nras*^G12D^/*Pten*^KO^ n=7 and *Nras^G12V^*/*Pten*^KO^ n=4). **B.** Quantification of circulating myeloid cells (CD45^+^ CD11b^+^) in the blood relative to total CD45^+^ leukocytes at the indicated timepoints for each of the genetically-distinct HCC murine models (Control n=10, *Myc*^OE^/*Trp53*^KO^ n=20, *Myc*^OE^/*Pten*^KO^ n=19, *Nras*^G12D^/*Pten*^KO^ n=32, *Nras^G12V^*/*Pten*^KO^ n=18). Arrows indicate timepoints when detectable tumors were visible by MRI. **C.** Genetically-distinct HCC tumors were processed for flow cytometry at intermediate and end-stages (see **Methods**) to assess the content and activation profiles of the indicated immune cell populations. Lymphoid cells: CD8^+^ T cells (CD45^+^ CD11b^-^ NK1.1^-^ CD19^-^ CD3^+^ CD8^+^ CD4^-^), CD4^+^ T cells (CD45^+^ CD11b^-^ NK1.1^-^ CD19^-^ CD3^+^ CD8^-^ CD4^+^); myeloid cells: DCs (CD45^+^ F4/80^-^ CD11c^high^MHCII^high^) neutrophils (CD45^+^ CD11b^+^ Ly6C^int^ Ly6G^+^), Ly6C^high^ monocytes (CD45^+^ CD11b^+^ Ly6C^high^ Ly6G^-^), Ly6C^low^ monocytes (CD45^+^ CD11b^+^ Ly6C^low^ Ly6G^-^), monocyte-derived macrophages (MDMs; CD45^+^ CD11b^+^ Ly6C^low^ Ly6G^-^ F4-80^int^CD11b^high^). Graphs depicting the percentage of the aforementioned populations relative to total CD45^+^ leukocytes in control and genetically-distinct HCCs at intermediate and end-stage (Control-4-weeks n=5; *Myc*^OE^/*Trp53*^KO^ intermediate n=4 and end-stage n=5; *Myc*^OE^/*Pten*^KO^ intermediate n=5 and end-stage n=5, Control-15-weeks n=5, *Nras*^G12D^/*Pten*^KO^ intermediate n=7 and end-stage n=7; and *Nras^G12V^*/*Pten*^KO^ intermediate n=4 and end-stage n=5) (see **Supplementary Data 1** and **2** for gating strategy). **D.** Dotplots depicting the quantification of CD11B, CD15, CD204 and S100A9 positive cells by immunohistochemistry in paraffin-embedded HCC patient samples from the Wu et al. dataset^65^ segregated according to p-ERK1/2 (positive n=99, negative n=369) and c-MYC (positive n=78, negative n=389) positive staining in cancer cells (see **Methods**). **E.** Heatmap of unsupervised hierarchical clustering depicting the transcriptome of CD45^+^ cells isolated from genetically-distinct HCC (mean of n=3-5 per model) and their classification according to the human HCC immune subtypes clustering^71^. **F.** Heatmap of unsupervised hierarchical clustering depicting the ssGSEA enrichment scores per HCC model using immune-related pathways presented in the Biocarta database (see **Supplementary Table 6**). The color scale represents the z-score normalized enrichment per pathway (row) between HCC models. Graphs show mean + (**A, B**) or ± (**C**) SEM. Statistical significance was determined by unpaired Student’s *t*-test (**A-E**). Significance was determined within the myeloid content (**A).** *p < 0.05; **p < 0.01; ***p < 0.001; ****p < 0.0001.

Profiling of the TME at different stages of genetically-distinct HCC progression confirmed that immune cell composition and activation of lymphocytes and myeloid cells were tailored to the cancer cell mutational status at the bulk tumor level (**Fig. 3C and S3B**). Irrespective of disease stages, T cells were scarce in the more aggressive, myeloid-enriched *Myc*^OE^- and *Nras*^G12D^-driven tumors. Contrastingly, *Nras*^G12V^/*Pten*^KO^- driven TME displayed an overall increased content and activation of effector T cells, emphasizing the striking differences enforced by distinct *Nras* point mutations on the HCC TME. T cells were distinctively distributed in the HCC TME, with *Myc*^OE^-driven tumors poorly infiltrated by CD3^+^ cells, consistent with the immune-desert properties attributed to MYC proto-oncogene^21,56^ (**Fig. S3C**). *Nras*^G12V^/*Pten*^KO^ nodules displayed an “inflamed/ immunological hot” phenotype^66^, while *Nras*^G12D^/*Pten*^KO^ tumors presented CD3^+^ cells restricted to the peritumoral and tumor edge regions (**Fig. S3C**), a pattern referred to as “excluded/immunological cold”^66^, suggesting the establishment of a pro-tumorigenic, immunosuppressive environment. Increased infiltration of neutrophils and Ly6C^high^ monocytes was observed in *Myc*^OE^ and *Nras*^G12D^-driven tumors. These cells displayed low antigen presentation capacity and heightened immunosuppressive potential in all three aggressive models, albeit to a greater extent in *Nras*^G12D^-driven tumors, as evidenced by MHCII and PD-L1 expression, respectively (**Fig. S3B**). Moreover, *Myc*^OE^- and *Nras*^G12D^-driven tumors exhibited an increase in monocyte-derived macrophages (MDMs), with the latter showing the highest contribution of this population within the TME (**Fig. 3C and S3D**). This was in sharp contrast with *Nras*^G12V^-driven tumors, where tissue-resident Kupffer cells (KCs) still composed the largest proportion of hepatic macrophages (**Fig. S3D**). Furthermore, *Nras*^G12D^-driven tumors exhibited the highest expansion of PD-L1^+^ dendritic cells (DC) (**Fig. 3C and S3B**), thereby likely contributing to a tumor-permissive microenvironment. Collectively, these results reveal the accumulation of infiltrating myeloid cells with heightened immunosuppressive capacity within the *Nras*^G12D^-driven tumor milieu, thus setting this model apart from its *Nras*^G12V^-driven counterpart, but also from *Myc*^OE^-driven HCC.

Crucially, comparable TME landscape alterations were associated to specific oncogenic signaling pathway activation identified in human HCC (**Fig. 2D**). Indeed, activation of both MAPK/ERK and MYC pathways correlated with an increased CD11B^+^ myeloid cell presence and CD15^+^ neutrophil recruitment (**Fig. 3D**), corroborating our findings in *Myc*^OE^- and *Nras*^G12D^-driven murine models (**Fig. 3A, B**). Patients with heightened MAPK/ERK activity exhibited increased numbers of CD204^+^ positive cells, which identifies macrophages with pro-tumorigenic functions^67,68^. In contrast, MYC-overexpressing patients showed increased infiltration of S1009^+^ immunosuppressive myeloid cells^69,70^. Thus, these results provide further evidence of the distinct impact Myc and MAPK/ERK exert on myeloid cell education, which correlated with poor patient prognosis (**Fig. S2F**), thus asserting the translational relevance of our genetically-distinct murine models.

### Transcriptome profiling reveals a proinflammatory and tumor-permissive immune cell contexture within the *Nras*^G12D^/*Pten*^KO^ HCC TME

We next delved into the transcriptional heterogeneity of immune cells populating genetically-distinct HCC by performing RNA sequencing on FACS-isolated CD45^+^ cells (**Fig. S3E,F** and **Supplementary Table 5**). To unbiasedly characterized the TME phenotype across the distinct models, we classified their transcriptome profiles according to recently published liver tumor immune microenvironment (TIME) subtypes^71^. The *Nras*^G12V^/*Pten*^KO^ TME significantly classified in the TIME-IA (Immune Active) subtype (**Fig. 3E),** associated with significant T cell activation and infiltration, in line with our previous findings (**Fig. 3C and S3B**). Distinctively, *Myc*^OE^/*Trp53*^KO^ and *Nras*^G12D^/*Pten*^KO^ TME tallied to the TIME-SM (Suppressive Myeloid) subtype (**Fig. 3E**), associated with poor patient prognosis, myeloid cell dominance, and increased expression of both immunosuppressive and IL1-related inflammatory signaling pathways. Moreover, human HCC samples displaying high MAPK/ERK or Myc pathway activation were over-represented in patients exhibiting a stronger enrichment of the prognostic myeloid signature MRS (Myeloid Response Score), which correlates with immunosuppressive and pro-tumorigenic TME features (**Fig. S3G**)^65^. Altogether our findings indicate that Myc and MAPK/ERK pathway activation in cancer cells drives a tumor-permissive, myeloid-enriched TME in both preclinical and human HCC.

Next, the education profiles of CD45^+^ cells were further analyzed using immune clusters previously described^72^, as well as by comparing their transcriptome profiles to additional immune-related pathways (**Fig. 3F and Supplementary Table 6**). Gene sets enriched in *Myc*^OE^-driven models encompassed increased proliferation signature, Th2 cells and Interferon Gene clusters (**Fig. 3F and S3H**). Protein-protein interaction enrichment analyses identified the latter to be related to type I interferon signaling (**Fig. S3I**), often associated with innate immune cell response^73^. Conversely, lymphoid-rich, slow-growing *Nras*^G12V^-driven tumors exhibited a significant enrichment for T cell gene sets, Angiogenesis and the Immunologic Constant of Rejection (ICR) score, generally related to Th1 immunity^74^ (**Fig. S3H**), as well as increased IL-2, T cell activation, and co-stimulatory pathways (CD40, 41BB) (**Fig. 3F**), altogether confirming heightened adaptive immune response in these tumors. Substantiating the flow cytometry analyses and further singling out the unique TME profile of these tumors (**Fig. S3E,F**), *Nras*^G12D^/*Pten*^KO^ HCC displayed an enrichment of macrophages, immature DCs, TGF- signaling and Neutrophils pathway activity (**Fig. S3H**). In line with the immunosuppressive profile observed within the myeloid compartment (**Fig. 3E and S3B**), T cell-related signaling pathways associated with activation and function were down-regulated in *Nras*^G12D^/*Pten*^KO^ immune cells (**Fig. 3F).** Concomitantly, *Nras*^G12D^/*Pten*^KO^ displayed an enrichment in several proinflammatory pathways with well-described HCC pro-tumorigenic features^6^ also characteristic of the TIME-SM human subtype, such as NF-B, IL1R, IL6 and TNFR1 (**Fig. 3F**). As these pathways are known to be related to inflammasome activity^75,76^, we evaluated the levels of active caspase-1 in bulk tumors, which revealed a significantly increase in *Nras*^G12D^/*Pten*^KO^ HCC (**Fig. S3J**).

Altogether, these observations highlight the distinct HCC immune landscapes associated with different cancer cell molecular profiles. Despite a shared enrichment in myeloid cell content amongst the more aggressive tumor models, we consistently identified diverse education profiles between *Myc*^OE^-driven and *Nras*^G12D^/*Pten*^KO^ HCCs. Indeed, the latter exhibited a unique inflammatory background with prominent immunosuppressive capacity and dampened T cell activation features.

### *Nras*^G12D^*/Pten*^KO^-associated myeloid cells display a prominent pro-tumorigenic inflammatory signature at the single cell level

We further explored the heterotypic interactions underlying the unique *Nras*^G12D^-driven HCC TME by performing single cell RNA sequencing (scRNA-seq) on end-stage *Nras*^G12D^*/Pten*^KO^-driven tumor and age-matched control livers. Differential expression analysis identified seven unique clusters defined as immune cells based on *Ptprc* (CD45) expression (**Fig. 4A and S4A-C**) and then classified as myeloid or lymphoid cells according to their transcriptional profiles (**Fig. S4B-D and Supplementary Table 7**). We determined that both myeloid cell subsets - monocytic cells and neutrophils - were significantly enriched in *Nras*^G12D^*/Pten*^KO^ HCC, while lymphoid cells (T cell, NK cells and B cells), pDCs and cDC1s were decreased compared to control (**Fig. 4B and S4C**), emphasizing the major contribution of the myeloid compartment to the *Nras*^G12D^*/Pten*^KO^ TME at the single cell resolution.

**Figure 4:**
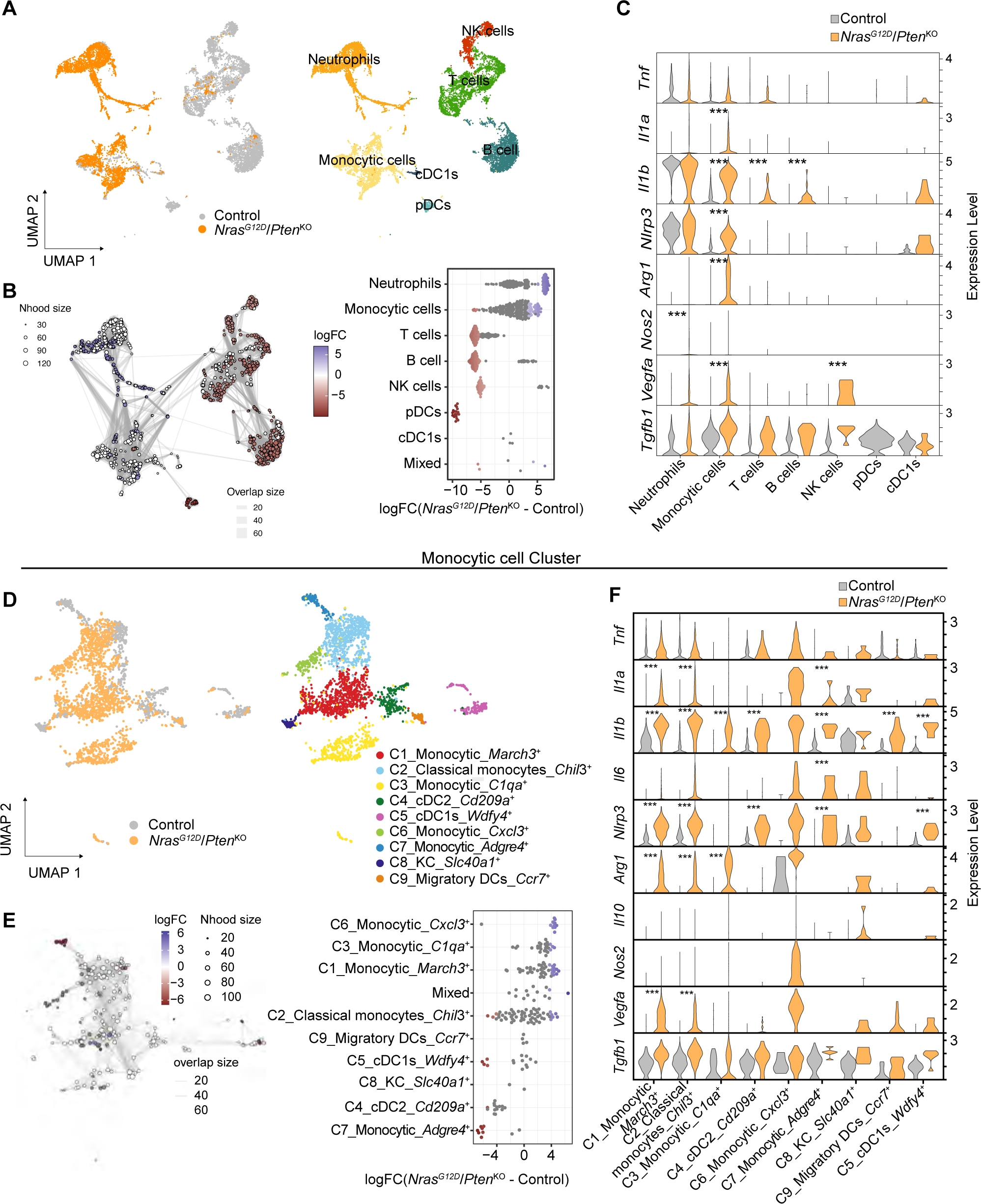
The *Nras^G12D^/Pten^KO^* TME displays transcriptomic heterogeneity with mixed pro-tumorigenic and inflammatory signatures. **A.** Uniform Manifold Approximation and Projection (UMAP) representation of CD45^+^ immune cells from control liver (n=7,447) and *Nras^G12D^*/*Pten*^KO^ HCC (n=4,814) (left) with annotated populations identified by scRNA-seq (right, see **Supplementary Table 7**). **B.** K-nearest neighbor (KNN) graph (left) and dotplot (right) depicting the differential abundance of cell types between *Nras^G12D^*/*Pten*^KO^ relative to control. Each dot represents a group of cells clustered in ‘neighborhoods’. Colors represent significant logFC (FDR ≤ 0.05), where white is non-significant difference in abundance. The thickness of the edges depicts the number of overlapping cells between neighborhoods. **C.** Violin plots depicting the normalized expression levels of inflammatory and immunosuppressive genes in the indicated immune cell subsets in control liver and *Nras*^G12D^/*Pten*^KO^ HCC (see **Supplementary Table 8**). **D.** UMAP representation of the ‘Monocytic cell’ subset (n=2,616) grouped with cDC1 (n=115) from (**A**). **E.** K-nearest neighbor (KNN) graph (left) and dotplot (right) depicting the differential abundance of myeloid cell subsets between *Nras^G12D^*/*Pten*^KO^ and control liver from cells grouped as ‘neighborhoods’ in (**D**). The colored dots represent significant changes in abundance using a threshold of FDR ≤ 0.05. **F.** Violin plots depicting the normalized expression levels of inflammatory and immunosuppressive genes in the indicated myeloid cell subsets in control livers and *Nras*^G12D^/*Pten*^KO^ HCC from the ‘Monocytic cell’ population (see **Supplementary Table 8**). Nhood=neighborhood. Statistical significance was determined by Wilcoxon Rank Sum test with multiple testing correction (**C, F).** *** FDR ≤ 0.001.

In light of the prominent inflammatory features observed in *Nras*^G12D^*/Pten*^KO^ CD45^+^ cells and bulk tumors (**Fig. 3F and S3J**), we next investigated the different immune-modulating functions altered within *Nras*^G12D^*/Pten*^KO^ TME cell subsets. Gene set analysis of *Nras*^G12D^*/Pten*^KO^ up-regulated genes showed significant enrichment for ‘NF- B signaling’ and ‘Inflammatory response’ in the ‘Monocytic cell’ cluster (**Fig. S4E and Supplementary Table 8**). Upon assessment of the genes included in these pathways, we observed a significant increase in proinflammatory (*Il1a, Il1b and Nlrp3*^75,77,78^) and immunosuppressive (*Nos2, Arg1 and Vegfa*^79-81^) gene expression (**Fig. 4C**).

We next explored the ‘Monocytic cells’ subset education profile, which we hypothesized likely contributed to the dual inflammatory and immunosuppressive phenotype of *Nras*^G12D^*/Pten*^KO^ HCC TME. Following the subsampling of this cluster, we identified nine subpopulations (**Fig. 4D**), from which the ‘C1_Monocytic_*March3*^+^’, ‘C2_Classical monocytes_*Chil3*^+^’, ‘C3_Monocytic_*C1qa*^+^’ and ‘C6_Monocytic_*Cxcl3*^+*’*^ were significantly more abundant in *Nras*^G12D^*/Pten*^KO^ compared to control (**Fig. 4E and S4F**). Unbiased cluster annotation using a publicly available human single cell RNAseq-dataset^71^ revealed that most monocytic subclusters exhibited macrophage, monocytes and DCs signatures (**Fig. S4G**). Differential expression (DE) analysis between control and *Nras*^G12D^*/Pten*^KO^ clusters within the ‘Monocytic cell’ population identified *Il1b* and *Nlrp3* up-regulation across all subpopulations (**Fig. 4F**), confirming the involvement of the inflammasome and IL-1 pathways in *Nras*^G12D^*/Pten*^KO^ tumorigenesis. Moreover, expression of immunosuppressive genes, such as *Arg1* and *Vegfa,* was increased in the ‘C1_Monocytic_*March3*^+^’, ‘C2_Classical monocytes_*Chil3*^+^’ and ‘C3_Monocytic_*C1qa*^+^ clusters (**Fig. 4F**). These findings were independently validated by RT-qPCR analyses comparing FACS-isolated MDMs and Ly6C^high^ monocytes from control livers and *Nras*^G12D^*/Pten*^KO^ HCC, which likely comprise the majority of cells identified within the ‘Monocytic cells’ subcluster (**Fig. S4H**). The significant increase in *Il1a*, *Il1b* and *Nlrp3* mRNA levels at different stages of *Nras*^G12D^*/Pten*^KO^ progression indicate that the IL-1 and inflammasome pathways are important attributes of myeloid cell education. Additionally, *Arg1* was also significantly increased in these myeloid cell populations (**Fig. S4G**), evoking their immunosuppressive phenotype, as previously reported in *Kras*^G12D^-driven PDAC tumors^82^. Collectively, these results highlight the heterogeneity of the *Nras*^G12D^- dictated TME, exposing a myeloid-rich immune landscape displaying mixed inflammatory and pro-tumorigenic features.

### GM-CSF signaling is uniquely activated in *Nras*^G12D^-driven tumors

We next sought to investigate the cancer cell-derived molecular players underlying *Nras*^G12D^/*Pten*^KO^ unique TME profile, and carried out RNA-seq of *Nras*^G12D^/*Pten*^KO^ and *Nras*^G12V^/*Pten*^KO^ HCC cell lines to complement bulk tumor analyses (**Fig. 5A and Supplementary Table 9A-B**). Expectedly, gene expression profiles of *Nras*^G12D^ and *Nras*^G12V^ cancer cells clustered away from each other (**Fig. S5A**), with changes largely conserved at the bulk tumor level (**Fig. 5A**). Gene set enrichment analyses of common DEG revealed that distinct lipid-associated pathways were significantly higher in *Nras*^G12V^ cancer cells (**Fig. S5B**), consistent with the steatotic features observed in *Nras*^G12V^/*Pten*^KO^ HCC (**Fig. 1D**). Processes related to inflammation, such as IL-2/STAT5, TNF-alpha signaling via NF-B and inflammatory response were significantly higher in *Nras*^G12D^*/Pten*^KO^ cancer cells and bulk tumors compared to their *Nras*^G12V^-driven counterparts (**Fig. 5B**). Altogether, these results suggest that cancer cell-intrinsic features shape several of the *Nras*-driven tumor characteristics, including the myeloid cell proinflammatory education identified in the CD45^+^ immune cells (**Fig. 3F**) and single-cell RNA-seq analyses (**Fig. 4**) of *Nras*^G12D^*/Pten*^KO^ HCC.

**Figure 5.**
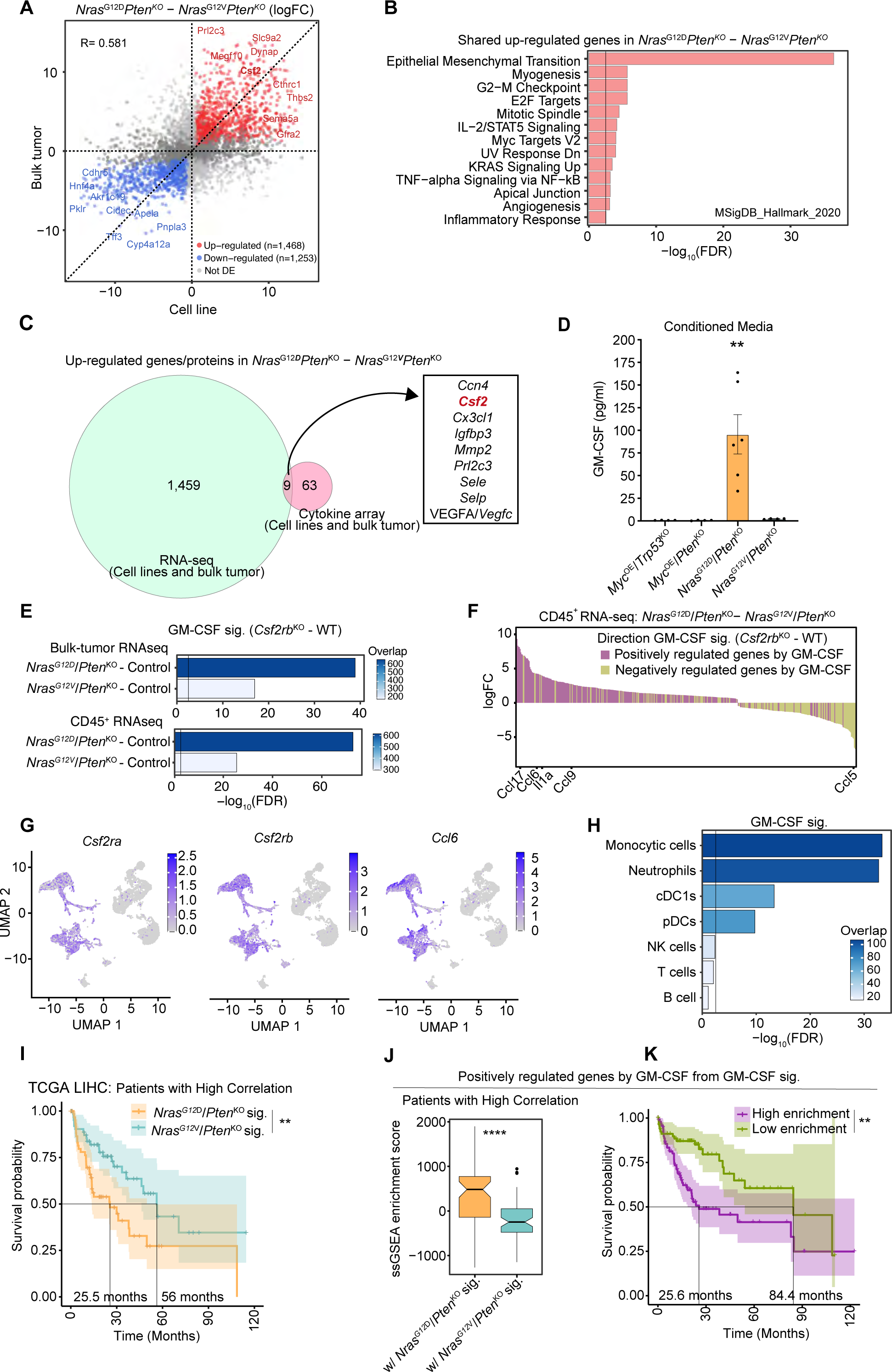
*Nras*^G12D^/*Pten*^KO^ mutated cancer cells specifically trigger GM-CSF signaling pathway in the TME. **A-C.** Differential expression analyses of RNA-seq datasets extracted from bulk tumors (presented in Fig. 1E) and cancer cell lines comparing *Nras^G12D^*/*Pten*^KO^ and *Nras^G12V^/Pten*^KO^ HCC and cancer cells (see **Supplementary Table 9A-B**). **A.** Scatterplot depicting the differential expressed genes between bulk tumors (y-axis) and cancer cell lines (x-axis). Colored dots represent genes significantly deregulated in both HCC bulk tumors and cancer cell lines. **B.** Barplot depicting the signaling pathways enriched in the up-regulated genes shown in (**A). C**. Venn Diagram depicting the overlap of up-regulated genes and proteins identified by RNA sequencing analyses presented in (**A**) and cytokine arrays (**Fig. S5C and D**), respectively. **D.** Barplot depicting the quantification of GM-CSF secretion in the conditioned media of genetically-distinct HCC cell lines (*Myc*^OE^/*Trp53*^KO^ n=4, *Myc*^OE^/*Pten*^KO^ n=4, *Nras*^G12D^/*Pten*^KO^ n=6, *Nras*^G12V^/*Pten*^KO^ n=6). **E.** Barplots depicting the enrichment of the GM-CSF signature^85^ (*Csf2rb*^KO^ *vs* WT, see **Methods**) in the DEG from *Nras*^G12D^/*Pten*^KO^ and *Nras*^G12V^/*Pten*^KO^ transcriptional signatures in bulk tumors (top) and CD45^+^ immune cells (bottom). Colors represent the overlap between the DEG from the *Nras-*driven HCC models and the GM-CSF signature. **F.** Barplot depicting the DEG from CD45^+^ immune cells RNA-seq of *Nras*^G12D^/*Pten*^KO^ relative to *Nras*^G12V^/*Pten*^KO^ that overlap with the GM-CSF signature. Colors represent the direction of GM-CSF regulation, specific genes are depicted (see **Supplementary Table 9C**). **G.** UMAP representations of *Csf2ra, Csf2rb* and *Ccl6* expression in the scRNA-seq from Fig. 4A. **H.** Barplot depicting the enrichment of the GM-CSF signature in each of the indicated immune cell subsets identified by scRNA-seq (presented in Fig. 4A). **I.** Kaplan-Meier survival curves displaying TCGA: LIHC patients segregated according to their high correlation with the *Nras*^G12D^/*Pten*^KO^ or *Nras*^G12V^/*Pten*^KO^ transcriptional signatures (presented in Fig. 2C) (*Nras*^G12D^/*Pten*^KO^ n=65, *Nras*^G12V^/*Pten*^KO^ n=66). **J.** Boxplot depicting the enrichment of GM-CSF positively regulated genes (n=628) from the GM-CSF signature in TCGA: LIHC patients segregated according to their high correlation with the *Nras*^G12D^/*Pten*^KO^ or *Nras*^G12V^/*Pten*^KO^ transcriptional signatures presented in Fig. 2C (*Nras*^G12D^/*Pten*^KO^ n=65, *Nras*^G12V^/*Pten*^KO^ n=66). **K.** Kaplan-Meier survival curves displaying TCGA: LIHC patients segregated according to their high/low enrichment of GM-CSF positively regulated genes (n=628) from the GM-CSF signature (High n=103, Low n=82). Graph show mean ± SEM (**D**). Statistical significance was determined by one-way ANOVA with Tukey’s multiple comparison test (**D**), log-rank test (**I, K**), unpaired Student’s *t*-test (**J**). Vertical lines at -log10(FDR)=2.5 are used as threshold for significance (**B, E, H**). **p < 0.01; ****p < 0.0001.

We next interrogated the distinctive TME features of all four HCC models at the protein level by performing cytokine profiling of end-stage bulk tumors and cancer cell lines (**Fig. S5C, D**). The secretome profiles of the two *Nras^OE^*-driven HCC models were then cross-checked with the up-regulated target genes identified by RNA sequencing. These unbiased analyses identified nine candidate factors up-regulated in *Nras*^G12D^*/Pten*^KO^ bulk tumors and cancer cells both at the transcriptome and proteome levels (**Fig. 5C**). Among these, *Csf2*/GM-CSF was of particular interest, with previous studies highlighting the increased levels of this cytokine in *Kras*^G12D^-driven PDAC associated with immature myeloid cell infiltration^82^. Indeed, GM-CSF is a master regulator of innate immune cells mediating their recruitment, survival and activation^83^. Crucially, GM-CSF can promote a proinflammatory response, for instance by directly inducing the expression and secretion of IL-1 cytokines^83-85^.

We thus hypothesized that the proinflammatory, myeloid-rich profile observed in the *Nras*^G12D^/*Pten*^KO^ TME may be driven by a cancer cell-intrinsic regulation of GM-CSF. Increased secretion of GM-CSF was independently confirmed in the supernatant of *Nras*^G12D^/*Pten*^KO^ cancer cells (**Fig. 5D and Fig. S5E**), and at the transcriptomic and protein levels (**Fig. S5F-I**). Importantly, by applying a publicly available GM-CSF signature^85^ (**Supplementary Table 9D**) to *Nras*^G12D^/*Pten*^KO^ bulk tumor and CD45^+^ RNA-seq datasets, we confirmed the significant enrichment of GM-CSF downstream pathways in this HCC model (**Fig. 5E**). Several cytokines and chemokines (e.g. *Ccl17*, *Cxcl14*, *Ccl24* and *Ccl6*) were up-regulated in CD45^+^ immune cells from *Nras*^G12D^/*Pten*^KO^ tumors compared to the *Nras*^G12V^/*Pten*^KO^ TME (**Fig. 5F and S5J**). Interestingly, GM-CSF increase was specific to the tumor bed, and no systemic upregulation was observed in the peripheral blood (**Fig. S5K**), suggesting a local re-education of the myeloid microenvironment rather than a systemic-elicited response.

We next queried the scRNA-seq dataset of *Nras*^G12D^/*Pten*^KO^ HCC (**Fig. 4**) to identify cell populations involved in GM-CSF signaling pathway. First, we validated that *Csf2* expression is virtually absent in stromal cells within the *Nras*^G12D^/*Pten*^KO^ TME (**Fig. S5L**), indicating that cancer cells are the major source of GM-CSF within the TME. Interestingly, while *Csf2ra*, *Csfr2b* and *Ccl6* were broadly expressed in myeloid cells (**Fig. 5G**), the GM-CSF gene expression signature was highly enriched in the two dominants ‘Monocytic cell’ and ‘Neutrophils’ myeloid cell clusters (**Fig. 5H and S5M,N**). These results suggest a pivotal role of cancer cell-derived GM-CSF signaling in shaping the immune landscape of *Nras*^G12D^-driven HCC.

Lastly, we investigated whether the *Nras*^G12D^/*Pten*^KO^*-*specific transcriptome and the GM-CSF signature may be of relevance in human HCC malignant features. First, we compared TCGA-LIHC patients presenting high correlation with either the *Nras*^G12D^/*Pten*^KO^ or *Nras*^G12V^/*Pten*^KO^ gene expression signatures (**Fig. 2C**) and determined that the former predicted poorer HCC patient survival (**Fig. 5I**). Patients with high *Nras*^G12D^/*Pten*^KO^ correlation scored significantly higher than *Nras*^G12V^/*Pten*^KO^- associated patients for GM-CSF signature enrichment (**Fig. 5J and S5O**). Furthermore, patients displaying high correlation with the GM-CSF signature exhibited a poorer prognosis (**Fig. 5K and S5P**), implying that *Nras*^G12D^/*Pten*^KO^ and GM-CSF transcriptional signatures are tightly connected in human HCCs and predict worse disease outcome. Overall, these results identified GM-CSF as a potential target in subgroups of HCC patients, while raising the questions of the underlying molecular mechanisms regulating GM-CSF secretion specifically in *Nras*^G12D^/*Pten*^KO^.

### Increased ERK1/2 activity drives GM-CSF secretion in a SP1-dependent manner and correlates with GM-CSF signature at the pan-cancer level

As GM-CSF is uniquely secreted by *Nras*^G12D^*/Pten*^KO^ cells (**Fig. 5D**), we sought to identify the molecular players downstream of Ras/MAPK and PI3K pathways involved in this specific regulation. We observed an increase in all phosphorylated MAPK-associated proteins in *Nras*^G12D^*/Pten*^KO^ compared to *Nras*^G12V^*/Pten*^KO^ cancer cells, including ERK1/2, AKT, S6RP, 4EBP1 and c- Jun (**Fig. 6A and S6A-B**), further validating the differential intensity of MAPK signal activity between the two *Nras*-driven HCC models. In order to identify which MAPK-associated proteins governed GM-CSF secretion in *Nras*^G12D^/*Pten*^KO^ cells, we used several inhibitors targeting RAS effector proteins, namely MEK (Trametinib), ERK1/2 (Temuterkib), ERK2 (Vx-11), mTOR (AZD8055), AKT (MK2066) and JNK (SP600125) (**Fig. S6C, D**). While inhibition of mTOR, AKT and JNK pathways did not affect GM-CSF level (**Fig. S6D**), MEK1/2, ERK1/2, and ERK2 inhibitors hampered *Nras*^G12D^*/Pten*^KO^-derived GM-CSF secretion (**Fig. 6B**) with no measurable effects on cancer cell proliferation (**Fig. S6E**). We further validated this finding by generating constitutive *Erk1* (sh*Erk1*) and *Erk2* (sh*Erk2*) knockdown cell lines (**Fig. S6F**) and assessed the levels of GM-CSF in their supernatants (**Fig. 6C**). The significant decrease in GM-CSF secretion observed in these cells in absence of proliferation defects (**Fig. S6G**) further confirmed the central role of the ERK1/2 signaling pathway activity in driving GM-CSF secretion.

**Figure 6.**
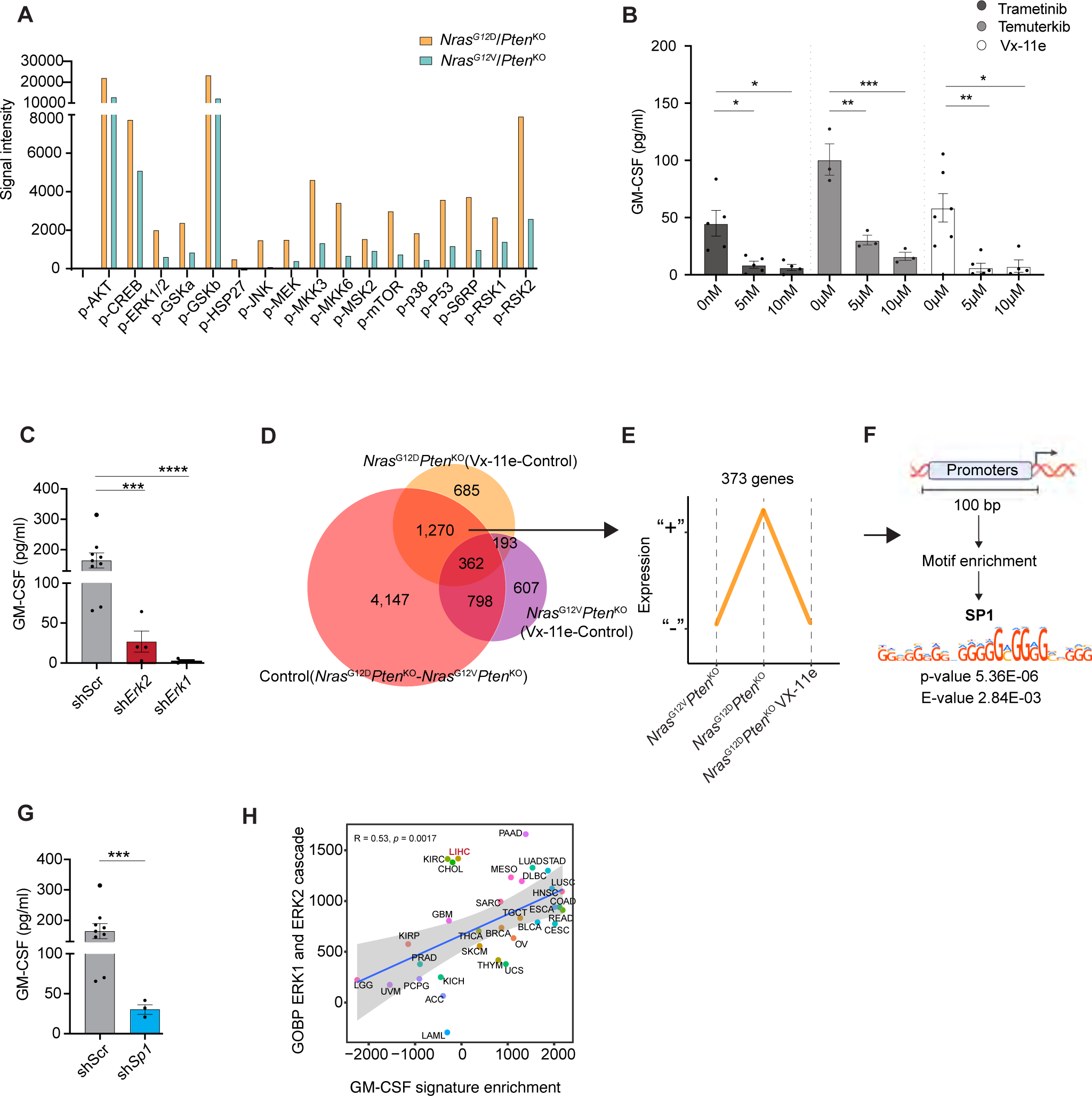
GM-CSF expression is regulated through activation of the ERK1/2 pathway and SP1 transcription factor. **A.** Barplot depicting the expression levels (signal intensity) of RAS/MAPK-associated phospho-proteins in *Nras*^G12D^/*Pten*^KO^ and *Nras*^G12V^/*Pten*^KO^ HCC cell lysates. **B.** Barplot depicting GM-CSF protein levels quantified in *Nras*^G12D^/*Pten*^KO^ cancer cell conditioned media after 24h of treatment with Trametinib (MEK1/2 inhibitor; n=4-5), Temuterkib (ERK1/2 inhibitor; n=3) and Vx-11e (ERK2 inhibitor; n= 4-6) at the indicated drug concentrations. **C.** Barplots depicting the GM-CSF protein level quantified in the supernatant of scramble (sh*Scr*, n=9), sh*Erk2* (n=4) and sh*Erk1* (n=4) *Nras*^G12D^/*Pten*^KO^ HCC cell lines. **D.** Venn diagram depicting the overlap of DEG between *Nras*^G12D^/*Pten*^KO^ and *Nras*^G12V^/*Pten*^KO^ cells treated or not with Vx-11e (**Supplementary Table 9A, E-F**). **E.** Diagram depicting the gene expression pattern (y-axis) across conditions (x-axis) that follow *Csf2* regulation (**D**; 1270 overlapping DEG). “+” indicates the up-regulated genes and “-“ indicates the down-regulated genes. **F.** Motif enrichment analysis in promoters from the 373 genes (**E**), identifying SP1 as a top candidate (**Supplementary Table 9G-H**). **G.** Barplots depicting the GM-CSF protein levels quantified in the supernatant of sh*Scr* (n=9) and sh*Sp1* (n=3) *Nras*^G12D^/*Pten*^KO^ HCC cell lines. **H.** Scatterplot depicting the correlation between the enrichment of ERK signaling pathway (y-axis) and GM-CSF signature (x-axis) for each TCGA cancer type. Correlation analyses were performed on the median pathway enrichment score of all patients per cancer type. Graph shows mean ± SEM (**B-C**, **G**). Statistical significance was determined by Student t-test (**B-C, G),** binomial test (**F**) and test for association between paired samples, using one of Pearson’s product moment correlation coefficient (**H).** *p < 0.05; **p<0.01, *** p < 0.001, **** p < 0.0001.

We next sought to identify ERK1/2 downstream factors regulating *Csf2* expression by performing RNA-seq on both *Nras*^G12D^*/Pten*^KO^ and *Nras*^G12V^*/Pten*^KO^ cells upon ERK signaling blockade (**Fig. S6H and Supplementary Table 9E-H**). We queried this dataset for changes unique to *Nras*^G12D^ point mutation following ERK inhibition (**Fig. 6D**) and selected promoters regulating genes which expression profiles followed the same trajectory than *Csf2* in the different samples analyzed (**Fig. 6E**). Next, we applied motif enrichment analysis on these promoters and identified the motif associated with the transcription factor SP1 as a top candidate (**Fig. 6F and Supplementary Table 9G-H**). To assess the role of SP1 in modulating *Csf2* expression and GM-CSF secretion, we generated a *Sp1* knockdown cell line (sh*Sp1*) from the parental *Nras*^G12D^*/Pten*^KO^ cells (**Fig. S6I**). Both GM-CSF gene expression and cytokine secretion were hindered in sh*Sp1 Nras*^G12D^*/Pten*^KO^ cells (**Fig. 6G and S6J**), positioning SP1 as an important regulator of the *Nras*^G12D^/MAPK-ERK/GM-CSF activation cascade. Finally, we validated the translational relevance of ERK1/2 and GM-CSF pathway correlation by assessing their transcriptional signature enrichment across all TCGA patients. Remarkably, the positive association between ERK1/2 pathway activity and GM-CSF signature was evident across several cancer types (**Fig. 6H**). Overall, this finding places ERK1/2 signaling node at the center of *Nras*^G12D^-driven GM-CSF secretion, while underscoring the broader impact of MAPK-ERK and GM-CSF axes at the pan-cancer level.

### *Nras*^G12D^/*Pten*^KO^-derived GM-CSF promotes the accumulation of proinflammatory monocyte-derived Ly6C^low^ myeloid cells

We next investigated the role of *Nras*^G12D^/*Pten*^KO^-derived GM-CSF in shaping myeloid cell differentiation and phenotype *in vitro*. We differentiated bone marrow (BM) cells using conditioned media (CM) prepared from *Nras*^G12D^/*Pten*^KO^ or *Nras*^G12V^/*Pten*^KO^ cancer cell lines (**Fig. S7A**), while recombinant M-CSF and GM-CSF were used for comparison. Interestingly, *Nras*^G12D^/*Pten*^KO^ CM specifically led to an accumulation of Ly6C^low^CD11b^high^F4-80^low^ cells, referred to as Ly6C^low^ herein (**Fig. 7A and S7B**) while other myeloid cell subset contents were largely unchanged (**Fig. S7C**). Analysis of the polarization profile of *Nras*^G12D^/*Pten*^KO^-induced Ly6C^low^ cells showed an enrichment of the immunosuppressive markers CD39 and PD-L1, a feature comparable to recombinant GM-CSF exposure (**Fig. 7B**) and corroborating with the phenotype of myeloid cells present within *Nras*^G12D^/*Pten*^KO^ HCC (**Fig. S3B**). In addition, the antigen presenting cell (APC)-related proteins CD80 and MHCII were similarly up-regulated (**Fig. 7B**), as previously reported in the context of GM-CSF stimulation^86^. Crucially, *Nras*^G12D^/*Pten*^KO^ CM-induced Ly6C^low^ cell abundance and education profile were both abrogated upon GM-CSF neutralization (**Fig. 7A, B**). We next investigated whether *Nras*^G12D^/*Pten*^KO^ CM induced gene expression changes in BM cells towards a more proinflammatory profile, as highlighted by the scRNA-seq changes within the *Nras*^G12D^/*Pten*^KO^ TME (**Fig. 4C, F**). We first confirmed the increased expression of GM-CSF downstream genes *Ccl6* and *Ccl17* in BM cells differentiated with *Nras*^G12D^/*Pten*^KO^ CM, which was abrogated upon GM-CSF neutralization (**Fig. S7D**). Similarly, expression of the proinflammatory genes *Il1a* and *Il6* were hindered in the context of GM-CSF blockade (**Fig. S7D**). Moreover, BM cells cultured in *Nras*^G12D^/*Pten*^KO^ CM displayed high IL-1R signaling activity which was reverted upon GM-CSF neutralization (**Fig. 7C**). Overall, these findings validate *Nras*^G12D^/*Pten*^KO^ cancer cell-derived GM-CSF as a central regulatory cytokine promoting the differentiation and re-education of Ly6C^low^ myeloid cells with mixed immunosuppressive and proinflammatory/APC-like features.

**Figure 7:**
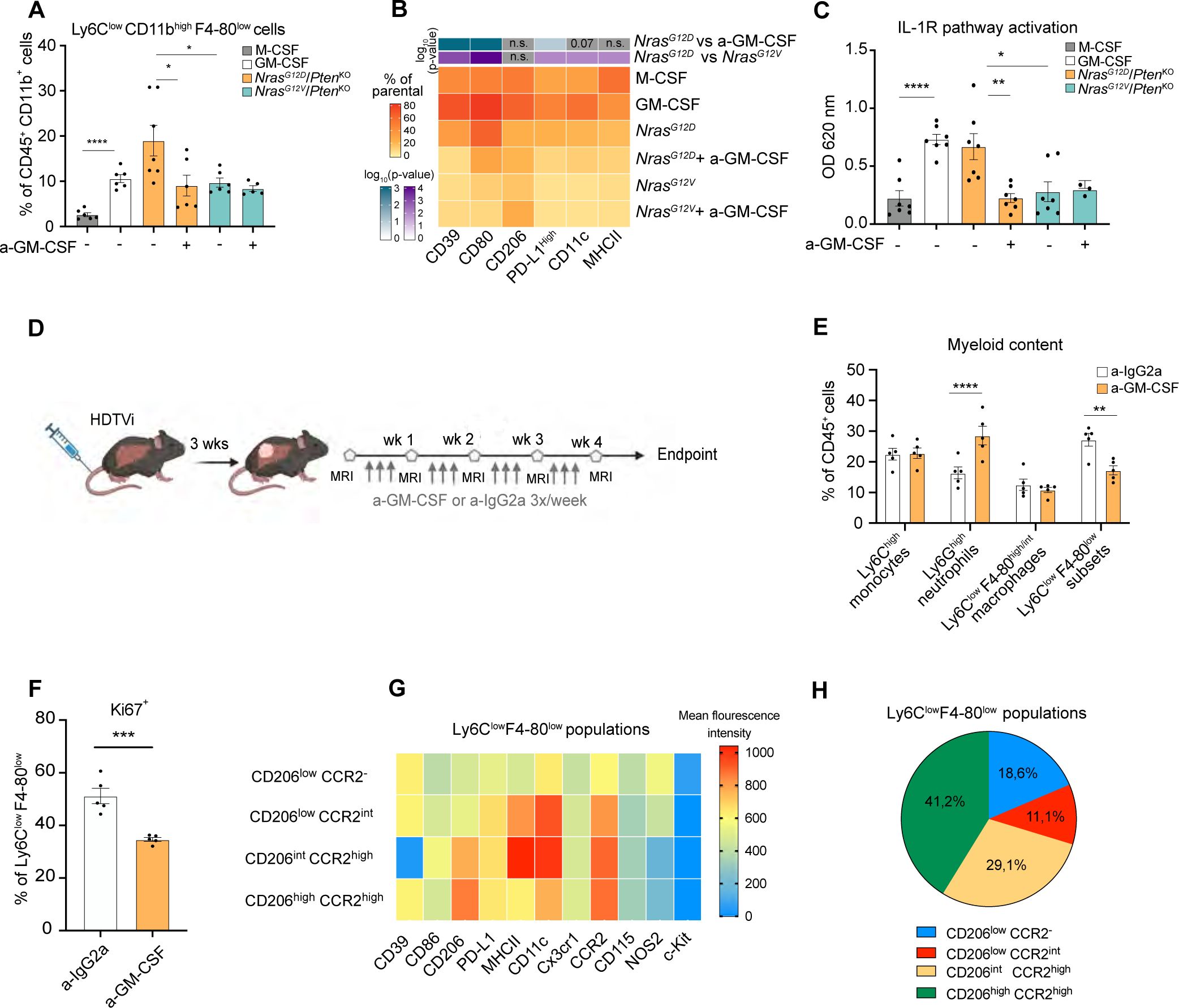
GM-CSF blockade curbs the accumulation of proinflammatory monocyte-derived Ly6C^low^ cells in the *Nras*^G12D^/*Pten*^KO^ HCC TME. **A.** Barplot depicting the percentage of Ly6C^low^F4/80^low^ cells relative to CD45^+^CD11b^+^ total myeloid cells obtained from bone marrow (BM) cells differentiated in either recombinant M-CSF or GM-CSF, or in conditioned media (CM) prepared from distinct HCC cell lines, with or without GM-CSF neutralizing antibody (a-GM-CSF) (M-CSF n=6, GM-CSF n=6, *Nras*^G12D^/*Pten*^KO^ n=7, *Nras*^G12D^/*Pten*^KO^ + a-GM-CSF n=6, *Nras*^G12V^/*Pten*^KO^ n=6, *Nras*^G12V^/*Pten*^KO^ + a-GM-CSF n=5). **B.** Heatmap depicting the median percentage of distinct BM-derived Ly6C^low^ F4/80^low^ cells expressing the indicated phenotypic markers. Percentage values shown for each phenotypic marker are relative to the total Ly6C^low^ F4/80^low^ population. **C.** Barplot displaying the IL-1R signaling activity in HEK reporter cells exposed to CM from BM cells differentiated with either recombinant M-CSF, GM-CSF, or to CM prepared from *Nras*^G12D^/*Pten*^KO^ or *Nras*^G12V^/*Pten*^KO^ HCC cell lines, in presence or not of a-GM-CSF (M-CSF n=7, GM-CSF n=7, *Nras*^G12D^/*Pten*^KO^ n=7, *Nras*^G12D^/*Pten*^KO^ + a- GM-CSF n=7, *Nras*^G12V^/*Pten*^KO^ n=7, *Nras*^G12V^/*Pten*^KO^ + a-GM-CSF n=3). **D.** Experimental design: HDTV injections were performed in C57/BL6 mice to induce *Nras*^G12D^/*Pten*^KO^ HCC. Mice were monitored by weekly MRI starting from 3 weeks post-injection, and enrolled into treatments with a-IgG2a (12.5 mg/kg three times per week) or a-GM-CSF (12.5 mg/kg three times per week) until timepoint sacrifice at 2 weeks after treatment initiation or end-stage (tumor volume ± 2cm^3^) for flow cytometry timepoint analyses. **E.** Barplots depicting the percentage of intratumoral Ly6C^high^ monocytes, Ly6G^high^ neutrophils, Ly6C^low^F4-80^high/int^ macrophages and Ly6C^low^F4-80^low^ subsets relative to total CD45^+^ leukocytes in HDTVi-induced *Nras*^G12D^/*Pten*^KO^ HCC-bearing mice 2 weeks post treatment with a-IgG2a (n=5) or a-GM-CSF (n=5). **F.** Barplot depicting the percentage of Ki67^+^Ly6C^low^F4/80^low^ cells in HDTV-induced *Nras*^G12D^/*Pten*^KO^ HCC sacrificed 2 weeks post treatment with a-IgG2a (n = 5) or a-GM- CSF (n = 5). **G.** Heatmap of the mean fluorescence intensity of the depicted phenotypic markers in four different Ly6C^low^F4-80^low^ subsets (CD206^low^CCR2^-^, CD206^low^CCR2^+^, CD206^+^CCR2^high^, and CD206^high^CCR2^high^) identified by FlowSOM analysis performed on HDTVi-induced *Nras*^G12D^/*Pten*^KO^ end-stage HCCs (n=3). **H.** Pie chart displaying the proportions of Ly6C^low^F4-80^low^ subsets (CD206^low^CCR2^-^, CD206^low^CCR2^+^, CD206^int^CCR2^high^, and CD206^high^CCR2^high^) identified by FlowSOM analysis performed on HDTVi-induced *Nras*^G12D^/*Pten*^KO^ end-stage HCCs (n=3) (see Fig. 7G). Graph shows mean ± SEM (**A**, **C**, **E**, **F**). Statistical significance was determined by unpaired Student’s T-test in (**A**, **C**, **E**, **F**) and log-rank test in (**B**). *p < 0.05; **p < 0.01; ***p < 0.001; ****p < 0.0001.

We next interrogated whether similar myeloid cell content and activation changes were recapitulated *in vivo* in the context of GM-CSF blockade (**Fig. 7D**). As observed *in vitro*, a-GM-CSF treatment led to a sustained decrease in the content of Ly6C^low^ myeloid cells within the *Nras*^G12D^/*Pten*^KO^ TME, observed as early as two weeks post treatment (**Fig. 7E and S7E;** for gating strategy, **Supplementary Data 1 and Fig. S7F**), whereas the content of lymphocytes was not altered (data not shown). No systemic changes were observed in peripheral myeloid cell content (**Fig. S7G**), suggesting that the effect of GM-CSF on reshaping the myeloid landscape is primarily local. Moreover, GM-CSF inhibition curbed the proliferation of Ly6C^low^ cells in the *Nras*^G12D^/*Pten*^KO^ TME (**Fig. 7F**), indicating that GM-CSF promotes the local expansion of this population. Furthermore, GM-CSF blockade compromised the mixed immunosuppressive/APC-like phenotype of Ly6C^low^ cells, with decreased expression of the immunosuppressive markers CD39 and PD-L1 and APC proteins CD80, MHCII and CD11c (**Fig. S7H).** Altogether, these results suggest that GM-CSF blockade dampens the abundance and the phenotype of Ly6C^low^ cells both *in vitro* and *in vivo*.

To investigate the cell of origin and further scrutinized the phenotype and function of the GM-CSF-driven Ly6C^low^ myeloid subset in *Nras*^G12D^/*Pten*^KO^ HCC, we probed for markers associated with distinct myeloid-derived cell types. FlowSOM analyses indicated that Ly6C^low^ cells comprised two dominant subsets, both characterized by heightened expression of CD206 and CCR2: CD206^high^CCR2^high^ and CD206^int^CCR2^high^ cells (**Fig. 7G-H**). Expression of CCR2 infers that the Ly6C^low^ pool is derived from a classical monocyte parental population^87^. The discrepant expression of CD206, a hallmark of immature DCs and a pro-tumorigenic macrophage marker^88- 91^, together with distinct levels of the APC proteins MHCII and CD11c ^92^, further highlight the mixed identity of the Ly6C^low^ pool presenting DC- and macrophage-like features (**Fig. 7G-H**). These results were corroborated by analyses of the C1_Monocytic_*March3*^+^ and C6_Monocytic_*Cxcl3*^+^ subclusters significantly expanded in the *Nras*^G12D^/*Pten*^KO^ TME (**Fig. 4D,E**), with both populations expressing *Ccr2* and *Mrc1* (CD206), low *Ly6c1*, *Ly6c2* and *Adgre1* (F4/80) levels, and distinct gene expression patterns of *H2-ab1*, *H2-eb1* and *Itgax* (CD11c) (**Fig. S7I**). Moreover, low expression of CD115 (Csf1r) and Cx3cr1 together with high CD11b levels in Ly6C^low^ myeloid cells (**Fig. 7G-H**) further supports their immature macrophage state^93^. Overall, our data indicates that the Ly6C^low^ myeloid cell pool is comprised of immature DC-like and macrophage-like monocytic cells that bear important pro-tumorigenic functions in *Nras*^G12D^/*Pten*^KO^ HCC. Indeed, loss of this myeloid cell subset upon GM-CSF blockade correlated with a significant decrease in tumor growth (**Fig. S7J**), underscoring this treatment as a potential therapeutic approach in *Nras*^G12D^/*Pten*^KO^ HCC.

### GM-CSF blockade curbs *Nras*^G12D^/*Pten*^KO^ HCC outgrowth and cooperates with VEGF inhibition to prolong animal survival

We next evaluated whether GM-CSF blockade would extend HCC-bearing mice survival. In light of the low penetrance of HDTV-induced *Nras*^G12D^/*Pten*^KO^ tumors (**Fig. S1B**), we developed liver orthotopic HCC models by injecting *Nras*^G12D^/*Pten*^KO^ cancer cells in WT C57/Bl6 mice (**Fig. 8A**) (herein referred to as the liver orthotopic injection-LOI-model). *Nras*^G12D^/*Pten*^KO^ LOI tumors exhibited comparable features to HDTVi-induced HCCs, with marked fibrosis (**Fig. S8A**), high GM-CSF levels (**Fig. S8B**), abundant myeloid cell content (**Fig. S8C**) and equivalent proportions of myeloid cell subsets within their TME (**Fig. S8D**). GM-CSF blockade reduced the expression of GM-CSF downstream genes *Ccl6* and *Ccl17* (**Fig. S8E, F**), asserting treatment efficacy, and substantially curbed tumor growth (**Fig. S8G**) leading to a significant increase in mice survival (**Fig. 8B**). Importantly, we observed that monocyte-derived Ly6C^low^ cell abundance was hindered in LOI tumors following a-GM-CSF treatment (**Fig. S8H**), in line with prior *in vitro* (**Fig. 7A**) and *in vivo* results (**Fig. 7E and S7E**). Mechanistically, the anti-tumorigenic effect of GM-CSF neutralization involved an increase in intratumoral cleaved caspase 3 (CC3) (**Fig. 8C, D**), decreased proliferation of parenchymal, non-immune cells (**Fig. S8I**) and lower proliferative capacity of Ly6C^low^ myeloid cells (**Fig. S8J**).

**Figure 8:**
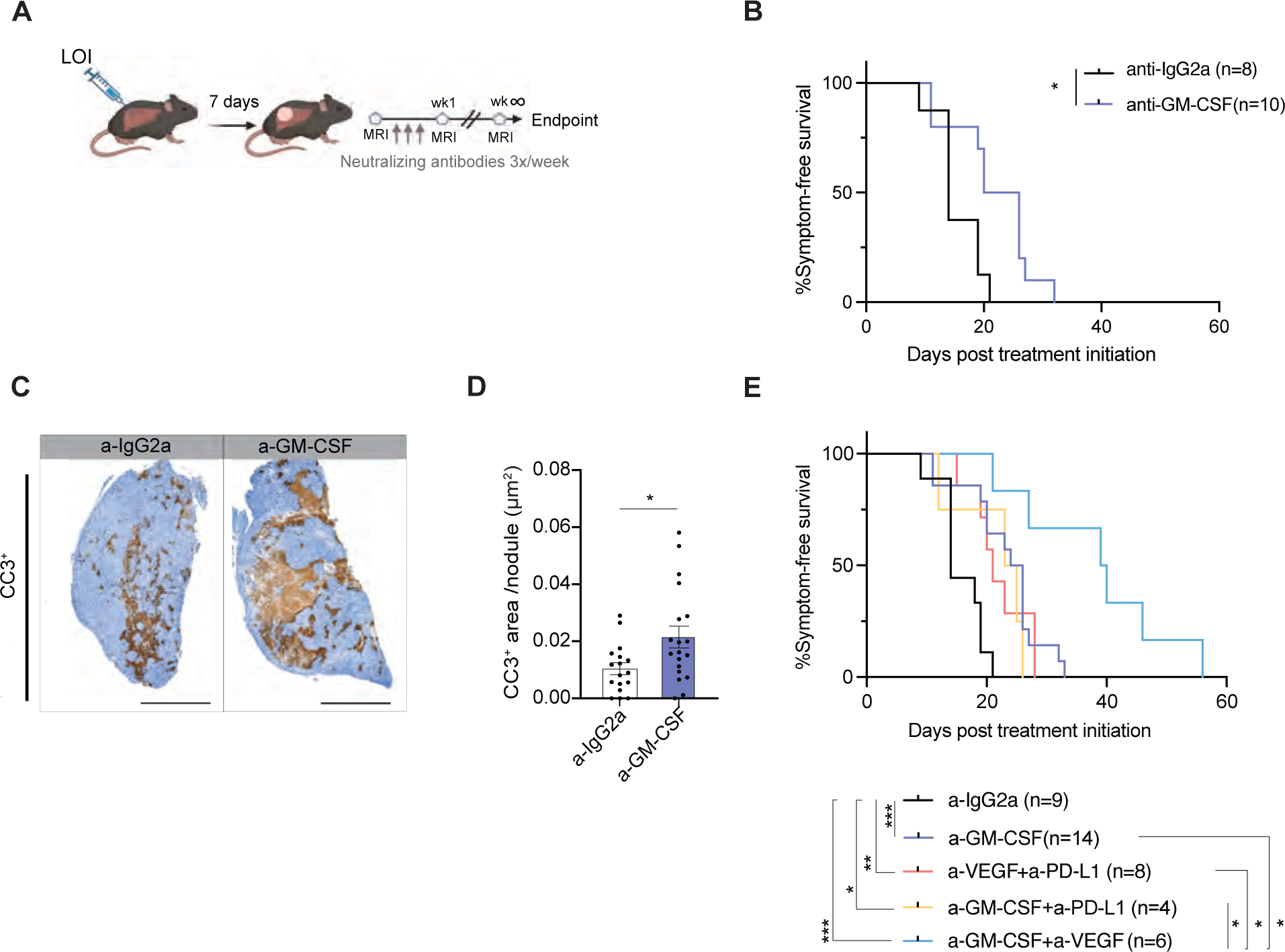
GM-CSF blockade hampers tumor growth and synergizes with VEGF inhibition to extend *Nras*^G12D^/*Pten*^KO^ HCC-bearing mice survival. **A.** Experimental design: liver orthotopic injection (LOI) were performed in C57/BL6 mice inoculated with 1×10^5^ cells *Nras*^G12D^/*Pten*^KO^ cancer cells. Mice were monitored by weekly MRI starting from 7 days post-surgery. Mice with visible tumors by MRI were enrolled into treatments with the following neutralizing antibodies (all three times per week): a-IgG2a (12.5 mg/kg), a-GM-CSF (12.5 mg/kg), a-PD-L1 (10mg/kg), or a- VEGF (10mg/kg) and sacrificed at end-stage (tumor volume ± 2cm^3^). **B.** Kaplan–Meier survival curves of LOI-induced *Nras*^G12D^/*Pten*^KO^ HCC-bearing mice treated with either a-IgG2a (n=8; median survival of 14 days) or a-GM-CSF (n=10; median survival of 23 days). **C.** Representative cleaved caspase 3 (CC3^+^) IHC staining performed on liver sections from end-stage LOI-induced *Nras*^G12D^/*Pten*^KO^ HCC-bearing mice treated with a-IgG2a or a-GM-CSF. Scale bars = 2 mm. **D.** Barplot depicting the cleaved caspase 3 (CC3^+^) positive area per μm^2^ in tumor nodules isolated from LOI-induced *Nras*^G12D^/*Pten*^KO^ HCC-bearing mice treated with either IgG2a (n =7) or a-GM-CSF (n=6). **E.** Kaplan–Meier survival curves of LOI-induced *Nras*^G12D^/*Pten*^KO^ HCC-bearing mice treated with either a-IgG2a (n=9; median survival of 14 days), a-GM-CSF (n=14; median survival of 25 days), a-VEGF + a-PD-L1 (n=8; median survival of 21 days), a- GM-CSF + a-PD-L1 (n=4; median survival of 24 days), a-GM-CSF + a-VEGF (n=6; median survival of 39,5 days). LOI-induced *Nras*^G12D^/*Pten*^KO^ HCC-bearing mice treated with a-IgG2a (n=8) and a-GM-CSF (n=10) shown in Fig. 8B are included in this graph. Graph shows mean ± SEM (**D**). Statistical significance was determined by unpaired Student’s T-test in (**D**) and log-rank test in (**B, E).** *p < 0.05; **p < 0.01; ***p < 0.001.

We next compared the therapeutic efficacy of GM-CSF blockade to that of the recently approved standard-of-care (SOC) HCC therapy comprising dual inhibition of VEGF and PD-L1. Interestingly, a-GM-CSF and SOC prolonged animal survival to a similar extent (**Fig. 8E**), highlighting the clear advantage of a-GM-CSF as a standalone treatment regimen in this disease. Next, we interrogated whether combining GM-CSF blockade with a-VEGF and/or a-PD-L1 could increase therapeutic response, as VEGF is strongly expressed in the *Nras*^G12D^/*Pten*^KO^ TME (**Fig. 4F and Fig. 5C**), while PD-L1 expression is heightened in *Nras*^G12D^/*Pten*^KO^ myeloid cells (**Fig. S3B**). Strikingly, the content of Ly6C^low^ cells was decreased in all treatment groups displaying improved survival benefit compared to IgG-treated *Nras*^G12D^/*Pten*^KO^ tumors (**Fig S8K**). Incorporating PDL-1 blockade to GM-CSF neutralization did not enhance therapeutic response. Contrastingly, dual GM-CSF and VEGF blockade significantly extended animal survival when compared to a-GM-CSF monotherapy or SOC (**Fig. 8E**). Altogether, these results suggest that a MAPK-ERK1/2-SP1-GM-CSF signaling node underlies several aspects of cancer cell-intrinsic pro-tumorigenic features, through increasing the abundance of monocyte-derived Ly6C^low^ cells in the *Nras*^G12D^/*Pten*^KO^ HCC TME and promoting a heightened inflammatory and immunosuppressive phenotype in this immature myeloid cell population. Interfering with these processes by blocking GM-CSF enforces a therapeutic vulnerability in the HCC TME that synergizes with the clinically-approved VEGF inhibition (**Fig. S8L**).

## Discussion

Faithfully recapitulating the HCC multilayered heterogeneity is essential to identify therapeutic approaches that can tackle the unmet clinical need of this disease. Here, we generated novel preclinical mouse models of HCC, each harboring genetic driver mutations altering clinically-relevant oncogenic signaling pathways that closely mimicked human HCC pathology and heterogeneity^4,71^. Notably, the myeloid-dominated *Nras*^G12D^/*Pten*^KO^- and *Myc*^OE^-driven HCCs (*Myc*^OE^*/Trp53*^KO^, *Myc*^OE^*/Pten*^KO^) correlated with the aggressive HCC proliferation subtypes, supporting the notion that myeloid cell recruitment enforces tumor aggressiveness^94^. *Myc*^OE^-driven models consistently clustered together, underlying the dominant role of Myc in shaping both cancer cell-intrinsic and extrinsic features of HCC fueling an immune-desert and immunosuppressive TME in several tumor types^17,18,21,26,56,95,96^.

While previous reports highlighted tissue-dependency and isoform specificity related to RAS signaling^50,97^, our study address the extrinsic effects of differential RAS/MAPK signaling pathway activation in cancer. Interestingly, G12D and G12V point mutations hold different enzymatic activity levels^98^, which might explain the differential MAPK signal intensity observed in the two *Nras*-driven models. Although RAS mutations are rare in HCC patients (∼1%)^99^, the RAS/MAPK pathway is overactivated in over 50% of them^46^, highlighting the translational potential of these findings. Herein, we revealed that varying degrees of RAS-MAPK signaling pathway intensity control unique biological and clinical behaviors in HCCs, with *Nras*^G12D^-driven HCC exhibiting a myeloid-enriched and fibrotic TME, and *Nras*^G12V^ eliciting a T cell-inflamed and lipid-enriched tumors, resembling NASH-like features. Of note, cancer cell-intrinsic lipid metabolic reprograming in *Nras*^G12V^/*Pten*^KO^ HCC may be therapeutically exploitable in future studies. Our results emphasize the relevance of investigating the activation states pertaining to a specific oncogenic signaling pathway to unravel the molecular bases of inter-patient heterogeneity.

The MAPK*-*enriched*, Nras*^G12D^*/Pten*^KO^-driven HCCs displayed a myeloid-rich TME, uniquely presenting a mixed inflammatory and immunosuppressive profile, thus highlighting the distinctive and yet under-appreciated immune-modulation capacity of the MAPK/ERK pathway hyperactivation through *Nras*^G12D^ point mutation. Indeed, cooperation of *Kras*^G12D^ and *Myc* oncogene was previously shown to fuel invasive lung adenocarcinoma immune-suppressed stroma^56^ while *Nras*^G12V^-induced senescent cells elicited a T cell response and modulated myeloid cell recruitment and maturity in the hepatic environment^35,55^. Complementing these reports, we unbiasedly identified GM-CSF as a key regulator of local myeloid cell education exclusively in *Nras*^G12D^- associated HCC, in a ERK1/2-SP1 dependent manner. In line with previous reports that identified GM-CSF as a pilot signal in *Kras*^G12D^-driven PDAC and cholangiocarcinoma^82,100^, we provide evidence of an analogous regulatory mechanism pertaining to distinct Ras isoforms in HCC. Moreover, our study unveils ERK1/2 and the downstream SP1 transcription factor as regulators of GM-CSF transactivation in HCC^101-103^. Hence, these findings shed light into novel alternative therapeutic targets exploitable in GM-CSF-enriched solid cancers.

Remarkably, while GM-CSF is known as a central mediator of myeloid cell function and differentiation^83-85^, we identified novel intricate connections between the GM-CSF and IL-1 signaling nodes specifying monocytic cell phenotype. We exposed a GM-CSF-driven local expansion of proinflammatory/immunosuppressive CCR2^+^ immature Ly6C^low^ myeloid cells within the *Nras*^G12D^*/Pten*^KO^ TME. The exact nomenclature for such subset is currently lacking, as these cells display characteristics associated with both macrophages and DCs, a confounding phenotype previously reported in progenitor cells exposed to GM-CSF *in vitro*^86^. Importantly, the anti-tumorigenic effects of GM-CSF neutralization involved the reduction and re-education of monocyte-derived Ly6C^low^ cells and the induction of cancer cell death in a T cell-independent manner. These results contrast with previous reports in *Kras*^G12D^-driven tumors^82,100^ in which T cells are central to GM-CSF blockade anti-tumor effect, suggesting the context dependency of this treatment. As inflammation is known to sustain cancer cell propagation and survival^77^, we propose that GM-CSF blockade hinders a myeloid-centric inflammatory cascade upheld by Ly6C^low^ immature cells. Nevertheless, acquired tumoricidal capacity in myeloid cells could also be hypothesized, as recently reported in metastatic disease following macrophage reprogramming^104^. Overall, our findings further expand the current appreciation of non-cell autonomous effects held by specific signaling pathways on the TME dynamics and therapy response. Indeed, recent generation of HCC somatic models were used to tailor anti-cancer therapies to HCC mutational background^26^. Here, we therapeutically exploited the myeloid-driven pro-tumorigenic effects orchestrated by GM-CSF to rewire the *Nras*^G12D^ TME, a process synergizing with the clinically-approved VEGF inhibition. As GM-CSF signature correlates with both poor patient prognosis and the *Nras*^G12D^*/Pten*^KO^-specific transcriptome, we advance the potential of GM-CSF neutralization as a novel myeloid-cell centric immunomodulation strategy in HCC, which can be further enhanced in combination with VEGF blockade standard of care. Recent changes in the treatment guidelines for advanced HCC patients have reinforced the importance of harnessing the TME as a successful therapeutic strategy^10^. Our study expands the understanding of HCC inter-tumor heterogeneity by revealing how cancer cell-intrinsic oncogenic pathways shape clinicopathological and TME profiles. Altogether, these findings set the basis for the design of personalized immune intervention strategies tailored to cancer cell-intrinsic features in HCC patients and expose novel treatment approaches for this disease.

## Methods

### HCC model generation and treatment

All animal experiments were reviewed and approved by the Animal Ethics Committee of the Netherlands Cancer Institute and performed in accordance with institutional, national and European guidelines for Animal Care and Use. All HCC mouse models were generated in C57BL/6J background from 6-8 weeks old females (Janvier laboratories).

HCC somatic mouse models were generated using hydrodynamic tail vein injections (HDTVi) as previously described^47,105^. Briefly, a volume equivalent to 10% of mouse body weight of sterile 0.9% NaCl saline solution containing plasmid mixtures was injected into the mouse lateral tail vein in 7-10 seconds. A total of 25 ug of DNA mixture was injected per mouse and prepared as followed: 10 ug of transposon vector, 10ug of CrispR/Cas9 vector, and 5 ug of Sleeping beauty (SB) transposase encoding vector (unless indicated). The *CMV-SB13*, the *pT3-EF1a-Myc* and the *pT3-EF1-Nras*^G12D^*-GFP* vectors were a kind gift from Dr Scott Lowe (Memorial Sloan Kettering Cancer Center, New York, USA). The *pT/CaggsNras*^G12V^*-IRES-Luc* vector^106^ was kindly provided by Lars Zender (University of Tuebingen, Tuebingen, Germany), the *pX330- Trp53* (Addgene 59910), the *pX330-Pten* (Addgene 59909), were previously validated and are publicly available. Mice were followed with weekly and bi-weekly MRI starting one-week post-HDTVi to measure the longitudinal progression of tumor volumes (as previously described^105,107^). Animals were sacrificed when symptomatic and/or when tumor volume reached a total volume ≥2 cm^3^ in survival curves and end-stage analyses (humane endpoint). For time-point analysis, HCC-bearing mice were sacrificed 1-2 weeks (*Myc*^OE^-driven models) and 2-3 weeks (*Nras*^OE^- models) post - tumor development, corresponding to a tumor volume ≥600mm^3^. Control mice underwent HDTVi of a DNA mixture prepared as described above with scramble empty vectors publicly available: *pT3-EF1-Neo-GFP* (Addgene 69134), and the *pX330-CBh- hSpCas9* (Addgene 42230) and were sacrificed 4 weeks (as referred to 4-weeks control) or 15 weeks (as referred to 15-weeks control) later.

For the generation of liver orthotopic (LOI) HCC mouse models, intrahepatic injection of HCC cell lines (from HDTVi-driven tumors) was carried out according to previously established protocols. Briefly, 1.0 × 10^5^ *Nras*^G12D^/*Pten*^KO^ cancer cells were resuspended in 5 μl of serum-free DMEM medium supplemented with 25% Matrigel (Corning, cat.no. 356230). Through an 8-mm midline, central incision, cells were slowly injected in the left-medial and/or left lateral liver lob using a microfine insulin syringe. Mice were followed with weekly MRI starting 7 days post-injection to measure tumor progression. For the a-GM-CSF, a-PDL-1, a-VEGF preclinical trials, when tumors were first visible by MRI, tumor size-matched animals (HDTVi- or LOI- generated) were randomized in the indicated treatment groups. Mice were treated three times a week intraperitoneally with a-IgG2A control (12.5 mg/kg BioXcell, BE0089), a-GM-CSF (12.5 mg/kg BioXcell, BE0259), a-PD-L1 (10 mg/kg BioXcell, BE0361), a-VEGF (10 mg/kg B20S, kindly gifted by Dr Iacovos Michael) and sacrificed at 2 weeks post-treatment (for HDTVi- *Nras*^G12D^/*Pten*^KO^) or at humane endpoint (for HDTVi- or LOI- *Nras*^G12D^/*Pten*^KO^).

#### Tissue microarray analyses of HCC patient samples

### Patient information

An independent cohort of 488 HCC patients who underwent primary curative resection between 2006 and 2010 were enrolled. The resection procedure and postoperative surveillance for recurrence were performed as described previously^65,108^. The Institutional Review Board of Sun Yat-sen University Cancer Center approved this study and all samples were anonymously coded in accordance with local ethical guidelines, as stipulated by the Declaration of Helsinki, with written informed consent obtained from all participants. The clinical characteristics of patient are summarized in Supplemental Table 4.

### IHC and image acquisition

Tissue microarrays of HCC samples were cut into 4 μm sections, and then processed for IHC according to our previous reports^65,108^. Primary antibodies against CD11b (ab133357, Abcam), CD15 (ZM-0037, ZSBio), S100A9 (34425, Cell Signaling Technology), CD204 (KT022, Transgenic) were used to determine myeloid cell subsets, p-ERK1/2 (ab214036, Abcam) and c-Myc (ab32072, Abcam) were used to evaluate the activation of signaling pathways. IHC-stained slides were scanned at ×20 magnification by digital pathology slide scanner (KFBIO).

### Image analysis

Cell numbers of myeloid subsets were estimated with object module of InForm Tissue Analysis Software (AKOYA). Myeloid response score was determined as described previously^65,108^. To define p-ERK1/2^+^ and c-Myc^+^ tumors, samples displaying unequivocal nuclei staining were classified as positive by 2 independent observers who were blinded to the clinical outcome.

### Cell lines, culture conditions and ex vivo experiment

*Myc^OE^*/*Trp*53^KO^, *Myc*^OE^/*Pten*^KO^, *Nras*^G12D^/*Pten*^KO^, and *Nras*^G12V^/*Pten*^KO^ HCC cell lines were generated and isolated from end-stage tumor-bearing mice of each distinct HCC mouse model as previously described^105^. Cells were grown on Collagen Type I rat tail (Corning, cat.no. 354236) pre-coated flasks/dishes and cultured using DMEM (Gibco, cat.no. 61965059) supplemented with 10% FCS, 1x penicillin/streptomycin (Roche) (referred to as complete medium).

AML12 cells were provided as a kind gift from Urszula Hibner (Institut de Genetique Moleculaire de Montpellier, France) and cultured with DMEM-F12+GlutaMax supplemented with 10% FBS (Capricorn, cat.no. FBS-12A), 1% penicillin/streptomycin (Roche) and 1% insulin-Transferrin-Selenium (ITS) (Gibco) (referred to as complete F12 medium)^109^.

HEK-Blue^TM^ cells (Invivogen cat. code hkb-il1r) were cultured in complete medium supplemented with 100 µg/mL Normocin according to the manufacturer’s instructions.

All cell lines were cultured at 37°C and 5% CO^2^ and tested for mycoplasma contamination using a MycoAlert^®^ mycoplasma detection kit (Lonza, cat: LT07-218). Only mycoplasma-negative cells were used.

### Generation of shRNA cell lines

The lentiviral PLKO.1-puro vectors containing a short hairpin RNA (shRNA) targeting *Erk1* (5’- AGGACCTTAATTGCATCATTA-3’), *Erk2* (5’GCTCTGGATTTACTGGATAAA-3’) and *Sp1 (5’* CCTTCACAACTCAAGCTATTT-3’) were obtained from the TRC library. Control (scrambled) vector was available through purchase (Addgene #136035). A total of 2 × 10^6^ HEK 293T cells were seeded and after 24 hours transfected with 1.5 μg of pLKO.1 vector encoding shRNAs, 1 μg of pPAX packaging vector and 1 μg of VSV-G envelope vector using FuGENE® HD Transfection Reagent (Promega, cat.no. E2311), according to the manufacturer’s instructions. 24 hours after transfection, supernatants were collected, filtered (0.45- mm pore size filter; Millipore), and added to *Nras*^G12D^/*Pten*^KO^ cells to be transduced for 16 h. After 48 hours, transfected cells were selected with puromycin (2.5 μg/ml) for 4 days.

### Generation of conditioned media

To generate conditioned media (CM), 5×10^6^ HCC cell lines or AML12 cells were seeded in complete medium and/or complete F12 medium, respectively. After 24 hours, medium was replaced with 0% FBS DMEM to generate CM. CM was collected and centrifuged at 1000 × g for 5 minutes to remove cell debris and stored at -80°C and subsequently used for bone marrow cell differentiation and cytokine array.

### Bone marrow cell differentiation

Bone marrow (BM) cells were freshly isolated from wild-type mice as previously described^31^. Briefly, both femurs and tibias were flushed in complete medium using a 23G needle. The cell suspension was filtered through a 100 μm cell strainer (Millipore) and cultured in 10ml Teflon bag (OriGen PermaLife) with either control conditions or CM generated from HCC cell lines (as explained above) supplemented with 2% FBS, in the presence or absence of 5μg/ml of a-GM-CSF (BioXcell, cat.no. BE0259). M-CSF and GM-CSF differentiated BM cells were cultured with complete medium +10ng/ml recombinant mouse M-CSF (Biolegend, cat.no. 576408) or +20ng/ml recombinant GM-CSF (Peprotech, cat.no. 315-03), respectively. Cells were maintained in culture for a total of 5 days and medium was refreshed every 2 days. On day 5, the cell suspension was harvested and centrifuged at 300 × g for 5 minutes. Supernatants were used for HEK-Blue^TM^ IL-1R assay. Cell pellets containing differentiated BM cells were used for RNA isolation and flow cytometry analyses.

### HEK-Blue IL-1R assay

IL-1R signaling activity was measured using HEK-Blue^TM^ IL-1R cells (Invivogen, cat. code hkb-il1r) following the manufacturer’s instructions. Briefly, differentiated BM cell supernatant was centrifuged at 1000 × g for 5 minutes to remove cell debris. 200 μl of 10x diluted-supernatant was added to 5 x10^4^ HEK-Blue^TM^ IL-1R cells seeded in 96- well plate. Complete medium supplemented or not with recombinant IL-1β (25ng/ml) (Abcam, cat.no. ab259421) was used as a positive and negative control, respectively. The next day, 20μl of HEK cell supernatant was transferred to a flat-bottom 96-well plate containing 180 μl of QuantiBlue (Invivogen, cat. code rep-qbs) detection reagent and incubated at 37°C for 30 minutes. IL-1R activity was determine with a spectrophotometer (Tecan) at 620 nm emission. Negative background was subtracted from the raw values.

### MAPK and PI3K/mTOR pathway inhibition

5×10^5^ HCC cell lines were seeded into 6-well plates with complete medium. The following day, cells were cultured in 0% FBS-DMEM for 24 hours with: Trametinib, MK2206, AZD8055, SP600125, Vx-11e and Temuterkib (S2673, S1078, S1555, S1460, S7709, S8534, Selleck Chemicals). Supernatants were collected to determine the secreted GM-CSF levels. Cell pellets were used for protein or RNA isolation.

Cell growth was determined by IncuCyte ZOOM (Essen BioScience) assays as previously described^107^. Briefly, 2×10^4^ *Nras*^G12D^/*Pten*^KO^ HCC cells were seeded in 48- well plates and treated with the MAPK and PI3K/mTOR pathway drugs mentioned above in DMEM-FCS 0%. Cells were imaged every 4 hours and phase-contrast images were analyzed to determine the relative cell growth based on cell confluency.

### RNA isolation, cDNA synthesis and RT-qPCR

RNA extracted from snap frozen intermediate and end-stage bulk tumors and from 4- to 15-week control HDTVi livers (∼5mg), from bulk tumor FACS-sorted myeloid cell populations, or from primary HCC cell lines was isolated using TRIzol (Thermo Fisher) according to the manufacturer’s instructions. RNA from educated BM cells was isolated using RNAeasy kit (Qiagen, cat.no. 74104).

For cDNA synthesis, the High-Capacity cDNA Reverse Transcriptase Kit (Thermo Fisher, cat.no. 4368814) was used with 500ng of RNA. The following Taqman probes (Thermo Fisher) were used for qPCR: *Ubc* (Mm01201237_m1), *Il6* (Mm00446190_m1), *Il1b* (Mm00434228_m1), *Il1a* (Mm00439620_m1), *Ccl6* (Mm01302419_m1), *Ccl17* (Mm00516136_m1), *Csf2* (Mm01290062_m1), *Nlrp3* (Mm00840904_m1), *Arg1* (Mm00475988_m1) and *Sp1* (Mm00489039_m1). Relative expression was calculated after normalization to the housekeeping gene *Ubc* for each sample.

### Protein isolation

Proteins were isolated from HCC cell lines using RIPA Lysis buffer (Thermo Fisher, cat.no. 89900) supplemented with Halt™ Protease and Phosphatase Inhibitor Cocktail (Thermo Fisher). Snap frozen tumor samples (∼5mg) were lysed in cOmplete™ Lysis- M lysis buffer (Roche, cat.no. 4719956001) supplemented with protease and phosphatase^107^. Protein lysates were sonicated and protein concentration was determined using Pierce™ BCA Protein Assay Kit (Thermo Fisher, cat.no. 23225) for subsequent analyses.

### Western blot

Equal concentration of proteins from total cell lysates (25 to 50 μg) were loaded on SDS-PAGE gels and transferred onto PVDF membranes. The membranes were blocked for 1 hour with 5% milk PBS + 0,05% Tween (PBS-T) and incubated overnight at 4°C with primary rabbit antibodies against p-AKT (4060S; CST; 1:1000 dilution), p-c-Jun (3270S; CST; 1:1000 dilution), p-ERK1/2 (4370S;CST;1:500 dilution), T-ERK1/2 (9102S;CST; 1:1000 dilution), p-S6RP (211S; CST; 1:1000 dilution), p-4EBP1 (9459S;CST; 1:1000 dilution), Ras-G12D (14429S; CST; 1:500 dilution), p-RSK-1 (9341S;CST; 1:1000 dilution), and vinculin (13901T;CST; 1:1000 dilution) Secondary conjugation was performed using anti-rabbit IgG, HRP-linked Antibody (CST, cat.no. 7074P2) for 1 hour at room temperature, and proteins were detected with Signal Fire™ ECL Reagent (CST, cat.no. 6883P3) using BioRad ChemiDoc™ XRS+ System. Bands from western blots were quantified using Image Lab Software (BioRad).

### Caspase-1 assay

To measure caspase-1 activity, end-stage bulk tumor and 4- to 15-weeks control HDTVi liver protein lysates (50 μg) were analyzed using Fluorometric Caspase-1 Assay Kit (Abcam, cat.no. ab273268) according to the manufacturer’s instructions. Signal was acquired using the TECAN plate reader at an emission of 505 nm and excitation of 400nm. Fold change values were obtained after normalizing the samples by dividing each value to one control liver sample following background subtraction.

### Cytokine array

Measurement of cytokine levels were assessed in HCC cancer cell’s CM, end-stage tumor bulk and 15-weeks control HDTVi liver lysates (1000 μg), and 5x diluted serum from end-stage HCC-bearing and 15-weeks control mice using the Proteome Profiler Mouse XL Cytokine Array (R&D systems, cat.no. ARY028) according to the manufacturer’s instructions. Pixel density was quantified using ImageLab software (Biorad). Each membrane array was normalized to its own reference control after subtracting the background signal. Fold change values of pixel densities were obtained by using control samples (AML12 for cell lines, 15-weeks control mice for serum and bulk analyses) as the baseline value (set to 1).

### Multiplex assay

GM-CSF level (shown for this assay in **Fig. S5I**) were assessed in end-stage HCC and 4- to 15-weeks control HDTVi liver protein lysates (350 μg) by Protavio Ltd with a 11-plex array, and analyzed according to the Luminex technology company’s protocol.

### Phospho protein profiling

To measure the phosphorylated levels of RAS-associated effector proteins, *Nras*^G12D^/*Pten*^KO^ and *Nras*^G12V^/*Pten*^KO^ cell line protein lysates (1000 μg each) were analyzed using the Proteome Profiler^TM^ Phospho-MAPK Array Kit (Raybiotech, cat.no. AAH-AMPK-1-2) according to manufacturer’s instructions. Pixel intensity was quantified using ImageLab Software (Biorad). Each membrane array was normalized to their own reference controls after subtracting the background signal.

### GM-CSF ELISA assay

To quantify the levels of secreted GM-CSF, CM was generated as mentioned above and GM-CSF was quantified using the mouse GM-CSF uncoated ELISA Kit (Thermo Fisher, cat.no. #88-7334-22) in accordance with the protocol provided by the manufacturer. GM-CSF concentration was normalized to the protein concentration of the respective HCC cell line lysate, quantified with Pierce™ BCA Protein Assay Kit (Thermo Fisher).

### Tissue collection and processing

Mice were euthanized by carbon dioxide asphyxiation. Blood was collected by heart puncture and subsequently transcardially perfused with PBS until liver is cleared of blood. Harvested livers were macro-dissected and used for further analysis^105^.

### Serum preparation

Blood was left to clot at room temperature for 15 minutes and then centrifuged at 2000 × g for 20 minutes at 4°C. Serum was collected from the supernatant and stored at - 80°C until further use.

### ALT activity

To assess hepatocellular damage, ALT activity was measured with the Alanine Transaminase Activity Assay Kit (Abcam, colorimetric, cat.no. ab105134) using the serum (5x diluted) of intermediate- and end-stage HCC-bearing and 4- to 15-weeks control mice according to the manufacturer’s instructions. Signal was acquired using the TECAN plate reader at 570nm emission and 37°C in 5-minute intervals during a total scanning time of 90 minutes. Negative background was subtracted from the raw values.

### AFP ELISA assay

To quantify the levels of AFP at the systemic level, Mouse alpha Fetoprotein ELISA kit (Abcam, colorimetric, cat.no. ab210969) was performed using the serum (5x diluted) of intermediate-stage HCC-bearing and 4- to 15-weeks control mice according to the manufacturer’s instructions. Signal was acquired using the TECAN plate reader at 450nm emission. Negative background was subtracted from the raw values.

### Tissue imaging and analysis

Murine liver specimens were obtained from HCC end-stage and control-15-weeks mice and processed for immunohistochemistry (IHC) and histochemistry (HC) analysis as previously described^105^. Briefly, formalin-fixed, paraffin-embedded samples were sectioned at 4 μm and either probed with the indicated antibodies listed in **Supplementary Table 10** or stained with Masson’s trichrome dye for collagen fibers staining. Alternatively, samples were frozen down in Optimal Cutting Temperature (OCT) compound (Tissue-Tek), sectioned at 10 μm and stained with Oil red O dye for lipid staining.

All stained slides were digitally processed using the Aperio ScanScope (Aperio, Vista, CA) at a magnification of 20x. Immunohistochemical and histochemical staining were performed by the Animal Pathology facility at the Netherlands Cancer Institute. Histopathological evaluation of HCC tissue samples was performed on H&E, ARG1, Masson’s trichrome and Oil red O-stained slides by an experienced liver pathologist (dr. Joanne Verheij, Amsterdam UMC). For the analysis of CC3 stained HCC tumor samples, nodule size and CC3+ areas were drawn by hand on HALO image-analysis software (Indica Labs) to quantify the CC3+ area/nodule per mm^2^.

### Flow cytometry

Flow cytometry procedures were performed as previously described^105,107^. Briefly, macrodissected HCC nodules and control liver samples were dissociated as single-cell suspensions using the Liver Dissociation kit (Miltenyi Biotec) and the gentleMACS Octo Dissociator following the manufacturer’s instructions. Samples were incubated with anti-CD16/CD32 antibody (BD Bioscience) and stained with the antibodies against surface markers following standard procedures. Samples were fixed with the eBioscience fixation and permeabilization kit (Invitrogen), and stained for intracellular markers (see **Supplementary Table 11**). All antibodies used for flow cytometry were titrated in a lot-dependent manner and are listed in **Supplementary Table 11**. All analyses were completed at the Flow Cytometry Core facility at the NKI. For blood, intermediate- and end-stage tumor analyses a four-laser BD Fortessa instrument (BD Bioscience, BD FACSDiva software v 8.0.2) was used. For the end-stage multidimensional data visualization and analyses in **Fig. 7**, a five-laser Aurora spectral flow cytometer (Cytek Biosciences) was used. Data was analyzed using the FlowJo v10 software. The Ly6C^low^F4-80^low^ populations obtained in the analyses was downsampled using the DownSample 3.3.1 plugin (FlowJo Exchange). The FlowSOM 3.0.18 and UMAP 3.1 plugins were used to analyze the Ly6C^low^F4-80^low^ populations from a concatenated dataset.

### Fluorescence-activated cell sorting

All sorting experiments were performed at the Flow Cytometry Core facility of the NKI with a BD FACSAria Fusion sorter (BD Bioscience) using a 100 μm nozzle.

For RT-qPCR gene expression analysis, macrodissected HCC nodules and 4- to 15- weeks control livers were dissociated in a single cell suspension and stained with antibodies as listed in **Supplementary Table 11**. Viable cells (NIR^low^) were sorted in 2% FBS-PBS based on CD45^+^CD11b^+^Ly6G^−^Ly6C^high^ and CD45^+^CD11b^+^Ly6C^−^Ly6G^−^F4-80^int^ to isolate Ly6C^high^ monocytes and MDMs, respectively. The sorted cells were centrifuged at 300 × g for 5 minutes at 4°C, washed with PBS and stored at -80°C in TRIzol (Thermofisher) for RNA isolation.

For transcriptomic analysis of the immune cell content, single viable cells (NIR^low^) were sorted based on CD45^+^ expression. 2% FBS-PBS sorted cells were centrifuged at 300 × g at 4°C and the cell pellets were stored at -80°C for RNA sequencing.

For single-cell RNA sequencing (scRNA-seq), three control mice (4 weeks post-HDTVi) were digested and pooled in a single cell suspension. Samples were stained with live/dead staining using Zombie NIR (Biolegend) and CD45 and CD11b (see **Supplementary Table 11**). Single viable (NIR^low^) cells were sorted based on CD45 expression (negative and positive) in 2% FBS-PBS. Sorted cells were washed with PBS with 0,04% BSA (PBS-BSA), and purified CD45^+^ and CD45^-^ cells were combined in a 1:1 ratio at the final concentration of 1000 cells/μl in PBS-BSA for scRNA-seq library preparation.

### Next generation *sequencing*

### RNA-seq

End-stage macro-dissected HCC nodules, control livers and cell lines non-treated or treated with Vx-11e were processed for RNA-seq as follows. Tissue samples were homogenized in 600μl of buffer RLT (79216, Qiagen) with the addition of 1% B-mercaptoethanol using the TissueLyserII (85300, Qiagen) in combination with 5mm stainless steel beads (69989, Qiagen). Cells were lysed with 350ul of buffer RLT (79216, Qiagen) and the total RNA was isolated using the RNeasy Mini Kit (74106, Qiagen), according to the manufacturer’s instructions. The library preparation for bulk tumors and cell lines was generated using the TruSeq Stranded mRNA sample preparation kit (Illumina Inc., San Diego, RS-122-2101/2) according to the manufacturer’s instructions (Illumina, Document # 1000000040498 v00). The libraries were sequenced with 54 paired-end reads on a NovaSeq-6000 using a Reagent Kit v1.5 (Illumina Inc., San Diego). For CD45^+^ RNA-seq and the cell line RNA-seq, the libraries were generated using Smart-Seq2 RNA (Illumina) according to the manufacturer’s instructions and paired-end sequenced using the Nextseq-550 High Output Kit v2.5.

### Whole exome sequencing (WES)

For WES, macro-dissected HCC were weighted and equal amounts (∼1mg each) from end-stage (n=3 each genotype) HCC-bearing mice and control liver (n=3). Following DNA extraction and fragmentation (200-300 bp), a maximum of 1 μg of sheared DNA was used for library preparation using the KAPA HTP Prep Kit (KAPA Biosystems, KK8234) following manufacturer’s instructions. The samples were 100bp paired-end sequenced Illumina Novaseq 6000.

### scRNA-seq

Chromium Controller platform of 10X Genomics was used for single cell partitioning and barcoding. Single Cell 3’ Gene Expression were prepared according to the manufacturer’s protocol “Chromium NextGEM Single Cell 3’ Reagent Kits v3.1” (CG000315, 10X Genomics). A NovaSeq 6000 Illumina sequencing system was used for paired end sequencing of the Single Cell 3’ Gene Expression libraries at a sequencing depth of approximately 17.000 mean reads per cell for control liver and 35.000 mean reads per cell for *Nras*^G12D^/*Pten*^KO^. NovaSeq 6000 paired end sequencing was performed using NovaSeq SP Reagent Kit v1.5 (cat# 20028401, Illumina) and NovaSeq S1 Reagent Kit v1.5 (cat# 20028319, Illumina).

#### Bioinformatic analyses

All the analyses were performed using R (v.4.1.1, 2021-08-10), unless mentioned otherwise. Plots were generated using the package *ggplot* (v.3.3.6) or *ggpubr* (v.0.4.0), except where mentioned.

### RNA-seq

Bulk tumors, CD45^+^ and cell lines RNA-seq were analyzed after paired-end sequencing reads were trimmed with *seqpurge* and aligned with *Hisat2* using as reference GRCm38 (mm10) (--min-intronlen 20 --max-intronlen 500000 -k 5 --minins 0 --maxins 500 --fr --new-summary --threads 16). The read counts were generated using the function *gensum* with gtf (v. 100) annotation. PCA was performed to verify accordance between replicates. The differential expression analysis was performed using *EdgeR*^110^. The samples were trimmed-mean-of-M-values (TMM) normalized to account for library size. To verify replicate agreement, the Pearson correlation was calculated on the TMM-normalized log2(TPMs) of each replicate. To test for differential expression, the generalized linear model method using the *glmLRT* function was used with a significance cutoff of FDR 1%. The log2(TPMs) were also calculated using the normalized read counts. The signatures for each genetically-distinct HCC bulk tumor samples were defined as the DEG between each model relative to the HDTVi control liver.

### Classification of murine HCC subtypes according to human molecular HCC subtypes

For the prediction of classification, the algorithm *NearestTemplatePrediction* (v.4 2015-12-02) was used on the Gene Pattern public server^111^. The input files of each of the replicates from the RNA-seq of the bulk tumor HCC models or CD45^+^ cells were TMM-normalized TPMs in a GCT format, the classes of the different classifications were based on published references^63,64,71,99,112-115^. Statistical significance cutoff was defined as p-value < 0.01 with multiple test correction using the Bonferroni method.

### Human TCGA-LIHC data analysis

TCGA-LIHC data was obtained from http://gdac.broadinstitute.org/ using the package *UCSCXenaTools* (v.1.4.8)^116^. The raw counts were processed using *EdgeR* (v.3.36.0)^110^ to obtain TMM-normalized log2(TPMs). Only patients with clinical and expression data were considered for downstream analysis (n=372). The segregation of patients by high/low correlation was determined by calculating the Pearson correlation between the signature of the genetically-distinct HCC models and the TCGA patients using their log2(TPMs). To define the patients with high correlation, the cut-off was a correlation ≥ the 3rd quantile of the data (n=93), whereas the low correlation patients were selected based on a correlation ≤ the 1st quantile (n=93). The conversion of mouse gene symbols to human was based on homology using ensembl^117^. For the calculation of the survival curves, the Kaplan-Meier estimator was obtained with the *survfit* function from the package ‘survival’ (v.3.2.11, https://github.com/therneau/survival). The statistical test applied was the log-rank test using the ‘survminer’ package (v.0.4.9, https://github.com/kassambara/survminer). The correlation between the LIHC patients per iCluster and the models was performed on the log2(TPMs) of all genes. The highest Pearson correlation was used as the determinant value to define iCluster for each murine model. Significance was tested on the Pearson’s coefficient. The function *Heatmap* from the library ‘ComplexHeatmap’ (v.2.10.0) was used to plot the heatmaps with the gene expression per patient. The gene expression is z-scored normalized per gene.

### Enrichment analysis

The enrichment of MAPK (KEGG_MAPK_SIGNALING_PATHWAY), PIK3 (REACTOME_PI3K_AKT_SIGNALING_IN_CANCER) and MYC (HALLMARK_MYC_TARGETS_V1) signaling pathways in the bulk tumor RNA-seq was performed using the *HypeR* (v.1.10.0) package^118^ with an hypergeometric test and for TCGA-LIHC patients, we applied single sample gene set enrichment analysis (ssGSEA v. 10.0.11)^119^ The gene set enrichment analysis in the CD45^+^ RNA-seq was performed with GSEA (v.4.2.2) using immune genesets^72^ to define the pan-immune categories with default parameters (permutation type: gene_set; number of permutations: 1000). To determine the enrichment of Biocarta and KEGG pathways, we applied single sample gene set enrichment analysis (ssGSEA v. 10.0.11)^119^ on the Gene Pattern public server^111^. The enrichment of GM-CSF signature (*Csf2rb*^KO^ - WT;^85^) was performed using the *Hyper* (v.1.10.0) package^118^ with an hypergeometric test.

### Protein-protein interaction analysis

The protein-protein interaction analysis was performed using Metascape^120^. In short, a network is created using databases for protein interactions. The resulting network is then tested for pathway enrichment.

### scRNA-seq analyses

The sequencing reads were mapped using Cell Ranger (6.1.2) to genome build GRCm38 (mm10). All downstream processing and analysis were performed using the R package *Seurat* (4.1.1). The counts were loaded into R and to remove low quality cells, only features (genes) detected in a minimum of three cells and cells expressing a minimum of 200 features (genes) were considered for the further quality control. To further filter out low quality cells with high percentage of mitochondrial reads and aberrantly low/high number of gene expressed depending on each sample, we used the median absolute deviation (MADs) higher (for mitochondrial percentage) or lower (for the number of genes detected) than the median value for these metrics to remove cells. A total of 12,313 cells remained and the samples were SCT normalized and integrated using the *IntegrateData()* function. The data was reduced and clustered (Louvain algorithm, 0.075 resolution) which were then annotated as Neutrophils, Monocytic cells, cDC1s, pDCs, B cell, T cell and NK cells (see **Supplementary Table 7**). The differentially expressed genes were calculated with the function *FindAllMarkers.* Re-clustering of the monocytic cells and the cDC1 using 0.25 resolution. The differential abundance was determined using the *MiloR* (v.1.2.0) package (https://marionilab.github.io/miloR). Briefly, this method is based on assigning cells to ‘neighborhoods’ from a k-nearest neighbor graph; the counts are then fitted using a negative binomial General Linear Model. This model is then used to test for differential abundance of cell proportions in the ‘neighborhoods’. The functions *enricher* and *compareCluster* from the R package clusterProfiler (v.4.2.2) were used to calculate the enrichment of the up-regulated DEG between *Nras*^G12D^/*Pten*^KO^ relative to control using the MSigDB Hallmarks database.

To integrate the annotation from both human HCC and our murine scRNA-seq, the function *FindTransferAnchors()* and *MapQuery()* were used from *Seurat* (4.1.1). The predicted annotations from Xue *et al*.^71^ were projected on our UMAP to confirm correct annotation of the cell types.

### Whole exome sequencing analysis

The paired-end reads were trimmed using SeqPurge and aligned with Burrows– Wheeler alignment (v.0.5.10) to GRCm38 (mm10). PCR duplicates were designated using rumidup (https://github.com/NKI-GCF/rumidup). For basecall recalibration and variant calling, the BaseRecalibrator and Mutect2 functions from the genome analysis toolkit^121^ (GATK; v.4.2.6.0) and were applied, respectively. SNPEff^122^ and SnpSift^123^, (v.5.1d), were used to annotate and select the somatic variants calls with TLOD greater or equal than 15 and filtered on the disruptive effects (conservative inframe deletion, disruptive inframe deletion, disruptive inframe insertion, frameshift variant, missense variant, start lost, stop gained, stop lost or stop retained variant). The counts of variants generated by SNPEff were used to calculate the TMB as follows: n * 1000000 / covered_bases, where the covered_bases were the number of bases that had at least 5x coverage, which varied between 37.47M to 37.48M bases per sample The TMB from human LIHC was obtained from https://www.cbioportal.org/ from different datasets^124-126^, and the simple nucleotide variation data was downloaded with TCGAbiolinks^127^ (v. 2.22.4). The variant types and base substitutions were extracted using maftools^128^ (v. 2.10.05). Analyses were done using the statistical programming language R version 4.0.3.

### Transcription factor motif analyses

The genes of interest were defined as the DEG between *Nras*^G12D^/*Pten*^KO^ cells and *Nras*^G12V^/*Pten*^KO^ that overlapped changes in Vx-11 treated compared to control in *Nras*^G12D^/*Pten*^KO^ cells and that excluded DEGs of *Nras*^G12V^/*Pten*^KO^cells also treated with Vx-11 (see **Supplementary Table 9E-F**). The genes selected for motif enrichment were both up-regulated in *Nras*^G12D^*/Pten*^KO^ in Vx-11e cells and compared to *Nras*^G12V^*/Pten*^KO^ cells. The motif enrichment was performed in the promoter regions extracted with *get_biomart_promoters* function from PromoterOntology (v.0.0.1) of mm10. The Simple Enrichment Analysis (SEA v. 5.4.1) tool from the MEME suite^129^ (sea --verbosity 1 --oc. --thresh 10.0 --align center –p file.fa --m db/MOUSE/HOCOMOCOv11_full_MOUSE_mono_meme_format.meme).

## Acknowledgements

We thank Drs L. Zender and S. Lowe for the kind gift of the *Ras* plasmids, U. Hibner for providing the AML12 cell line and I. Michael for providing the B20S VEGF neutralizing antibody. We thank all the members of the Akkari lab for insightful comments and discussion, and Jules Gadiot, Tamara Filipovic, Martina Färber, Naz Kocabay and Cankat Ertekin for excellent technical support. We are grateful to the facilities of the Netherlands Cancer Institute: Flow Cytometry, Animal Laboratory, Mouse Clinic Imaging Unit, Experimental Animal Pathology and Genomics. We thank members of the Karin de Visser and the René Bernards laboratories for insightful discussion during the preparation of the manuscript and for sharing reagents. This research was supported by the Dutch Cancer Society (KWF 12049 to L.A. and KWF 13476 to S.V.), Oncode Institute (L.A.), Cancer Genomics Center (L.A.) and of the Dutch Ministry of Health, Welfare and Sport. Figures **1A**, **7D**, **8A** and **S7A** were created with BioRender (https://biorender.com/).

## Author Information

These authors contributed equally: Christel FA Ramirez, Daniel Taranto and Masami Ando-Kuri.

## Author Contributions

L.A., S.V, C.F.A.R and D.T conceived the study, designed experiments, interpreted data. L.A., S.V., D.T., C.F.A.R and M.A.K wrote the manuscript. C.F.A.R, D.T, S.V, M.H.P.G and E.T. performed and analyzed experiments. M.A.K. performed all computational analyses. D.J.K. contributed to the CD45^+^ RNAseq analysis. D.G. and R.J.C.K. performed the WES analysis. Z.H. and J.X. performed and analyzed the HCC patient TMA. J.V performed histological analyses. All authors edited or commented on the manuscript.

## Data availability

Raw RNA-seq, scRNA-seq and WES data can be found in the Gene Expression Omnibus (GEO) database under the accession number GSE234897. Further information and requests for resources and reagents should be directed to the corresponding authors.

## Ethics Declarations

The authors have no competing interests to declare.

## Supplementary Figure Legends

**Figure S1:**
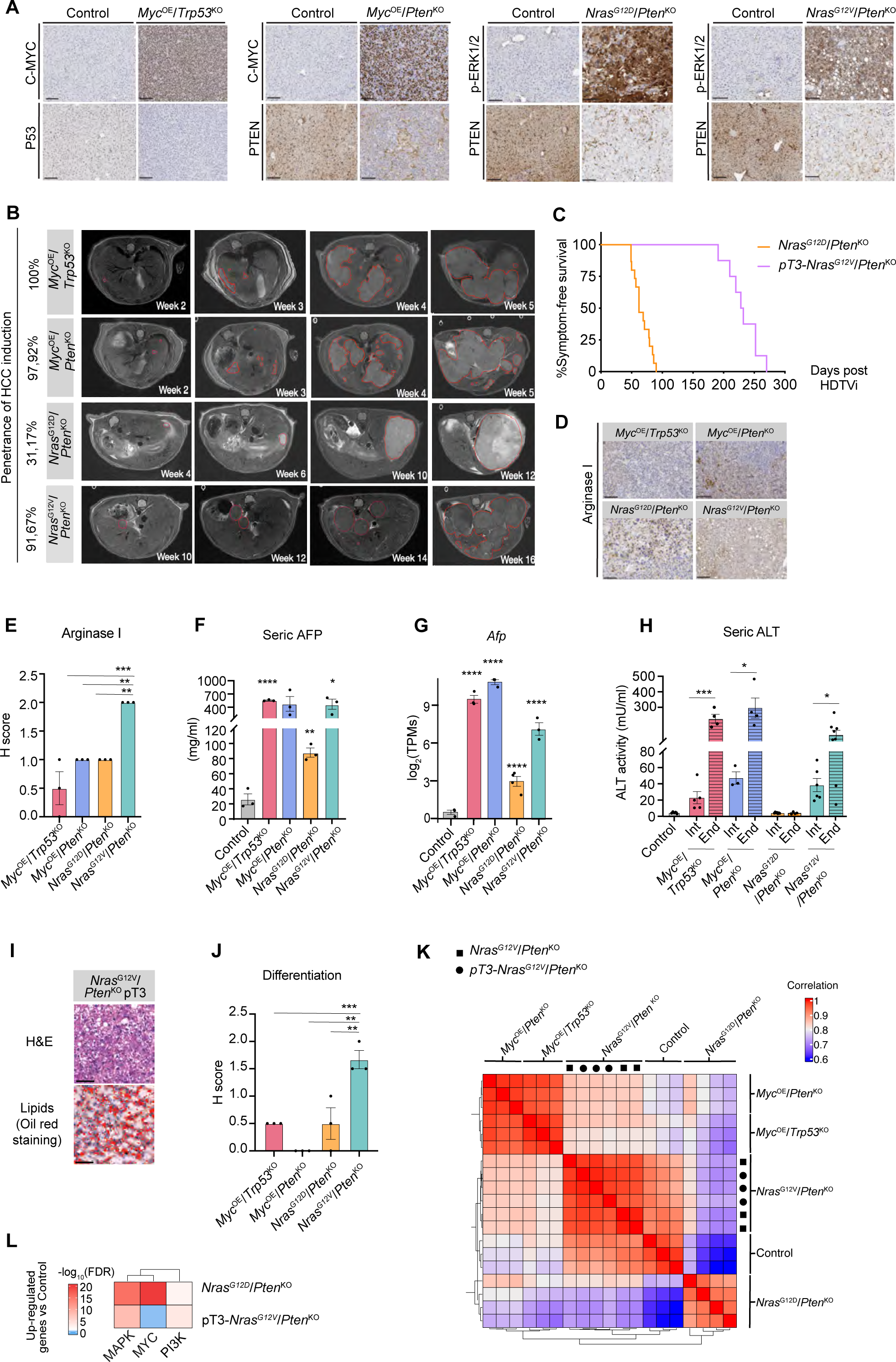
Genetically-distinct HCC display histopathological, systemic and molecular features of malignant transformation. **A.** Representative IHC images of C-MYC, P53, PTEN and phospho-ERK1/2 staining performed on liver sections from genetically-distinct HCC-bearing and control mice. Scale bars = 100 µm. **B.** T2-weighted MRI scans illustrating the penetrance of HCC induction, the tumor latency and growth in the genetically-distinct HCC models. Tumors were monitored weekly/bi-weekly one-week post-HDTVi. Red lines indicate the regions of interest used to calculate the tumor volumes (see **Methods**) presented in Fig. 1C. **C.** Kaplan–Meier survival curves of HCC-bearing mice HDTV injected with *Nras*^G12D^/*Pten*^KO^ (n=15; median survival of 62 days shown in **Fig 1B**), and *pT3- Nras^G12V^*/*Pten*^KO^ oncogenic drivers (n=8; median survival of 230 days). **D.** Representative IHC images of Arginase-1 performed on liver sections from genetically-distinct end-stage HCC-bearing mice. **E.** Barplot depicting the histopathology scoring of Arginase-1 expression assessed in genetically-distinct, end-stage HCC (presented in (**D**); n=3 for all HCC models). **F.** Barplot depicting AFP quantification performed in the sera of control and genetically-distinct HCC-bearing mice at intermediate stage; (n=3 for all HCC models). **G.** Barplot showing *Afp* gene expression in control and genetically-distinct end-stage HCC from the bulk RNA-seq; (n=3 for all HCC models). **H.** Barplot depicting ALT activity in the sera of control and genetically-distinct HCC-bearing mice; (*Myc*^OE^/*Trp53*^KO^ intermediate [Int] n = 5 and end [End]-stage n = 4, *Myc*^OE^/*Pten*^KO^ Int n= 3 and End n = 4, *Nras*^G12D^/*Pten*^KO^ Int n= 7 and End n = 4, *Nras*^G12V^/*Pten*^KO^ Int n= 6 and End n = 8 (see **Methods** for intermediate and end-stage definition). **I.** Representative images of H&E and Oil Red staining performed on sectioned livers collected from end-stage *pT3-Nras^G12V^*/*Pten*^KO^ tumor-bearing mice. Scale bars = 100 µm. **J.** Barplot depicting the differentiation status of genetically-distinct end-stage HCC tumors according to the 2019 WHO classification histopathology grading guidelines (see **Methods**); (n=3 for all HCC models). **K.** Heatmap of unsupervised hierarchical clustering depicting the Pearson correlation calculated from the normalized Transcripts per Million (TPMs) of the genetically-distinct HCC tumors and control (n=3 for all HCC models). Squares represents *Nras*^G12V^/Pten^KO^ and circles represents pT3- *Nras*^G12V^/Pten^KO^. **L.** Heatmap of unsupervised hierarchical clustering depicting the geneset enrichment of the MAPK, PI3K and MYC signaling pathways in *Nras*^G12D^/*Pten*^KO^ (shown in **Fig. 1F**) and pT3-*Nras*^G12V^/*Pten*^KO^ HCCs compared to control. The color scale represents the significance of the enrichment in –log10(FDR). FDR False Discovery Rate. Graphs show the mean ± SEM (**E-H, J**). Statistical significance was determined by one-way ANOVA with Tukey’s multiple comparison test (**E, J**), differential expression analyses (**G**) and unpaired Student’s *t*-test in (**F, H**). *p < 0.05; **p < 0.01; ***p < 0.001; ****p < 0.0001.

**Figure S2.**
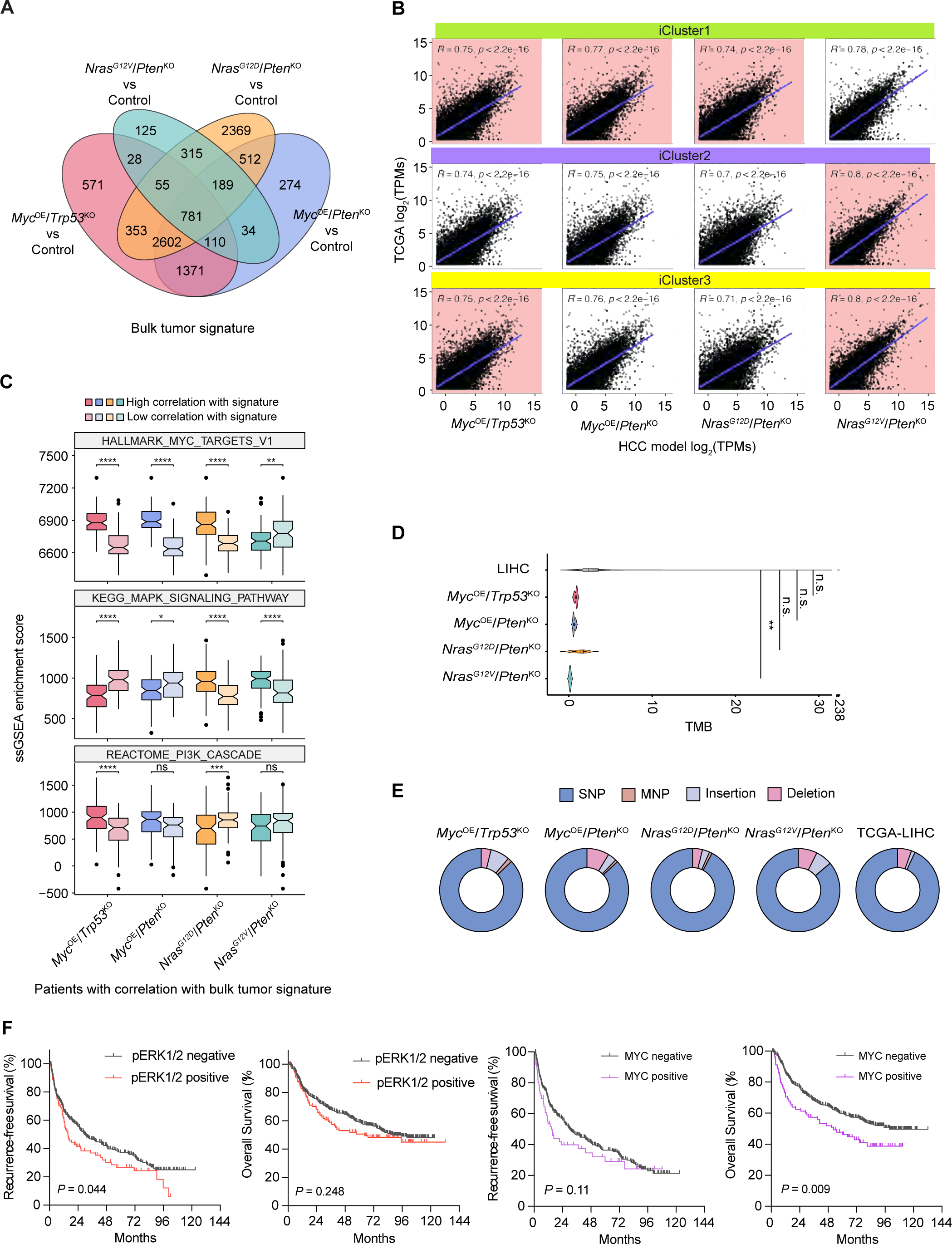
HCC patients with similar profiles to genetically-distinct murine models display distinct molecular and prognostic features. **A.** Venn diagram of the differentially expressed genes (DEG) between genetically-distinct HCC and control livers as determined in RNA-seq analyses (see **Supplementary Table 1**). **B.** Scatterplots depicting the correlation in global gene expression between TCGA-LIHC patients and each of the genetically-distinct HCC tumors. TCGA-LIHC patients were divided according to their TCGA classification^99^. **C.** Boxplots depicting the enrichment of the MYC, MAPK and PI3K/mTOR signaling pathways in patients segregated according to their high/low correlation with each of the genetically-distinct HCC bulk tumor transcriptional signatures from **Fig. 1E** (sample size n=93 for each set of patients). **D.** Violin plots depicting the tumor mutational burden (TMB) of genetically-distinct HCC compared to human LIHC (n= 723), as determined by whole exome sequencing analyses of bulk tumors (n=3 per genotype) (see **Supplementary Table 3**). **E.** Donut charts of the variant type distribution of mutations of genetically-distinct HCC (n=3 per genotype) compared to TCGA-LIHC (n= 371). SNP: single nucleotide polymorphism. MNP: multiple nucleotide polymorphism. **F.** Kaplan-Meier curves displaying the overall survival and recurrence-free survival of HCC patients (from the Wu *et al*. dataset^65^) segregated according to p-ERK1/2 and c-MYC positive or negative staining in cancer cells (see **Supplementary Table 4** for median OS and RFS time). Statistical significance was determined by unpaired Student’s T-test in (**C**), unpaired Wilcoxon test in (**D**), and log-rank test in (**F**). *p < 0.01, **p < 0.01, ****p < 0.0001; n.s. non-significant.

**Figure S3:**
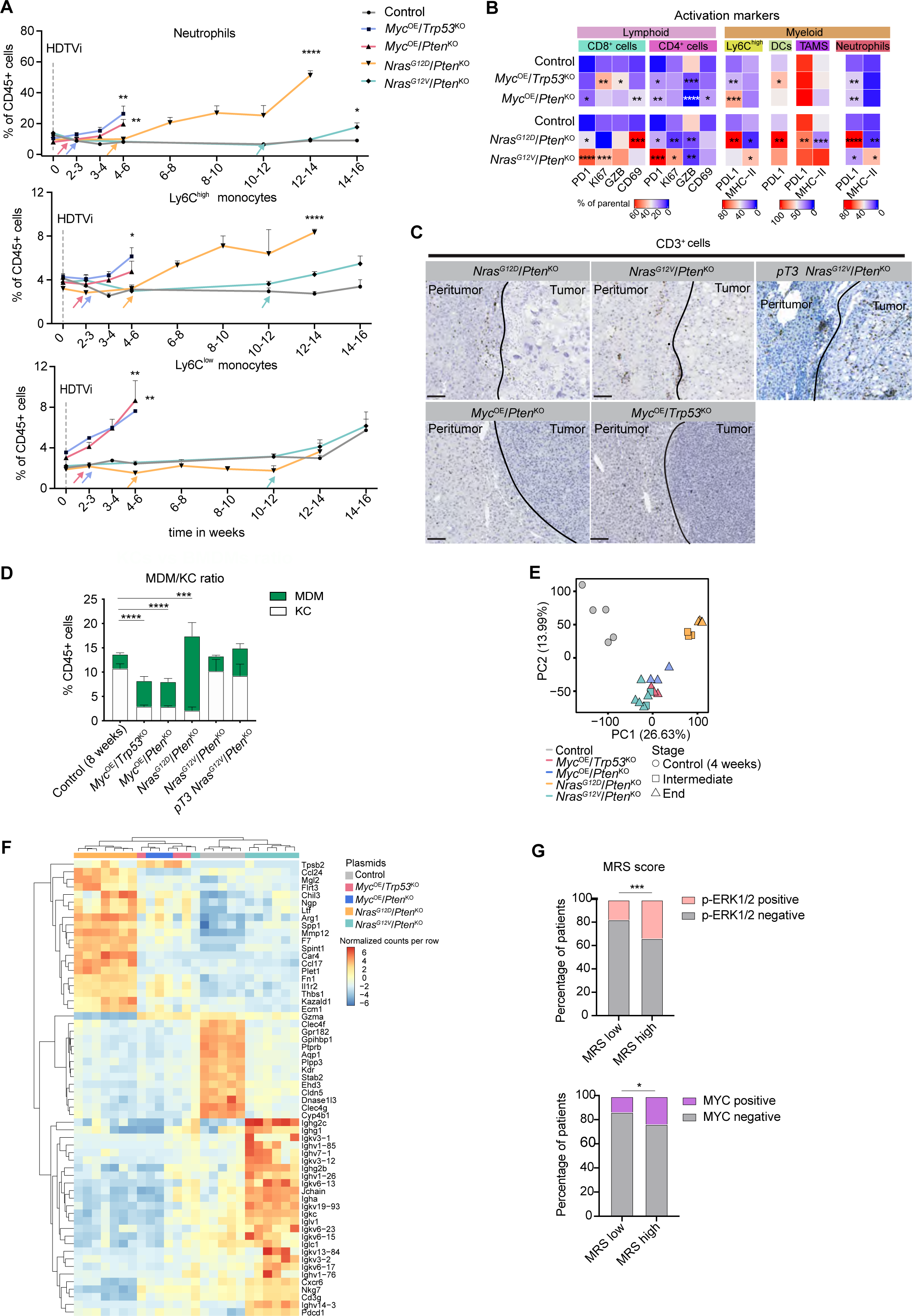
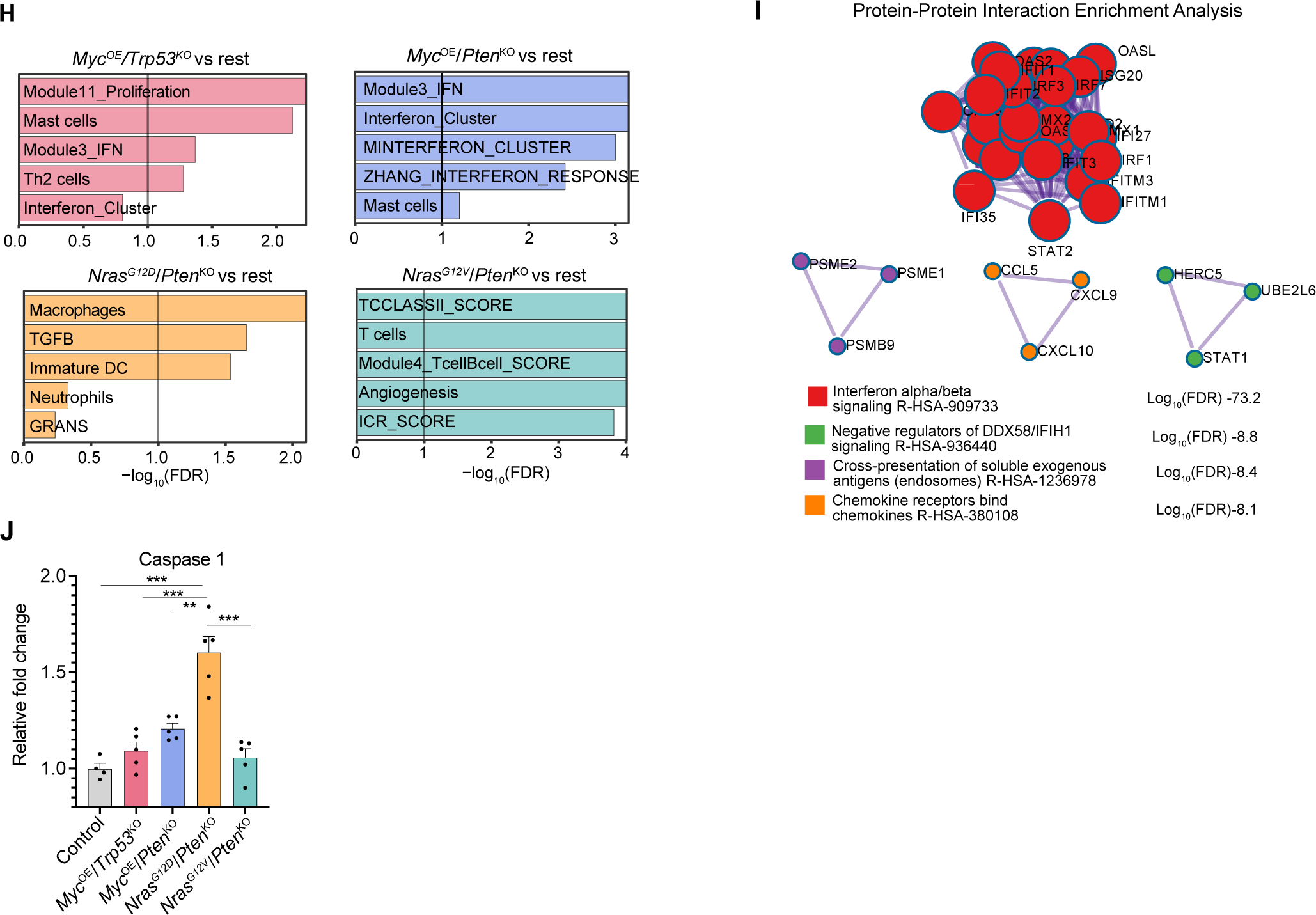
Dynamic content and transcriptional education of HCC immune landscape are shaped by cancer cell-intrinsic signaling pathway. **A.** Quantification of the longitudinal content of neutrophils, Ly6C^high^ monocytes and Ly6C^low^ monocytes (from the parental CD45^+^ CD11b^+^ cells) in blood collected at the indicated time points post-HDTVi (Control n=10, *Myc*^OE^/*Trp53*^KO^ n=20, *Myc*^OE^/*Pten*^KO^ n=19, *Nras*^G12D^/*Pten*^KO^ n=32, and *Nras^G12V^*/*Pten*^KO^ n=18). Arrows indicate the time points when detectable tumors were visible by MRI. **B.** Heatmap depicting the median percentage of the different immune cell populations in end-stage HCC (presented in **Fig. 3C**) expressing the indicated phenotypic markers. Statistical significance tested against aged-matched control livers. **C.** Representative CD3 IHC staining performed in liver sections from genetically-distinct HCC-bearing mice. Black lines indicate the tumor borders. Scale bars= 100 μm **D.** Barplots depicting the TAM content as percentages of F4/80^high^CD11b^low^ tissue-resident Kupffer cells (KCs) and F4/80^int^CD11b^high^ infiltrating MDMs (from the parental CD45^+^ cells). (Control n=7, *Myc*^OE^/*Trp53*^KO^ n= 6, *Myc*^OE^/*Pten*^KO^ n= 5, *Nras*^G12D^/*Pten*^KO^ n= 4, *Nras*^G12V^/*Pten*^KO^ n= 4, *pT3*-*Nras*^G12V^/*Pten*^KO^ n = 4). **E.** PCA plot depicting the CD45^+^ immune cell transcriptome of control livers and genetically-distinct HCCs at intermediate and end-stages following RNA-seq analyses (Control n=5, *Myc*^OE^/*Trp53*^KO^ n= 3, *Myc*^OE^/*Pten*^KO^ n= 3, *Nras*^G12D^/*Pten*^KO^ intermediate n= 3 and end-stage n=3, *Nras*^G12V^/*Pten*^KO^ intermediate n= 2 and end-stage n=5, see **Supplementary Table 5**). **F.** Heatmap of unsupervised hierarchical clustering depicting 60 genes with the highest variation between samples in the CD45^+^ immune cell transcriptome of control and genetically-distinct HCC models at intermediate and end-stages (presented in (**E**)). **G.** Barplots depicting the myeloid responsive score (MRS) in HCC patients from the Wu et al. dataset^65^ segregated according to p-ERK1/2 (top) and Myc (bottom) expressing cancer cells (see **Methods** for quantification). **H.** Barplots depicting the significant geneset enrichment for specific immune genesets (see **Methods**) in the transcriptome of CD45^+^ cells isolated from end-stage genetically-distinct HCC and compared to the rest of the HCC models (vs rest). Vertical line at - log10(FDR)=1 is used as threshold for significance. **I.** Network plots depicting enrichment analyses of the protein-protein interactions from the interferon response genes identified in the *Myc^OE^* HCC models (presented in (**H**)). Node size represents the -log10 (FDR). **J.** Barplots depicting caspase 1 activity in control and end-stage tumors from genetically-distinct HCCs. Results are shown as relative fold change compared to control. (Control n=4, *Myc*^OE^/*Trp53*^KO^ n= 5, *Myc*^OE^/*Pten*^KO^ n= 5, *Nras*^G12D^/*Pten*^KO^ n= 5, *Nras*^G12V^/*Pten*^KO^ n= 5). Graphs show mean + SEM (**A**, **D** and **J**). Statistical significance was determined by two-sided unpaired Student’s *t*-test analysis in (**A, D, J),** *χ*² test in (**G**) and one-way ANOVA with Tukey’s multiple comparison test (**B**). Significance was determined for the KC content (**D**). *p < 0.05; **p < 0.01; ***p < 0.001; ****p < 0.0001.

**Figure S4:**
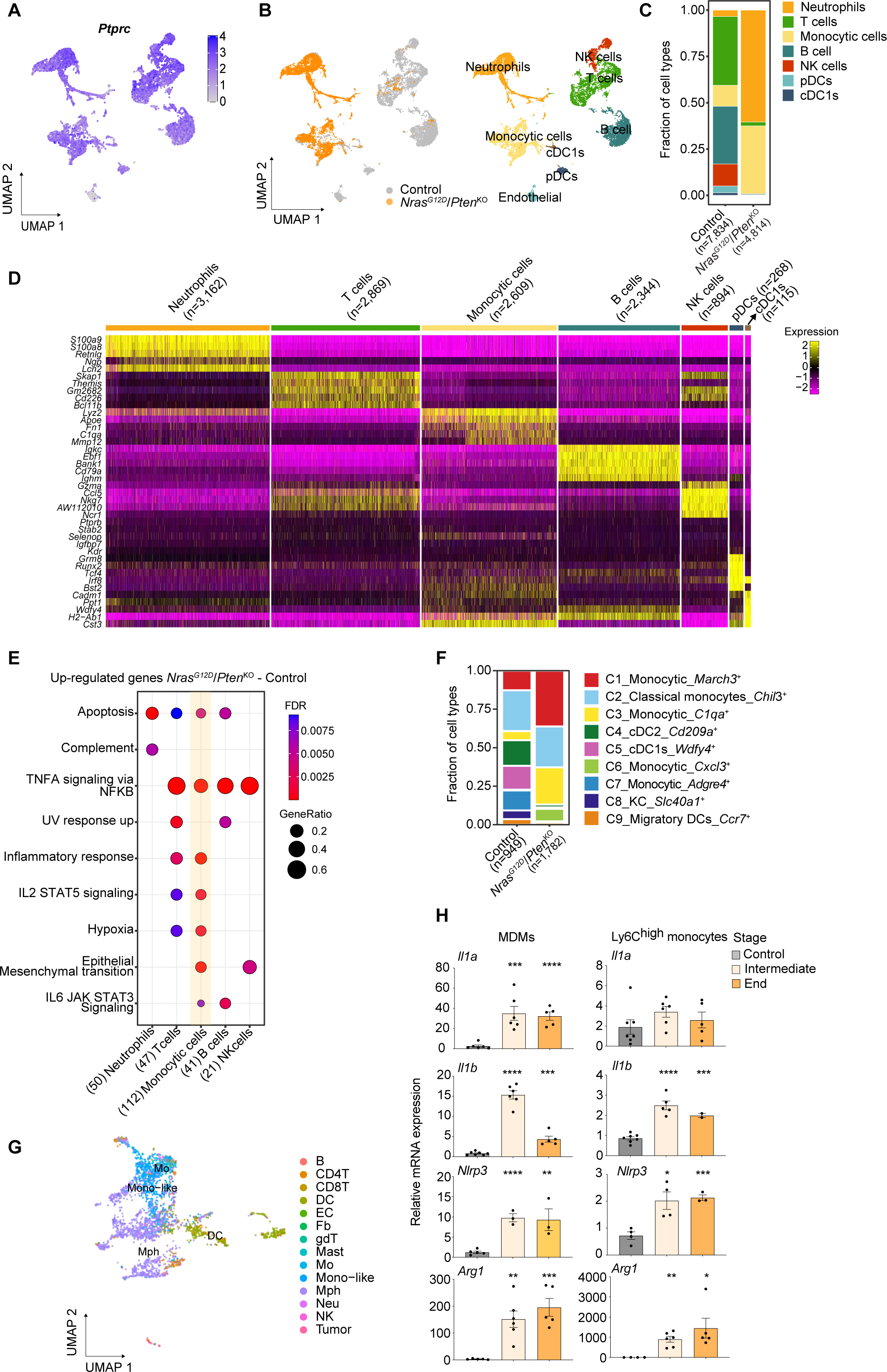
In-depth analysis of *Nras*^G12D^/*Pten*^KO^ tumor immune landscape. **A.** *Ptprc* (CD45) gene expression levels as identified in the scRNA-seq dataset of control liver and *Nras*^G12D^/*Pten*^KO^ HCC shown in **Fig. 4A**. **B.** UMAP representation as shown in **Fig. 4A** including the ‘Endothelial cells’ cluster. **C.** Barplots depicting the proportions of the indicated immune cell subsets as identified in the scRNA-seq dataset of control liver and *Nras*^G12D^/*Pten*^KO^ HCC. **D.** Heatmap displaying the expression levels of a selection of the top most differentially expressed genes specific to each of the indicated immune cell subsets and their expression across the other immune clusters from the scRNA-seq dataset in **Fig. 4A** (see **Supplementary Table 7**). **E.** Bubble plot showing the gene set enrichment of up-regulated genes in *Nras*^G12D^/*Pten*^KO^ HCC compared to control liver in the indicated immune cell subsets. Circle size represents the ratio of overlapping genes between the gene set and the DEG. Significance threshold FDR<0.05 (see **Supplementary Table 8**). **F.** Barplots depicting the proportions of the indicated myeloid cell subsets as identified in the ‘Monocytic cell’ cluster in control liver and *Nras*^G12D^/*Pten*^KO^ HCC scRNA-seq dataset (presented in **Fig 4D**). **G.** UMAP of the subclusters from the ‘Monocytic cell’ population in control liver and *Nras*^G12D^/*Pten*^KO^ HCC scRNA-seq dataset (presented in **Fig 4D**) displaying the annotations from human HCC scRNA-seq dataset^71^ subclusters after integration of both datasets (see **Methods**). **H.** Barplots showing the relative mRNA expression levels of indicated genes in FACS-isolated MDMs and Ly6C^high^ monocytes from control liver and *Nras*^G12D^/*Pten*^KO^ HCC at intermediate and end-stages. *Ubc* was used as a housekeeping gene in all analyses. All samples are relative to one control liver sample. Graphs show mean ± SEM (**H**). Statistical significance was determined by two-sided unpaired Student’s *t*-test (**H**). *p < 0.05; ** p < 0.01; *** p < 0.001; **** p < 0.0001.

**Figure S5.**
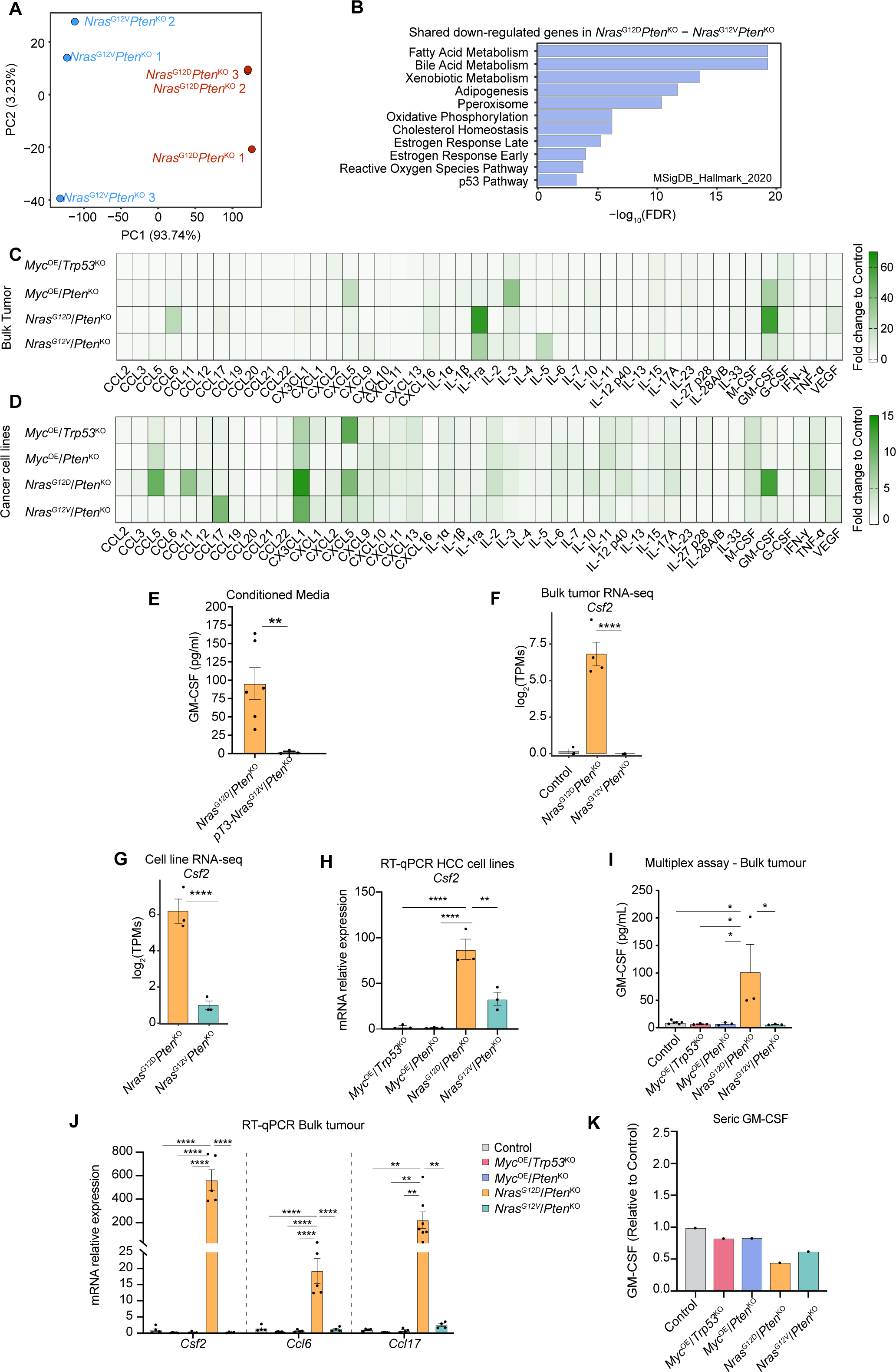
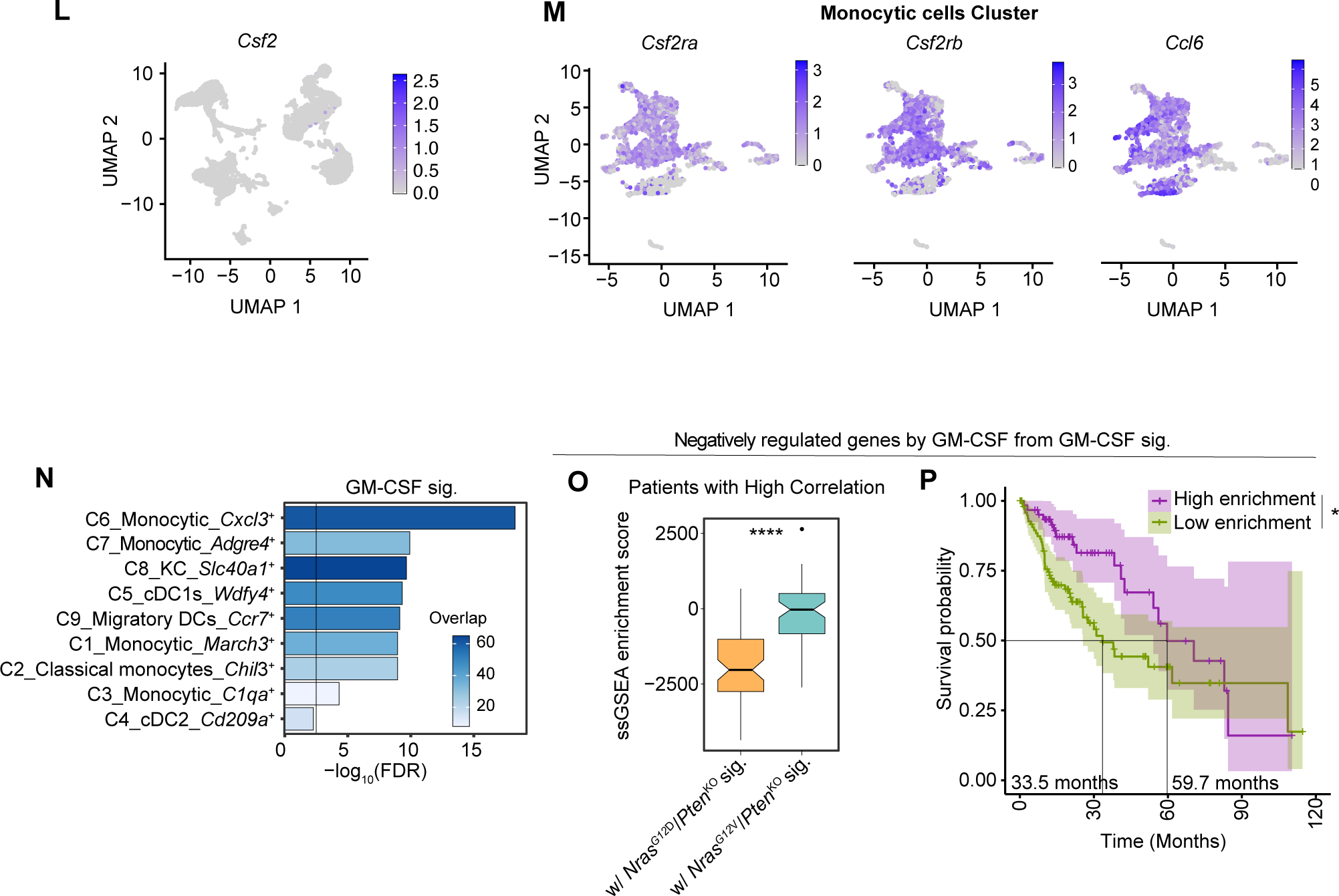
The GM-CSF signaling pathway underlies the reciprocal interaction between *Nras*^G12D^/*Pten*^KO^ cancer cells and their TME. **A.** PCA plot depicting the *Nras*^G12D^/*Pten*^KO^ and *Nras*^G12V^/*Pten*^KO^ HCC cell line transcriptome (n=3 per cell line). **B.** Barplot depicting the signaling pathways enriched in the down-regulated genes shown in **Fig. 5A** (blue colored dots). **C-D.** Proteomic profile analyses of *Myc^OE^/Trp53^KO^*, *Myc^OE^/Pten^KO^*, *Nras*^G12D^/*Pten*^KO^ and *Nras*^G12V^/*Pten*^KO^ bulk tumors (**C**) or conditioned media collected from HCC cancer cell lines (**D**), relative to control livers or AML12 cell line, respectively. Indicated values are shown as fold change of the pixel density compared to control for each assessed protein. **E.** Barplot depicting the quantification of GM-CSF secretion in the conditioned media of *Nras*^G12D^/*Pten*^KO^ (n=6, from **Fig. 5D**) and *pT3-Nras*^G12V^/*Pten*^KO^ cell lines (n=3). **F-G.** Barplot depicting the *Csf2* gene expression levels extracted from RNA-seq datasets of control livers (n=3), bulk *Nras*^G12D^/*Pten*^KO^ (n=4) and *Nras*^G12V^/*Pten*^KO^ tumors (n=3) (**F**) or *Nras*^G12D^/*Pten*^KO^ (n=3) and *Nras*^G12V^/*Pten*^KO^ HCC cell lines (**G**) (n=3 per cell line). H. Barplot depicting the *Csf2* gene expression levels assessed by RT-qPCR in *Myc^OE^/Trp53^KO^*, *Myc^OE^/Pten^KO^*, *Nras*^G12D^/*Pten*^KO^ and *Nras*^G12V^/*Pten*^KO^ HCC cell lines (n=3 for each cell line). One *Myc^OE^/Trp53^KO^* sample is set as baseline level. **I.** Barplot depicting the GM-CSF protein levels assessed by Luminex assay in control livers (n=6), *Myc^OE^/Trp53^KO^*(n=3), *Myc^OE^/Pten^KO^* (n=3), *Nras*^G12D^/*Pten*^KO^ (n=3) and *Nras*^G12V^/*Pten*^KO^ (n=3) bulk tumor lysates. **J.** Barplots depicting the *Csf2*, *Ccl6* and *Ccl17* gene expression levels assessed by RT-qPCR in control (n=4), *Myc^OE^/Trp53^KO^* (n=5), *Myc^OE^/Pten^KO^* (n=5), *Nras*^G12D^/*Pten*^KO^ (n=5) and *Nras*^G12V^/*Pten*^KO^(n=4) end-stage bulk tumors. One control liver sample is set as the baseline level for all genes. **K.** Barplot depicting the relative pixel density of GM-CSF levels determined by proteomic profile analyses of control mouse and genetically-distinct HCC-bearing mice sera. Control sample is set as the baseline level. **L.** UMAP representation of *Csf2* expression in the scRNA-seq from **Fig. 4A**. **M.** UMAP representation of *Csf2ra, Csf2rb* and *Ccl6* expression in the ‘Monocytic cell’ clusters identified by scRNA-seq (**Fig. 4D**). N. Barplot depicting the enrichment of the GM-CSF signature in each of the indicated subpopulations of the ‘Monocytic cells’ cluster identified by scRNA-seq (presented in **Fig. 4D**). **O.** Boxplot depicting the enrichment of GM-CSF negatively regulated genes (n=709) from the GM-CSF signature in TCGA: LIHC patients segregated according to their high correlation with the *Nras*^G12D^/*Pten*^KO^ or *Nras*^G12V^/*Pten*^KO^ transcriptional signatures (same sample size as **Fig. 5I**). **P.** Kaplan-Meier survival curves of TCGA: LIHC patients segregated according to their high/low enrichment of GM-CSF negatively regulated genes (n=709) from the GM-CSF signature (High n=62, Low n=103). Graphs show mean ± SEM (**E-J**). Statistical significance was determined by differential expression analyses (**F, G**) one-way ANOVA (**H-J**), unpaired Student’s T-test (**E, O**) and log-rank test (**P**). Vertical lines at -log10(FDR)=2.5 are used as threshold for (**B, N**). *p < 0.05; **p < 0.01; ****p < 0.0001.

**Figure S6:**
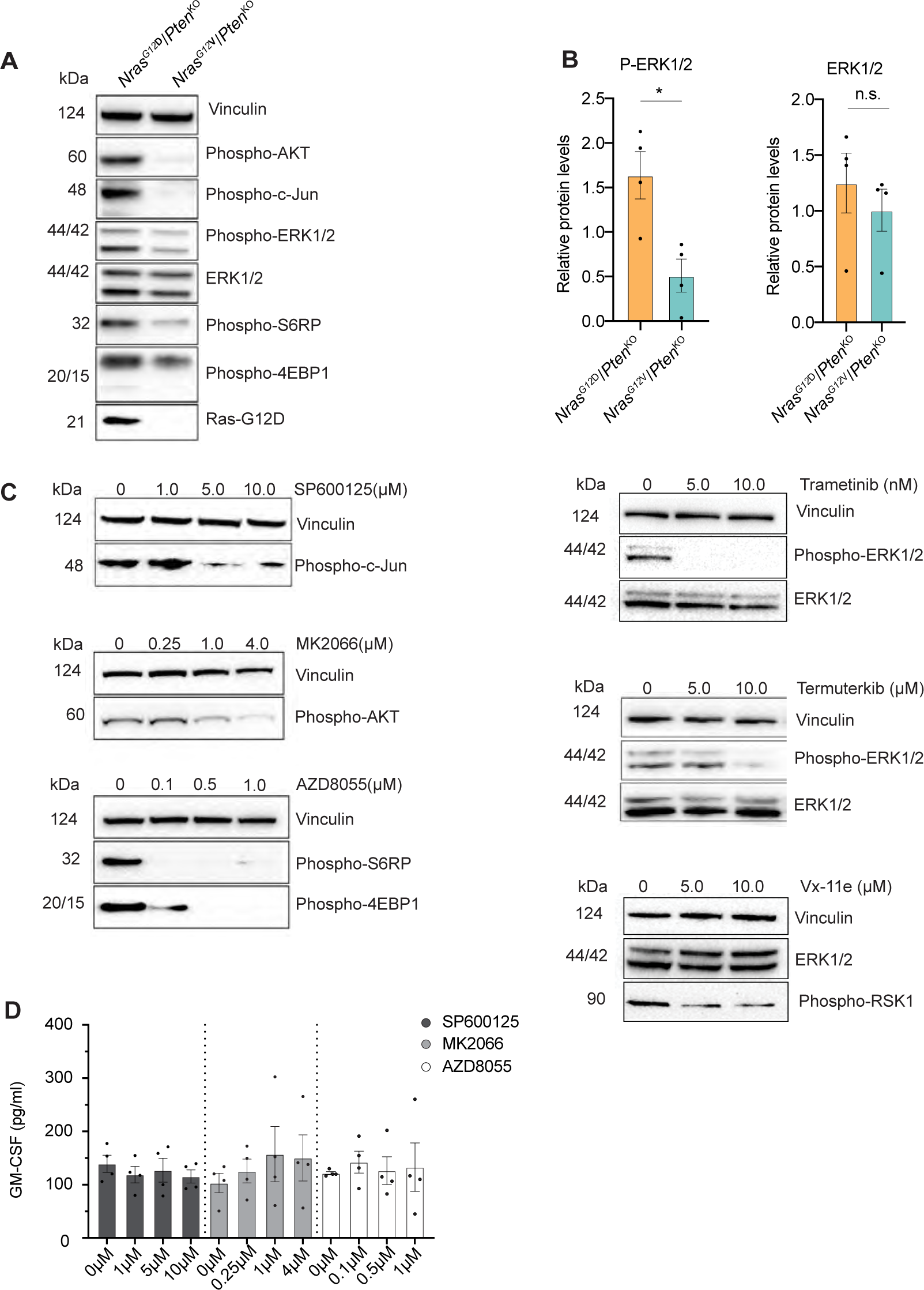
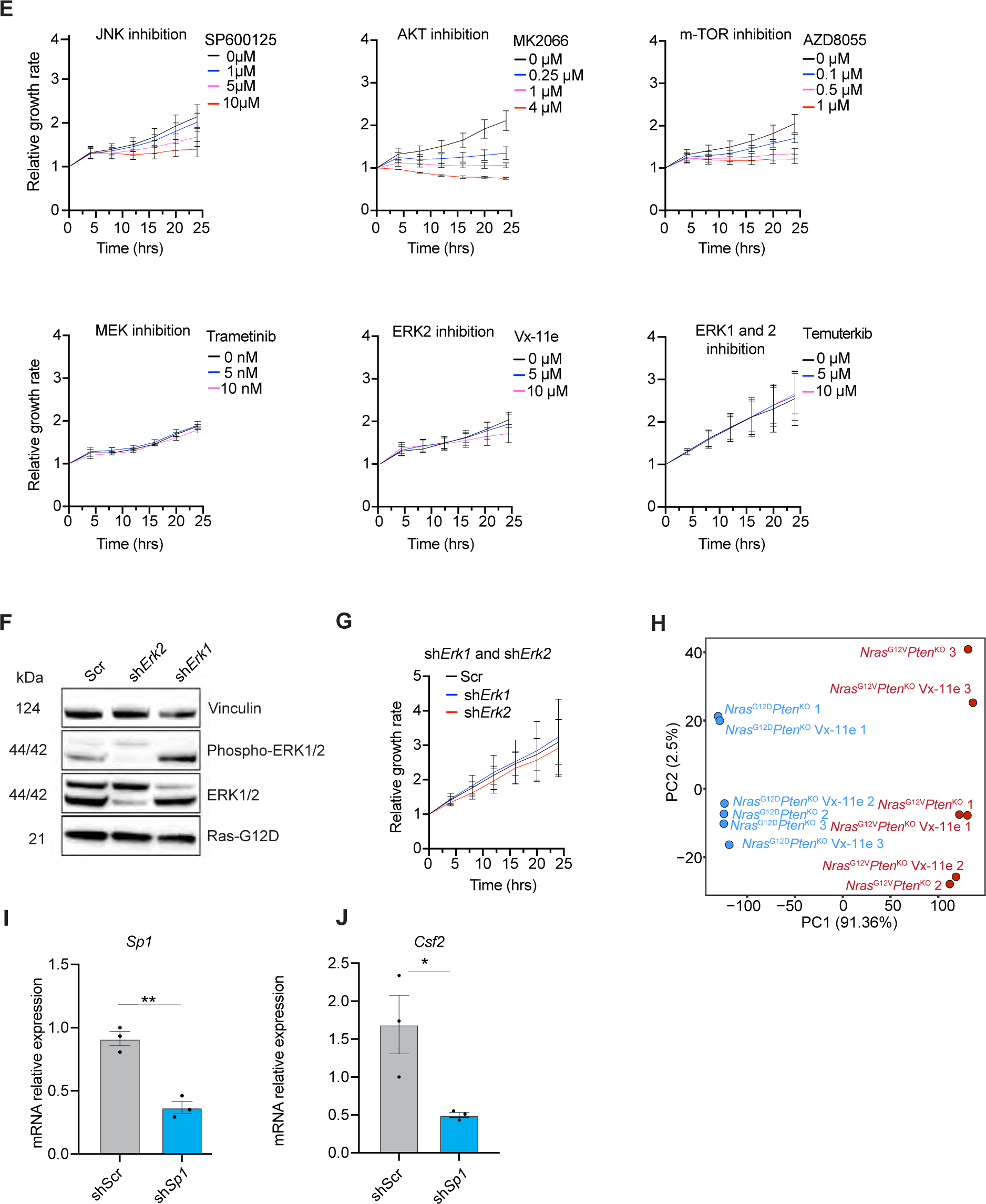
*Nras*^G12D^-induced MAPK-ERK1/2 signaling pathway specifically modulates GM-CSF levels via the transcription factor SP1. **A.** Representative western blots assessing the expression of the indicated proteins in *Nras*^G12D^/*Pten*^KO^ and *Nras*^G12V^/*Pten*^KO^ HCC cell lines. **B.** Barplots depicting the expression levels of phosphorylated and total ERK1/ERK2 proteins relative to vinculin (presented in **A**, n=4 per cell line). **C.** Representative western blots assessing the expression of the indicated proteins in *Nras*^G12D^/*Pten*^KO^ HCC cell lines after 24h of treatment with SP600125 (JNK inhibitor), MK2066 (AKT inhibitor), AZD8055 (mTOR inhibitor), Trametinib (MEK1/2 inhibitor), Temuterkib (ERK1/2 inhibitor) and VX-11e (ERK2 inhibitor) at the indicated concentrations. **D.** Barplots depicting the GM-CSF protein levels quantified in *Nras*^G12D^/*Pten*^KO^ HCC cell line conditioned media after 24h of treatment with SP600125 (JNK inhibitor; n=4), MK2066 (AKT inhibitor; n=4) and AZD8055 (m-TOR inhibitor; n=4) at the indicated drug concentrations. **E.** Growth curves showing the proliferation of *Nras*^G12D^/*Pten*^KO^ cancer cells treated with Trametinib (MEK1/2 inhibitor), Temuterkib (ERK1/2 inhibitor), Vx-11e (ERK2 inhibitor) SP600125 (JNK inhibitor), MK2066 (AKT inhibitor) and AZD8055 (mTOR inhibitor) at the indicated concentrations (n=3 independent experiments). Growth rate is relative to the 0h timepoint. **F.** Representative western blots assessing the expression of the indicated proteins in *Nras*^G12D^/*Pten*^KO^ HCC cell line expressing either a non-targeting control shRNA (sh*Scr*), an shRNA directed against *Erk1* (sh*Erk1*), or *Erk2* (sh*Erk2*). **G.** Growth curves showing the proliferation of sh*Scr*, sh*Erk1* and s*hErk2 Nras*^G12D^/*Pten*^KO^ cancer cell lines (n=3 independent experiments). Growth rate is relative to the 0h timepoint. **H.** PCA plot depicting the transcriptome of genetically-distinct HCC cell lines (*Nras*^G12D^/*Pten*^KO^, *Nras^G12V^*/*Pten*^KO^) treated with Vx-11e and control (each sample n=3), following RNA-seq analyses (see **Supplementary Table 9**). **I-J.** Gene expression analysis of *Sp1* (**I**) and *Csf2* (**J**) performed in sh*Scr* (n=-3) or sh*Sp1* (n=3) *Nras*^G12D^/*Pten*^KO^ cells. *Ubc* was used as a house keeping gene. All samples are relative to one sh*Scr* sample. Graph shows mean ± SEM (**B, D, E, G, I-J**). Statistical significance was determined by Student t-test (**B, I-J).** * p < 0.05, ** p < 0.01; n.s. non-significant.

**Figure S7:**
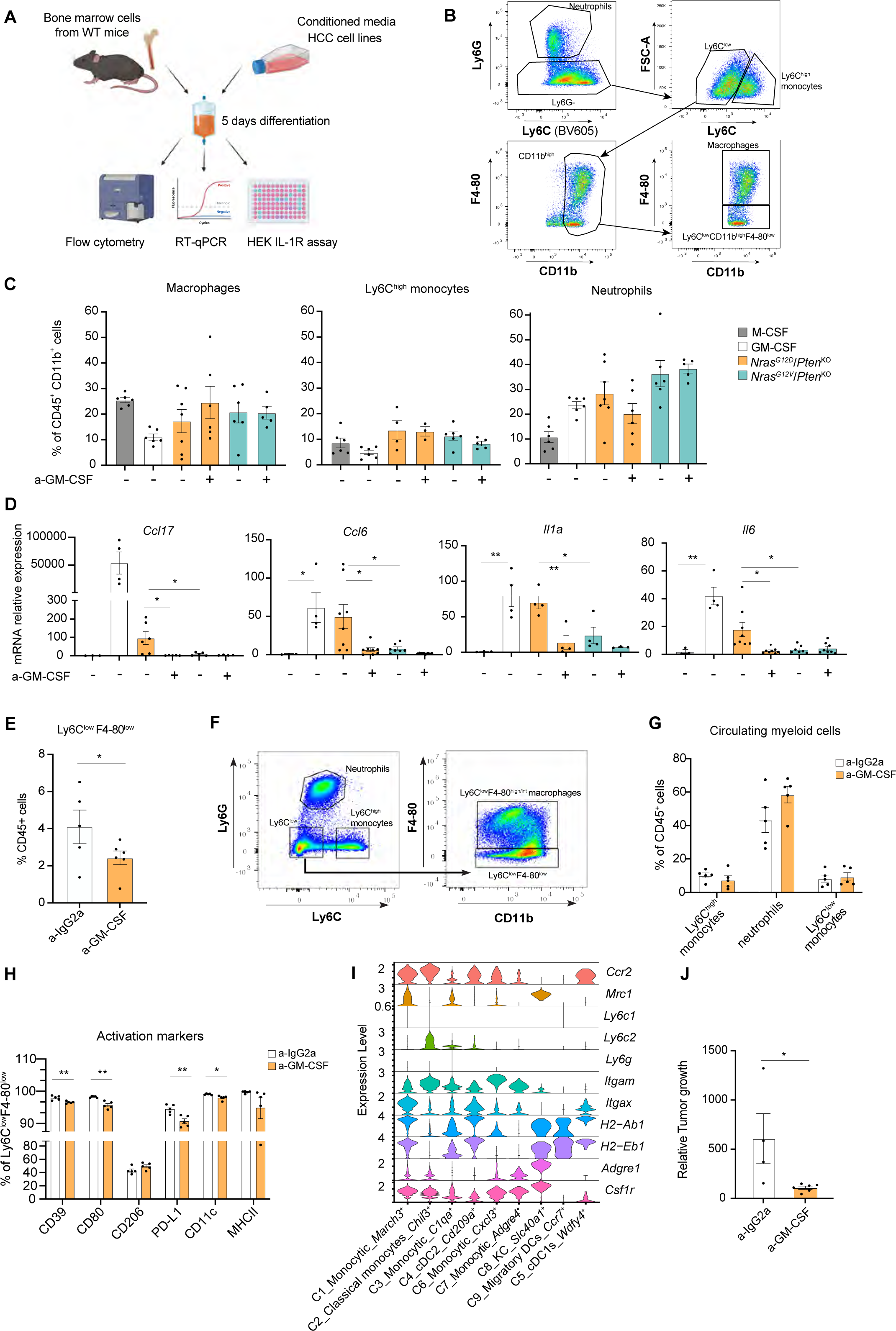
GM-CSF inhibition hinders the differentiation of inflammatory monocyte-derived Ly6C^low^ myeloid cells *in vitro* and *in vivo*. **A.** Experimental design depicting the pipeline to differentiate bone marrow (BM) cells in distinct conditioned media (CM) from HCC cell lines and follow-up procedures applied. BM cells are exposed to HCC cell-derived CM (generated in 0% media for 24h) for 5 days with 10% FCS supplementation to allow differentiation. BM cells differentiated in recombinant M-CSF or GM-CSF in DMEM 10% FCS were used as controls. Differentiated BM cells were then used in flow cytometry assays, RT-qPCR analyses, and CM assessment of IL-1R signaling activity using HEK IL-1R reporter cells (see **Methods**). **B.** Flow cytometry gating strategy used to determine the relative abundance of distinct myeloid cells (neutrophils, Ly6C^high^ monocytes, macrophages and Ly6C^low^CD11b^high^ F4/80^low^cells) generated in BM differentiation experiments. The first gate is based on Ly6C and Ly6G expression and is set on the total myeloid cell population (CD45^+^ CD11b^+^). Sample shown here represents one replicate of BM cells differentiated in *Nras*^G12D^/*Pten*^KO^ CM. **C.** Barplots depicting the percentage of macrophages (CD45^+^ CD11b^+^ Ly6G^-^ Ly6C^low^ F4/80^+^), neutrophils (CD45^+^ CD11b^+^ Ly6G^inter^ Ly6G^high^) and Ly6C^high^ monocytes (CD45^+^ CD11b^+^ Ly6G^-^ Ly6C^high^) relative to CD45^+^CD11b^+^ total myeloid cells obtained from BM cells differentiated in either recombinant M-CSF or GM-CSF, or in CM prepared from distinct HCC cell lines, with or without GM-CSF neutralizing antibody (a-GM-CSF) (M-CSF n=6, GM-CSF n=6, *Nras*^G12D^/*Pten*^KO^ n=7, *Nras*^G12D^/*Pten*^KO^ + a- GM-CSF n=6, *Nras*^G12V^/*Pten*^KO^ n=6, *Nras*^G12V^/*Pten*^KO^ + a-GM-CSF n=5). **D.** Barplots depicting the relative mRNA expression of indicated genes in BM cells differentiated in either recombinant M-CSF (n=3), GM-CSF (n=4), or in the CM of *Nras*^G12D^/*Pten*^KO^ and *Nras*^G12V^/*Pten*^KO^ HCC cell lines (n = 4-8), with or without a-GM-CSF (n = 4-8). Values are represented as a relative fold change compared to one M-CSF BM sample. *Ubc* was used as a housekeeping gene. **E.** Barplot depicting the abundance of Ly6C^low^F4/80^low^ cells from HDTVi-induced *Nras*^G12D^/*Pten*^KO^ HCC-bearing mice at end-stage treated with a-IgG2a (n=5) or a-GM-CSF (n=6). **F.** Flow cytometry gating strategy used to determine the relative abundance of Ly6C^low^ F4/80^low^ from HCC samples. Myeloid cells were gated according to **Supplementary Data 1**. **G.** Barplot depicting the abundance of Ly6C^high^ monocytes, neutrophils and Ly6C^low^ monocytes in the blood collected from HDTVi-induced *Nras*^G12D^/*Pten*^KO^ HCC-bearing mice 2 weeks post-treatment with a-IgG2a (n=5) or a-GM-CSF (n=5). **H.** Barplots depicting the percentage of Ly6C^low^F4-80^low^ cells from HDTVi-induced *Nras*^G12D^/*Pten*^KO^ HCC-bearing mice sacrificed at 2 weeks post treatment with a-IgG2a (n=5) or a-GM-CSF (n=5) (shown in **Fig. 7E**) expressing the indicated phenotypic markers. **I.** Violin plots depicting the expression levels of selected genes in the indicated myeloid cell subsets in control livers and *Nras*^G12D^/*Pten*^KO^ HCC from the ‘Monocytic cell’ subpopulation. **J.** Barplot depicting the relative tumor growth of HDTVi-induced *Nras*^G12D^/*Pten*^KO^ mice treated with either a-IgG2a (n=4) or a-GM-CSF (n=6) at 4 weeks post-treatment initiation. Values are relative to tumor volumes at treatment start. Graph shows mean ± SEM (**C**-**E, G, H, J**). Statistical significance was determined by unpaired Student’s T-test in (**C-E, H, J**). *p < 0.05; **p < 0.01.

**Figure S8:**
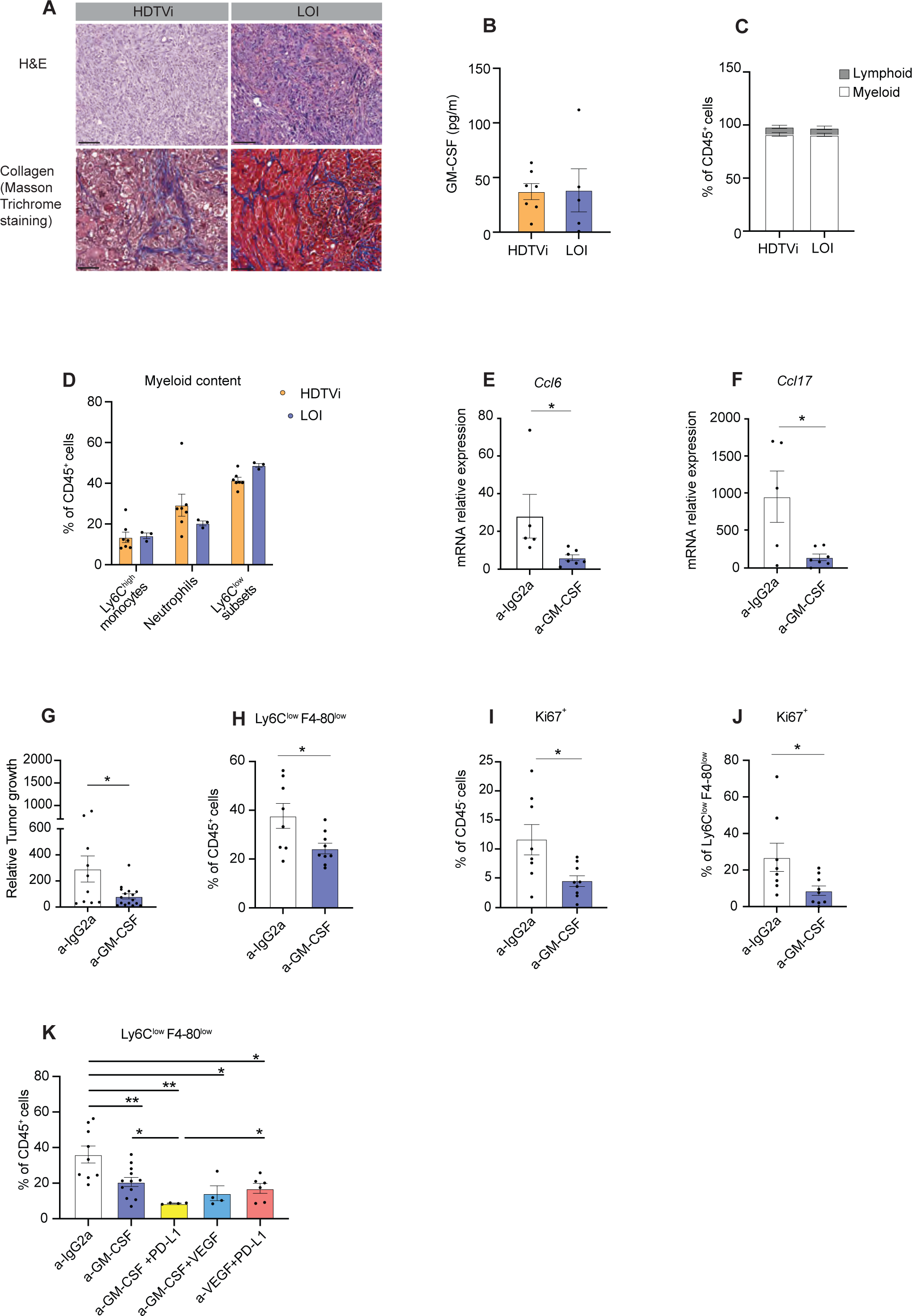
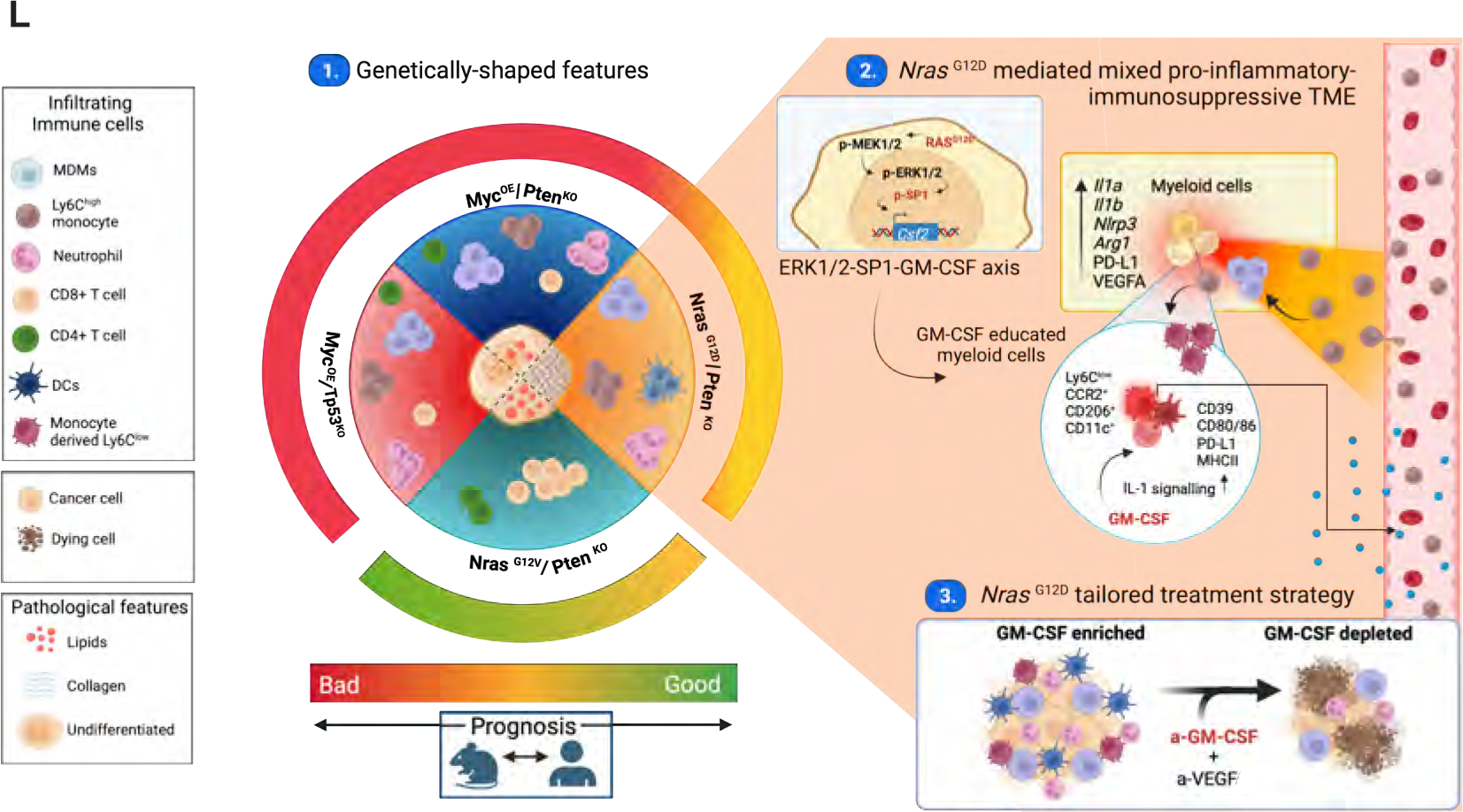
*In vivo* GM-CSF and VEGF blockade abrogate the accumulation of pro-tumorigenic Ly6C^low^ myeloid cells in *Nras*^G12D^/*Pten*^KO^-driven HCC. **A.** Representative IHC images of H&E and Masson Trichrome staining performed on liver sections from HDTVi- and LOI-induced *Nras*^G12D^/*Pten*^KO^ HCC-bearing mice at end-stage. **B.** Barplot depicting the quantification of intratumoral GM-CSF levels present in end-stage HDTVi- (n=7) and LOI-induced (n=4) *Nras*^G12D^/*Pten*^KO^ HCC. **C.** Quantification of the content of myeloid (CD45^+^ CD11b^+^) and lymphoid (CD45^+^ CD11b^-^) cells relative to total CD45^+^ leukocytes in tumors collected from end-stage HDTVi- (n=7) and LOI-induced (n=5) *Nras*^G12D^/*Pten*^KO^ HCC. HDTVi-induced end-stage *Nras*^G12D^/*Pten*^KO^ HCC-bearing mice shown in **Fig. 3A** are included in this graph. **D.** Barplots depicting the percentage of intratumoral Ly6C^high^ monocytes, neutrophils and Ly6C^low^ subsets relative to total CD45^+^ leukocytes in end-stage HDTVi- (n=7) and LOI-induced (n=3) *Nras*^G12D^/*Pten*^KO^ HCC. **E-F.** Barplot depicting the relative mRNA expression of *Ccl6* (**E**) and *Ccl17* (**F**) genes in end-stage LOI-induced *Nras*^G12D^/*Pten*^KO^ HCC post-preclinical trial treatment with a- IgG2a (n=5) or a-GM-CSF (n=7). Values represent relative fold changes compared to one IgG2a-treated sample. *Ubc* was used as a housekeeping gene. **G.** Barplot depicting the relative tumor growth of LOI-induced *Nras*^G12D^/*Pten*^KO^ HCC at 2 weeks post-treatment initiation with either a-IgG2a (n = 10) or a-GM-CSF (n = 16). **H.** Barplot depicting the percentage of intratumoral Ly6C^low^F4/80^low^ cells relative to total CD45^+^ leukocytes in LOI-induced *Nras*^G12D^/*Pten*^KO^ HCC post-preclinical trial treatment with a-IgG2a (n = 8) or a-GM-CSF (n = 9). **I.** Barplot depicting the percentage of CD45^-^Ki67^+^ cells in LOI-induced *Nras*^G12D^/*Pten*^KO^ HCC post-preclinical trial treatment with a-IgG2a (n = 8) or a-GM-CSF (n = 9). **J.** Barplot depicting the percentage of Ki67^+^Ly6C^low^F4/80^low^ cells in LOI-induced *Nras*^G12D^/*Pten*^KO^ HCC post-preclinical trial treatment with a-IgG2a (n = 8) or a-GM-CSF (n = 9). **K.** Barplot depicting the percentage of intratumoral Ly6C^low^F4/80^low^ cells relative to total CD45^+^ leukocytes in LOI-induced *Nras*^G12D^/*Pten*^KO^ HCC post-preclinical trial treatment with a-IgG2a (n = 9), a-GM-CSF (n = 12), a-GM-CSF+a-PD-L1 (n =4), a- GM-CSF- + a-VEGF (4), and a-VEGF + a-PD-L1 (n = 6). LOI-induced *Nras*^G12D^/*Pten*^KO^ HCC-bearing mice treated with a-IgG2a (n=8) and a-GM-CSF (n=9) shown in **Fig. S8H** are included in this graph. **L. 1.** Cancer cell genetics shape distinct histopathological features and TME contexture in genetically-distinct murine HCC, which overall predict the prognostic rates of distinct human HCC subclasses **2.** *Nras*^G12D^/*Pten*^KO^ secretome fosters a myeloid dominant, pro-inflammatory/immunosuppressive TME wherein *Nras^G12D^* driven, ERK1/2-SP1-dependent GM-CSF expression drives the accumulation of pro-tumoral Ly6C^low^ monocyte-derived cells. **3.** Neutralizing GM-CSF and VEGF in *Nras^G12D^/Pten^KO^* HCC reprograms the TME, promotes cancer cell death and significantly extends animal survival. Graph shows mean ± SEM (**B, D-K**). Statistical significance was determined by unpaired Student’s T (**E- K**). *p < 0.05; **p<0.01; ***p < 0.001.

